# Thirty clues to the exceptional diversification of flowering plants

**DOI:** 10.1101/279620

**Authors:** Susana Magallón, Luna L. Sánchez-Reyes, Sandra L. Gómez-Acevedo

**Author notes:** Author for correspondence: Susana Magallón.

## Abstract

- As angiosperms became one of the megadiverse groups of macroscopic eukaryotes, they forged modern ecosystems and promoted the evolution of extant terrestrial biota. Unequal distribution of species among lineages suggests that diversification, the process which ultimately determines species-richness, acted differentially through angiosperm evolution.
- We investigate how angiosperms became megadiverse by identifying the phylogenetic and temporal placement of exceptional radiations, by combining the most densely fossil-calibrated molecular clock phylogeny with a Bayesian model that identifies diversification shifts among evolutionary lineages and through time. We evaluate the effect of the prior number of expected shifts in the phylogenetic tree.
- Major diversification increases took place over 100 Ma, from the Early Cretaceous to the end Paleogene, and are distributed across the angiosperm phylogeny. Angiosperm long-term diversification trajectory shows moderate rate variation, but is underlain by increasing speciation and extinction, and results from temporally overlapping, independent radiations and depletions in component lineages.
- The identified deep time diversification shifts are clues to identify ultimate drivers of angiosperm megadiversity, which probably involve multivariate interactions among intrinsic traits and extrinsic forces. An enhanced understanding of angiosperm diversification will involve a more precise phylogenetic location of diversification shifts, and integration of fossil information.

## Introduction

Flowering plants (Angiospermae) represent the most recent evolutionary explosion of embryophytes, a lineage that occupied land at least 470 million years ago (Ma) (Rubinstein *et al*., 2010), and diverged from their closest living relatives 300-350 Ma (Magallón *et al*., 2013). Since their first appearance in the fossil record, angiosperms have radiated exceptionally, surpassing all other plant lineages not only in sheer species-richness, but to become ecologically predominant forming the structural and energetic basis of nearly all extant terrestrial biomes. Through their ecological expansion, angiosperms promoted the diversification of other plants (Schneider *et al*., 2004, Laenen *et al*., 2014), animals (Cardinal & Danforth, 2013; Wang *et al*., 2013), fungi (Guzmán *et al*., 2013, Kraichak *et al*., 2015), and bacteria (Goffredi *et al*., 2011). Human nutrition, culture, and well-being inextricably depend on angiosperms.

With between 295,000 and 304,000 described species (Christenhusz & Byng, 2016; The Plant List, 2017) and an estimated total of >350,000 species (Joppa *et al*., 2010), angiosperms are among the megadiverse groups of macroscopic eukaryotes. Their exceptional diversity is distributed unequally among evolutionary lines, some of which include tens-of-thousands of species (e.g., orchids, composites) and others fewer than ten (e.g., lotus, London planes), indicating that the process of diversification, the balance between speciation and extinction which ultimately determines species-richness (subsequently diversity), has acted differentially through angiosperm evolution.

Many studies have attempted to identify the factors that underlie angiosperm exceptional diversity, including intrinsic attributes (Farrell *et al*., 1991; Sargent, 2004), ecological interactions (Weber & Agrawal, 2014; van Der Niet & Johnson, 2012), extrinsic opportunity (Hughes & Atchinson, 2015; Moore & Donoghue, 2007), or complex interactions among them (Vamosi & Vamosi, 2011; Spriggs *et al*., 2015). Nevertheless, little is known about the dynamics of the diversification process that underlies the acquisition of angiosperm megadiveristy through time, and its unequal distribution among phylogenetic branches. A long tradition considered that the myriad of angiosperm unique vegetative and reproductive attributes made them competitively superior (Stebbins, 1974). Studies based on current phylogenetic understanding have shown that it is unlikely that increased phylogenetic branching characterizes angiosperms as a whole (Sanderson & Donoghue, 1994). Groups with unexpectedly high or low diversity, given a time-homogeneous diversification rate, have been identified (Magallón & Sanderson, 2001); and diversification tests based on tree asymmetry or model selection have found significant imbalance in net diversification rates among angiosperm lineages (Davies *et al*., 2004); that diversification shifts do not always correspond to named taxonomic entities (Smith *et al*., 2011); and that some occurred downstream major genomic duplications (Tank *et al*., 2015).

The fossil record unequivocally documents an Early Cretaceous crown angiosperm explosive radiation shortly following the appearance between the Valanginian and the Hauterivian (140-130 Ma) of pollen grains with a combination of detailed microstructural attributes found only among some angiosperm lineages (Hughes & McDougall, 1987; Brenner, 1996). Immediately younger sediments contain an explosively increasing diversity of pollen types and vegetative and reproductive remains representing early angiosperm branches (Friis *et al*., 2004, 2009) and their major evolutionary lineages (Doyle *et al*., 1977; Eklund *et al*., 2004; Mohr & Bernardes-de-Oliveira, 2004; Herendeen *et al*., 2017).

In this study we investigate the dynamics of angiosperm macroevolutionary diversification to investigate the phylogenetic placement of major diversification shifts, if there are phylogenetic regions and times in which radiations are concentrated, if there have been diversification decreases through angiosperm evolution, and, by examining their long term diversification trajectory, if angiosperms are in decline. We use a molecular Bayesian phylogenetic tree in which ca. 90% of angiosperm families are represented. This tree was dated with a relaxed molecular clock informed by 136 critically selected fossil-derived calibrations (Magallón *et al*., 2015), and the crown group age constrained within a confidence interval derived from a quantitative paleobiology method (Marshall, 2008). To our knowledge, it represents the most densely-calibrated molecular time-tree available. Using this comprehensive time-tree, we applied Bayesian analysis of Macroevolutionary Mixtures (BAMM; Rabosky, 2014: Rabosky *et al*., 2014), a method that through a Compound Poisson Process, identifies major shifts in the rate of diversification among phylogenetic branches and through time. To account for the possibility that BAMM produces posterior estimates of the number of diversification shifts that are indistinguishable from the prior (Moore *et al*., 2016), we conducted independent analyses covering a range of values for the prior on the number of expected shifts across the tree, and provide technical results. Our results identify the phylogenetic and temporal placement of diversification shifts associated with the origins of angiosperm megadiversity, and model angiosperm long-term diversification trajectory, suggesting ongoing species accumulation.

## Materials and Methods

### Taxonomic sample, molecular data and phylogenetic analyses

The diversification study is based on a previously published, temporally calibrated phylogenetic tree of angiosperms (Magallón *et al*., 2015). The taxonomic sample includes 792 angiosperms, six gymnosperms representing Cycadophyta, Gnetophyta and Coniferae, and a fern belonging to Ophioglossaceae. The angiosperms belong to 374 families, representing 87% of those recognized by the Angiosperm Phylogeny Website in April 2013 (Stevens, 2013), and encompass 99% of angiosperm total species-richness. The molecular data are the concatenated sequences of three plastid protein-coding genes (*atpB*, *rbcL* and *matK*) and two nuclear markers (18S and 26S nuclear ribosomal DNA), which form an alignment of 9089 base pairs (bp). We attempted to maximize data completeness by including sequences of species of the same genus when sequences of the same species were not available. When unavailable at the genus level, the sequence was left as missing data. Separate alignments of the sequences of different markers were conducted with MUSCLE v3.7 (Edgar, 2004), followed by manual refinements with BIOEDIT v7.0.9.0 (Hall, 1999). The sampled species and families, and GenBank accessions are shown in Table S1. The molecular data set is available from the author.

Phylogenetic estimation was conducted with maximum likelihood (ML) in RAxML v7.2.8 (Stamatakis, 2006a). The data was divided into four partitions: first and second codon positions of *atpB* plus *rbcL*; third codon positions of *atpB* plus *rbcL*; *matK;* and 18S plus 26S. Substitution parameters were estimated independently for each, using unlinked GTRCAT models. Topological constraints were implemented to specify phylogenetic relationships among major clades derived from an analysis based on a larger data matrix (Soltis *et al*., 2011). Trees were rooted rooted with the fern *Ophioglossum*. One thousand bootstrap replicates were implemented (Stamatakis *et al*., 2006b; Stamatakis, 2008).

### Dating analyses

The ML phylogenetic tree was dated by combining a method derived from quantitative paleobiology to constrain the age of the angiosperm crown node (Marshall, 2008), with an uncorrelated relaxed molecular clock to estimate dates within angiosperms (Drummond *et al*., 2006). Detailed information about dating analysis, including justification for all calibrations, is provided in the original study (Magallón *et al*., 2015). Briefly, the confidence interval on the age of the crown node was calculated considering the age of the oldest angiosperm fossils (Hughes & McDougall, 1986; Hughes *et al*., 1991; Brenner, 1996), and the number of branches in a family-level tree that are represented in the fossil record (Marshall, 2008).

Ages within angiosperms were estimated with the uncorrelated lognormal method in BEAST v1.7.5 (Drummond *et al*., 2006). The data were the alignment used for phylogenetic estimation (above) divided into plastid and nuclear partitions. A birth-death model was used as a tree prior. The root node, corresponding to the seed plant crown node, was calibrated with a uniform distribution between 314 and 350 Ma (Magallón *et al*., 2013). Internal nodes were calibrated with 136 fossils that were critically evaluated for their affinity to taxa represented in the tree, based either on morphological attributes, or, if available, on explicit phylogenetic placement. Prior age distributions of calibrated nodes were lognormal, with the mean equal to the fossil age plus 10%, and a standard deviation of 1. The calibration fossils, the attributes supporting clade affinity, their assignment to the stem or the crown node of the clade, their stratigraphic position and age are available in the original publication (Magallón *et al*., 2015) and from the author. The analysis consisted of eight independent MCMC runs for a total of 170 x 10^6^ steps, sampling one every 5000. The initial 600 trees from every run were excluded as burn-in. The outputs of the runs were analysed with Tracer v1.5 (Rambaut *et al*., 2014), and the estimated parameters joined with LogCombiner v1.7.5 and TreeAnnotator v1.7.5 (Drummond *et al*., 2006).

### Diversification Analyses

The macroevolutionary diversification dynamics of angiosperms were investigated with the C++ programme Bayesian Analysis of Macroevolutionary Mixtures (BAMM) v2.5.0 (Rabosky, 2014, 2017; Rabosky *et al*., 2014, 2017; Mitchell & Rabosky, 2016), which, through a compound Poisson process (CPP) implemented as a reversible jump Markov Chain Monte Carlo (rjMCMC), estimates major shifts in the rates of speciation, extinction, and diversification among the branches of a phylogenetic tree, and through time. Post-run analyses were conducted with the R package BAMMtools v2.5.0 (Rabosky *et al*., 2014).

### The BAMM method

BAMM simulates a posterior distribution of shift configurations, each corresponding to a particular combination of the number shifts (increases and decreases in diversification, speciation or extinction), and their phylogenetic and temporal placement among the branches of the tree. Given the posterior sample of shift configurations derived from the rjMCMC, it is possible to obtain a phylorate plot in which the rate of speciation, extinction or diversification averaged across all the configurations in the posterior distribution is plotted on each time unit on each branch (Rabosky, 2017).

In the BAMM model, the prior distribution of the expected number of shifts in the tree (expectedNumberOfShifts parameter) influences the number of shifts in the posterior distribution. Under the prior, the distribution of shifts across the tree is uniform, and the probability of finding a shift on a given branch depends on the specified prior for the number of shifts, and on the branch length (Rabosky, 2014, 2017). Thus, a high prior value specified for expectedNumberOfShifts will result in a high number of shifts in the posterior distribution, although many of them might be weakly supported by the data (Rabosky 2014, 2017). Significant shifts can be distinguished by considering the Marginal Odds Ratio (MOR) of a shift being present in a given branch. Because under the prior long branches are more likely to contain a shift, the MOR provides a measure of the amount of evidence supporting a shift on a given branch after normalizing for its length, that is, independently of the prior (Rabosky, 2017; Shi & Rabosky, 2015).

Moore and collaborators (Moore *et al*., 2016) pointed out that BAMM cannot provide reliable estimates of diversification rate shift models, or diversification rate parameters, mainly because its likelihood function does not account for rate shifts that took place on extinct branches; and, because there might be an infinite amount of equally likely CPP model parameterizations for the number and magnitude of shifts on the branches of the phylogeny, the posterior estimates are extremely sensitive to the prior number of shifts in the tree. After critically examining these claims, Rabosky and collaborators (Rabosky *et al*., 2017) concluded that although BAMM and (possibly except for Monte Carlo likelihood in Moore *et al*., 2016) all models that estimate rate shifts through time and on different parts of the phylogeny, do not account for rate shifts on extinct (or otherwise unobserved) branches; the effects that these unaccounted diversification shifts have in empirical cases are likely to be small. Also, although all Bayesian posterior estimates are influenced by the prior, it is possible to identify posterior estimates that are strongly supported independently from the prior (Rabosky *et al*., 2017). Hence, the use of Bayes Factor to select BAMM outcomes that are well supported, independently of the prior on the expected number of shifts, is strongly advocated (Rabosky, 2017). Specifically, in BAMM and BAMMtools 2.5.0 it is possible to estimate the MOR of a shift on a given branch (Rabosky, 2017). Diversification model selection with BAMM is largely robust to choice of model prior (Rabosky *et al*., 2017; Mitchell & Rabosky, 2016).

Identifying diversification shifts, and calculating diversification parameters across a phylogeny are complex estimation problems. We are aware that all currently available tools are imperfect, yet agree that BAMM v2.5.0 has been reasonably proven to achieve the task of providing realistic estimates of diversification parameters (Rabosky *et al*., 2017). While we recognize the potential problems in BAMM estimates of diversification shifts and rate parameters, to our knowledge no other currently available software can simultaneously achieve identification of rate shifts among phylogenetic branches and through time given the sampling density in this study (i.e., RPANDA [Morlon *et al*., 2011, 2016] requires at least three terminals per clade to evaluate different diversification models). We nevertheless exert caution regarding potential prior sensitivity when applying BAMM, and conduct several independent diversification analyses under a wide range of magnitudes for the prior of the expected number of shifts (see below). In each analysis we obtained the MOR associated to the presence of a shift on any branch, and made comparisons across analyses to identify a congruent set of highly supported shifts.

### Investigating Angiosperm Diversification

The angiosperm dated tree described above (Magallón *et al*., 2015) was used as input chronogram. BAMM was set to conduct a speciation-extinction analysis. We performed six analyses under different magnitudes for the prior of the expected number of shifts (expectedNumberOfShifts = 0.1, 1, 5, 10, 50, 100). Priors on rate parameters were scaled to our dated tree using the setBAMMpriors function in BAMMtools. The rate parameter of the exponential priors for the initial speciation and extinction values (lambdaInitPrior and muInitPrior) were both set to 4.66095688667462. The prior for the standard deviation of the normal distribution (mean fixed at zero) of the speciation rate regimes shift parameter (lambdaShiftPrior) was set to 0.00825917607286971. Constant diversification rate branch segments were set to 3 Ma by setting the segLength parameter at 0.02152148, given a crown node age of 139.3956 Ma. Rates were allowed to vary through time (lambdaIsTimeVariablePrior = 1).

The prior distribution of the number of rate shifts was calculated with BAMMtools v2.5.0 (Rabosky *et al*., 2014) by considering that the number of rate shifts follows a Poisson distribution with a rate parameter determined by an exponential hyperprior (Mitchell & Rabosky, 2016), and the finding that the probability of a given number of shifts in BAMM is the product of Poisson and exponential densities, which reduces to a simple geometric distribution on the number of rate shifts (Mitchell & Rabosky, 2016). Non-random incomplete taxon sampling of full angiosperm diversity was accounted for by indicating that clade-specific sampling probabilities would be used, and by specifying the sampled fraction of clades in the tree. Most of the clades correspond to angiosperm families recognized in the Angiosperm Phylogeny Website in April, 2013 (Stevens, 2013), with the total number of species in each family obtained from this same source. Families not represented in the dated tree were accounted for by aggregating their species-richness to that of their sister clade, according to relationships in the Angiosperm Phylogeny Website. Following BAMMtools documentation, for each terminal in the tree, we specified the represented fraction of the clade to which it belongs by dividing the total species-richness of the clade (i.e., a family or a family plus unsampled sister families) by the number of terminals belonging to that clade. The backbone of the phylogeny is fully sampled. Clade sampling fraction indicated on each terminal are shown in Table S2.

Each MCMC simulation consisted of 300 x 10^6^ steps, sampling a shift configuration every 200,000 steps. The initial 10% of the MCMC was discarded as burn-in, hence the total number of analysed posterior samples is 1351. The input tree and control files, and BAMM output event data files are available from the author. For each analysis, we calculated the phylorate plot (Fig. S1), and identified the best and the MSC configurations (available from authors).

For each analysis, we identified the shifts found in all the configurations in the posterior distribution, and estimated their MOR. We sorted the identified shifts by their MOR magnitude. We considered those shifts with a MOR value that falls within the 95% of the magnitude of the maximal MOR found in that analysis (e.g., if for a given analysis the maximal observed MOR is 100, we considered all the shifts with MOR ≥5). We then compared the shifts selected in each analysis across the six analyses, and chose to discuss only those that are shared by at least four analyses. To obtain the time of a shift on a given branch in each analysis, we extracted the age of the shift in all the configurations in which it was detected, and obtained the mean, minimal and maximal values (Fig. 2, Table S3).

**Table 1.**
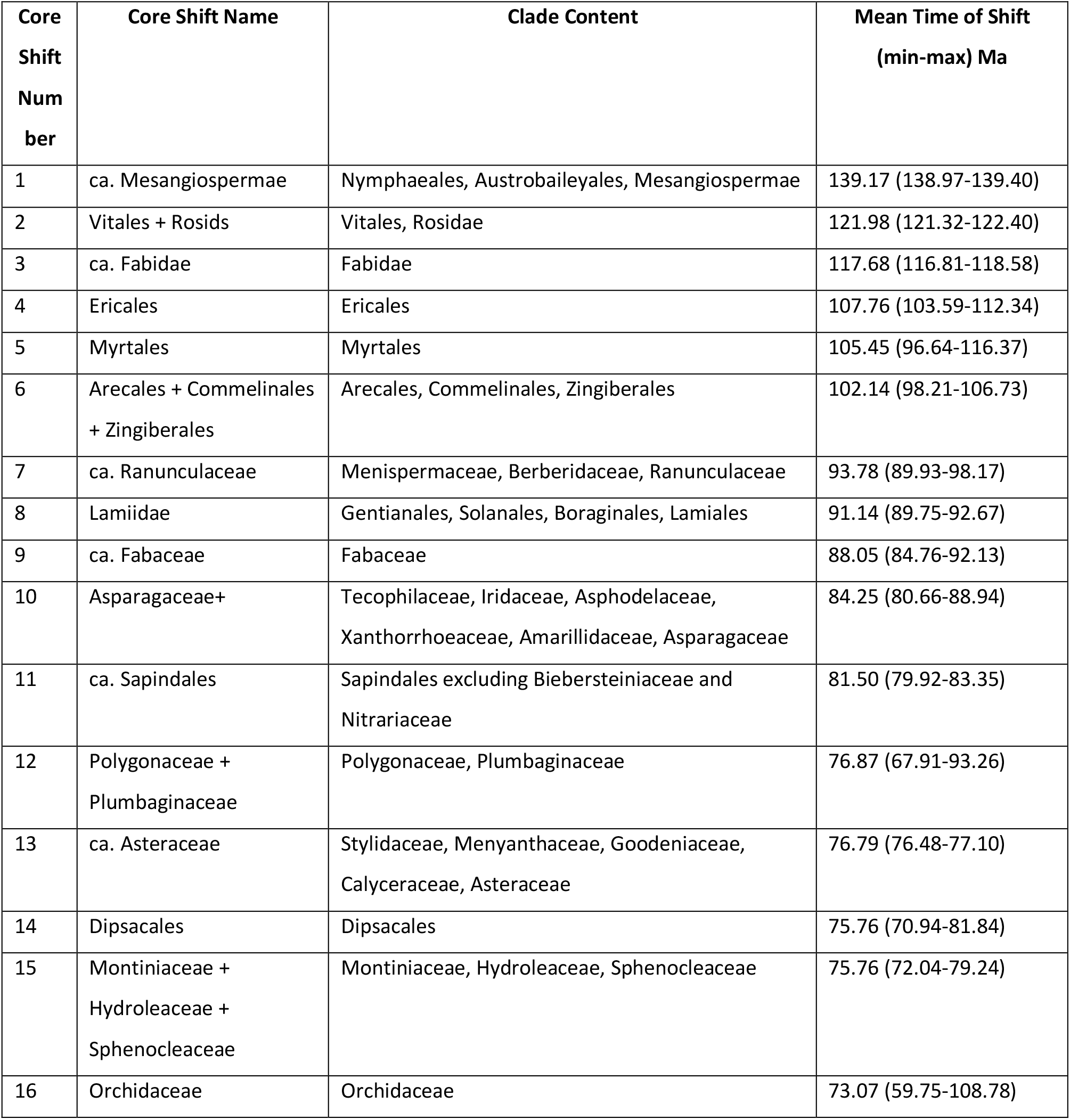
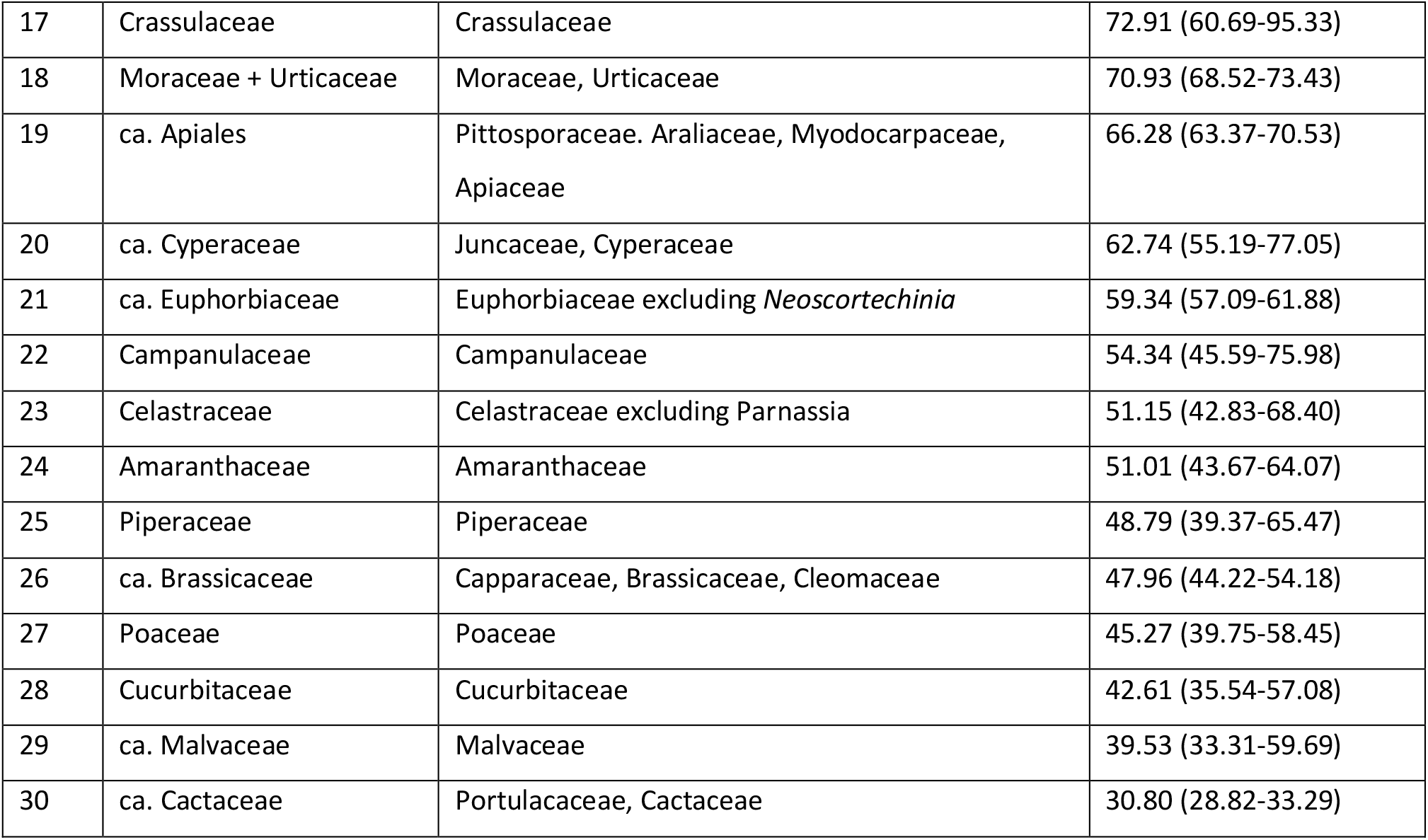
Thirty core angiosperm rate shifts. Core shifts correspond to those within 95% of the magnitude of the highest Marginal Odds Ratio (MOR) in each analysis with different prior for the expected number of shifts (i.e., 0.1, 1, 5, 10, 50, 100); and that are found in at least four (out of six) analyses. Core shifts detected on two or more adjacent branches are indicated as “ca. clade name”.

## Results

### Technical Results

BAMM is a method that, through a compound Poisson process (CPP) allows identification of diversification shifts among branches in the phylogenetic tree, and model their change through time (Rabosky, 2014; Rabosky *et al*., 2014). It also allows for estimation of the average rates of speciation, extinction and diversification of selected lineages, and for the calculation of their temporal trajectories. Being aware of potential nonidentifiability of the model on the number and distribution of diversification shifts, we conducted six independent analyses over a wide range of magnitudes for the prior on the expected number of shifts (expectedNumberOfShifts = 0.1, 1, 5, 10, 50 and 100). In all our analyses, the posterior distribution of the number of shifts is distinctly decoupled from the prior (Fig. 1). A larger magnitude on the prior on the expected number of shifts is less restrictive (Rabosky *et al*., 2017), and as expected, leads to the recognition of a larger number of shifts in the posterior distribution, each with a lower frequency (Fig. 1). Nevertheless, shifts that are highly supported by the data, as indicated by their MOR, are largely congruent among the six analyses (Tables S3, S4).

**Fig. 1.**
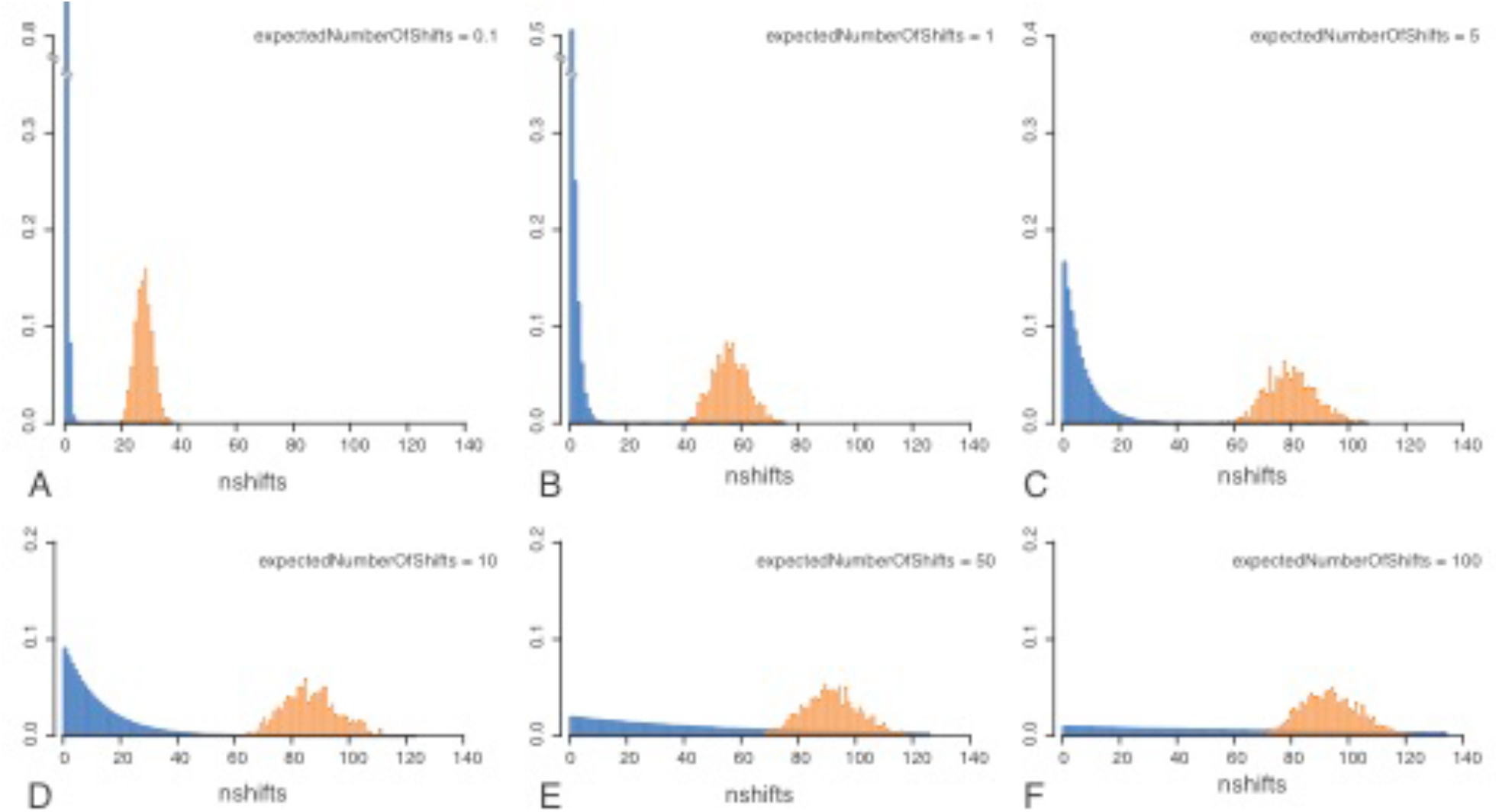
Prior and posterior distributions of the number of rate shifts. Prior (blue) and posterior (orange) distributions of the number of shifts in six BAMM analyses conducted under different prior values for the expectedNumberOfShifts parameter: A. 0.1; B. 1; C. 5; D. 10; E. 50; F. 100. In all cases, the prior and posterior distributions are distinct and non-overlapping. As expected (Rabosky *et al*., 2017), as the value on the prior on the expected number of shifts increases, the prior distribution becomes less informative, and the posterior distribution is wider, and centered around a higher value for the number of shifts. All plots are at the same scale, but the Y-axis in A and B was trimmed.

As the MOR allows to distinguish shifts that are strongly supported by the data, but cannot indicate the number of significant shifts in an analysis, we chose to discuss those shifts that, first, in any given analysis have a MOR that falls within the 95% of the maximum MOR value identified in that analysis; and, from the previous set, those that are shared among at least four of the six analyses. We identified 30 such shifts, and consider them as the core diversifications to be discussed (Tables 1, S4). In each analysis, the distribution of shifts sorted by their MOR is a hollow curve (Fig. S2, Table S5). Eighteen core shifts are distributed on single branches, and twelve drift in two or more adjacent branches, within a particular phylogenetic region (Fig. 2, Table S4). Twenty-six core shifts are towards increased diversification rates; three core shifts contain many nested shifts, and only one core shift is towards decreased diversification (Figs. 2, 3).

**Fig. 2.**
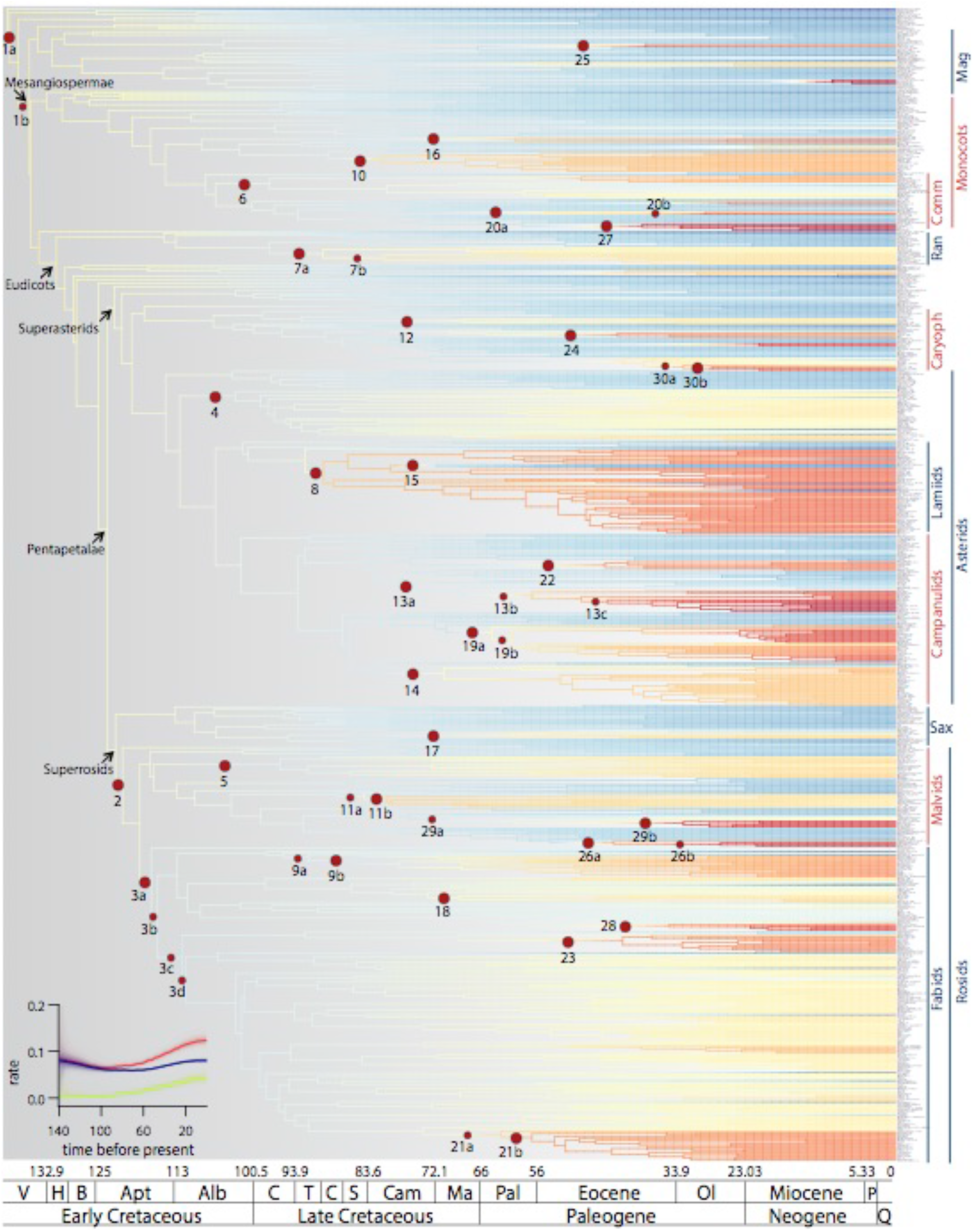
Angiosperm diversification phylorate plot. Rate of diversification per time interval (3 Ma) averaged across all configurations within the 95% credible set obtained in the BAMM analysis with the prior for expected number of shifts = 100. Major angiosperm clades (APG IV, 2016) are indicated with arrows. In chronological order, core shifts correspond to: 1. ca. Mesangiospermae; 2. Vitales + Rosids; 3. ca. Fabidae; 4. Ericales; 5. Myrtales; 6. Arecales + Commelinales + Zingiberales; 7. ca. Ranunculaceae; 8. Lamiidae; 9. ca. Fabaceae; 10. Asparagaceae + Amarillidaceae + Xanthorrhoeaceae + Asphodelaceae + Iridaceae + Tecophilaceae; 11. ca. Sapindales; 12. Polygonaceae + Plumbaginaceae; 13. ca. Asteraceae; 14. Dipsacales; 15. Montiniaceae + Hydroleaceae + Sphenocleaceae; 16. Orchidaceae; 17. Crassulaceae; 18. Moraceae + Urticaceae; 19. ca. Apiales; 20. ca. Cyperaceae; 21. ca. Euphorbiaceae; 22. Campanulaceae; 23. Celastraceae; 24. Amaranthaceae; 25. Piperaceae; 26. ca. Brassicaceae; 27. Poaceae; 28. Cucurbitaceae; 29. ca. Malvaceae; 30. ca. Cactaceae. See Table 1 and Supplementary Table 1 for description of the content of each clade. Some core shifts consist of moderately supported shifts on two or more adjacent branches, and are indicated with letters (e.g., 1a, 1b). The sub-shift indicated with a larger circle has the highest Marginal Odds Ratio (MOR). The inset shows the Diversification-Through-Time (DTT) plot for angiosperms as a whole, including graphs for diversification (blue), speciation (red) and extinction (green).

**Fig. 3.**
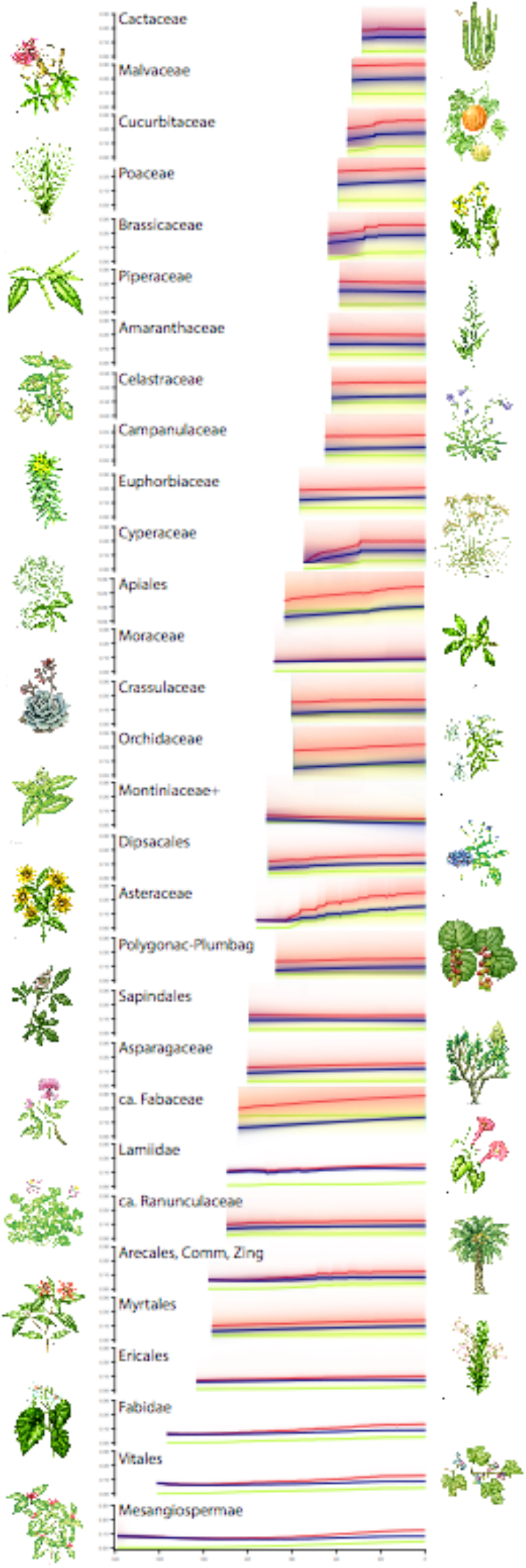
Diversification Through Time plots for core shift clades. Diversification Through Time (DTT) plots for clades resulting from core diversification shift, sorted by onset time (Ma; from bottom to top), including graphs for diversification (blue), speciation (red) and extinction (green).

### Angiosperm Diversification

The onset of angiosperm diversification into extant lineages in the Early Cretaceous was soon followed by the differentiation of Mesangiospermae (Cantino *et al*., 2007) – a clade that contains the vast majority of angiosperm’s diversity, morphological variety and ecological breadth – and its diversification into five evolutionary lineages, including the Magnoliidae (magnoliids), Monocotlyedoneae (monocots), and Eudicotyledoneae (eudicots). Most eudicots belong to the large clade Pentapetalae (Cantino *et al*., 2007) which in turn is composed of Superrosids and Superasterids (APG IV, 2016; Fig. 2). Together, monocots, Superrosids, and Superasterids include approximately 95% of living angiosperm species.

Core diversification shifts are distributed all across the angiosperm phylogeny, and took place between the Early Cretaceous (Valanginian) to the latest Eocene (Priabonian) or earliest Oligocene (Rupelian), spanning a period of over 100 million years (Fig. 2). Six core diversification shifts took place in the Early Cretaceous. The first is approximately associated with the origin of Mesangiospermae (1a-1b; Tables 1, S4), and is followed by shifts along the spine of Superrosids, including one on the branch subtending Rosids (2), and another spanning from Fabids to a clade formed by Oxalidales and Malpighiales (3a-3d). Each of these three core shifts contain many nested shifts. Later during the Early Cretaceous one shift took place within Asterids, subtending Ericales (4); another one within Rosids, in Malvids, subtending Myrtales (5), and a third one within monocots, subtending a clade approximately corresponding to Commelinids (6, Fig. 2, Tables 1, S4). Unless otherwise noted, all shifts represent increasing diversification.

During the Late Cretaceous, 15 core shifts gave rise to clades within eudicots and monocots. The earliest shift took place within eudicots, corresponding approximately to Ranunculaceae (7a-7b). All other eudicot shifts are nested in Pentapetalae. Within Superrosids, there was shift in Saxifragales, subtending Crassulaceae (17); another one within Malvids, corresponding approximately to Sapindales (11a-11b); and three shifts within Fabidae: approximately Fabaceae (9a-9b), Moraceae plus Urticaceae (18), and approximately Euphorbiaceae (21a-21b). Within Superasterids, one shift took place in Caryophyllales, subtending Polygonaceae plus Plumbaginaceae (12); two shifts in Lamiidae, corresponding to a clade containing Solanales, Gentianales, Boraginales, and Lamiales (8), and nested within it, a shift towards decreased diversification in a clade formed by Montiniaceae, Hydroleaceae, and Sphenocleaceae (15); and three shifts in Campanulidae: approximately Asteraceae (13a-13c), Dipsacales (14), and approximately Apiales (19a-19b). Three shifts took place within monocots: a clade containing Asparagaceae and five additional families (10), Orchidaceae (16), and approximately Cyperaceae (20a-20b; Fig. 2, Tables 1, S4).

Nine core shifts took place during the Paleogene. Within Rosids there were two shifts in Malvids, corresponding approximately to Brassicaceae (26a-26b), and approximately to Malvaceae (29a, 29b); and two shifts in Fabidae: Celastraceae (23) and Cucurbitaceae (28). There were two shifts in Superasterids, both within Caryophyllales, corresponding to Amaranthaceae (24), and approximately to Cactaceae (30a-30b). There was a single shift within Asterids, in Campanulids, subtending Campanulaceae (22), and also a single shift within monocots, corresponding to Poaceae (27). A single core shift was detected within magnoliids, subtending Piperaceae (25; Fig. 2, Tables 1, S4). However, two additional strong diversification increases, both within magnoliids, are noticeable in the phylogram (Fig. 2), corresponding to Annonaceae (Magnoliales) and to a clade that includes *Cinnamomum*, *Sassafras* and *Laurus*, within Lauraceae (Laurales). These diversification increases were not identified as core shifts according to the delimiting criteria.

Shift times and diversification parameters of angiosperms as a whole, and of the 30 core shifts, estimated in analyses with different prior for the number of expected shifts are very similar (Tables S3, S4). We discuss diversification parameters and times derived from the analysis with the prior for expected number of shifts equal to 100 because it includes the 30 core shifts selected through our combined criteria (Table 1). Estimated speciation (*λ*) and extinction (*μ*) rates for angiosperms as a whole are 0.0988 (0.0910-0.1086) and 0.0277 (0.0192-0.0384), respectively, which are congruent with previous estimates (Magallón & Sanderson, 2001).

Diversification-through-time (DTT) plots for angiosperms as a whole estimated with different values for the prior on the expected number of shifts are virtually equal. The rate of diversification is modelled as having undergone a moderate early decline (ca. 140-100 Ma), followed by stabilization (ca. 100-50 Ma) and moderate increase towards the present (Fig. 2). Speciation and extinction rates are modelled as having a pronounced increase towards the present (Fig. 2). Individual shift clades have widely varying increasing or decreasing DTT trajectories, which overlap through time (Fig. 3).

## Discussion

### Implications of backbone sampling

The identified shifts provide information of major diversification changes for angiosperms as a whole at a deep phylogenetic scale. They represent an important starting point to understand major features of angiosperm evolution (Uyeda *et al*., 2017). The phylogenetic level at which shifts were identified is a function of the used backbone sampling, in which major angiosperm lineages were represented by a small number of placeholders, which convey limited information about species richness distributed among and within angiosperm clades. The taxonomic selection in the dated phylogenetic tree aimed to represent, as much as possible, all angiosperm clades recognized as families. This type of selection results in a highly biased depiction of species distribution, as members of very small clades, with a very low probability of being represented under random sampling, were selectively sampled. Each family, regardless of its species richness, is represented by a similar number of placeholders, which correspond to very different proportions of the total species richness in each clade. This biased sampling is necessary if as many as possible angiosperm families are to be included, but has important consequences when attempting to estimate macroevolutionary parameters. To mitigate the effect of biased sampling, we specified clade-specific sampling fractions in diversification analyses. Sampling fractions varied widely, from 1.0 (e.g., Amborellaceae, Trochodendraceae, Eucommiaceae), to <0.0003 (e.g., Rubiaceae, Lamiaceae, Orchidaceae; Table S2). Not many viable alternatives are available. Ideally, the difference in sampling fractions among clades could be reduced by very greatly increasing sampling within large families, although this strategy might be limited by molecular data availability, and contingent on the capability of models and computer power to handle taxonomically massive datasets in macroevolutionary analyses. Another potential strategy is to implement random sampling among angiosperms, but after including representatives of clades unlikely to be sampled randomly due to their small size (e.g., O’Meara *et al*., 2016). Bioinformatic methods to incorporate missing taxa into backbone phylogenies are available (e.g., PASTIS, Thomas *et al*., 2013), but given the colossal species richness of many angiosperm families, and the very small number of placeholders in the phylogenetic backbone, we suspect this type of approach would be unfeasible in this study.

As a consequence of extremely reduced sampling, estimates of diversification dynamics can only associate shifts to major clades, usually on their stem lineage, as information (i.e., species sampling) that could document one or more shifts within the clade is lacking. The reduced taxonomic sampling also precludes replicating diversification shifts detected in studies focused on delimited, and much more densely sampled clades (e.g., Lagomarsino *et al*., 2016). The absence of diversification shifts younger than the Paleogene may also be a consequence of sparse taxonomic sampling.

### Previous Angiosperm Diversification Studies

Few previous studies have investigated long term angiosperm diversification dynamics, in particular, identifying diversification shifts (Table 2). Davies and collaborators (Davies *et al*., 2004) conducted the first study to identify diversification shifts at an angiosperm-wide scale, using a family-level supertee, in which diversification shifts were approximated with species richness imbalance among families (Fusco & Cronk, 1995; Purvis *et al*., 2001), from which they detected the most (top ten) imbalanced family pairs. Of these, five may correspond to shifts detected in our study (Table 2). Smith and collaborators (Smith *et al*., 2011) applied SymmeTREE (Chan & Moore, 2005) – a method that considers the topological distribution of species richness to fit models of constant or variable rates (Chan & Moore, 2002) – to an angiosperm megaphylogeny. They identified between 16 and >2700 potential shifts, which were not named. The closest precedent to our study is the analysis of Tank and collaborators (Tank *et al*., 2015), who investigated links between diversification shifts and whole genome duplications, by applying MEDUSA (Alfaro *et al*., 2009) – a maximum likelihood stepwise method to identify the best-fitting diversification model and detect significant shifts in diversification and relative extinction – to a set of bootstrapped chronograms representing 325 angiosperm families. Over 140 unique shifts were identified across all bootstrapped chronograms, 27 of which occurred in at least 75% of all the set (Table 2 in Tank *et al*., 2015). Of these, nine are approximately equivalent to shifts detected here (Table 2). In spite of some profound sampling and methodological differences, as well as analytical limitations associated to each method, it is noteworthy that these studies detect partially overlapping sets of diversification shifts, the most recurrent ones being associated to Piperaceae, Asparagaceae and related families, a clade containing Arecales, Commelinales and Zingiberales, Cyperaceae, Poaceae, Cactaceae, ca. Lamiidae, Asteraceae, Myrtaceae, Brassicaceae, and Fabaceae.

**Table 2.**
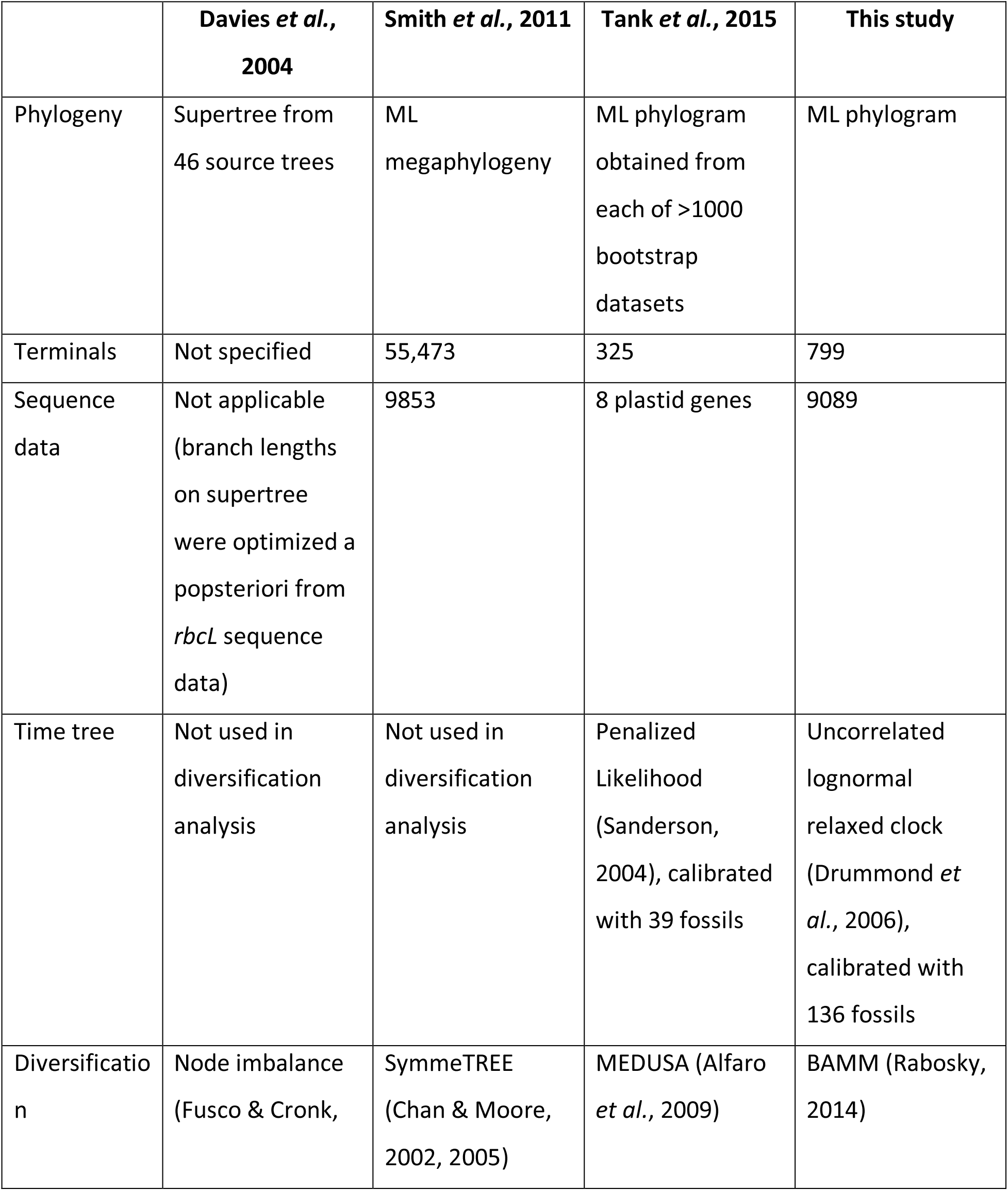
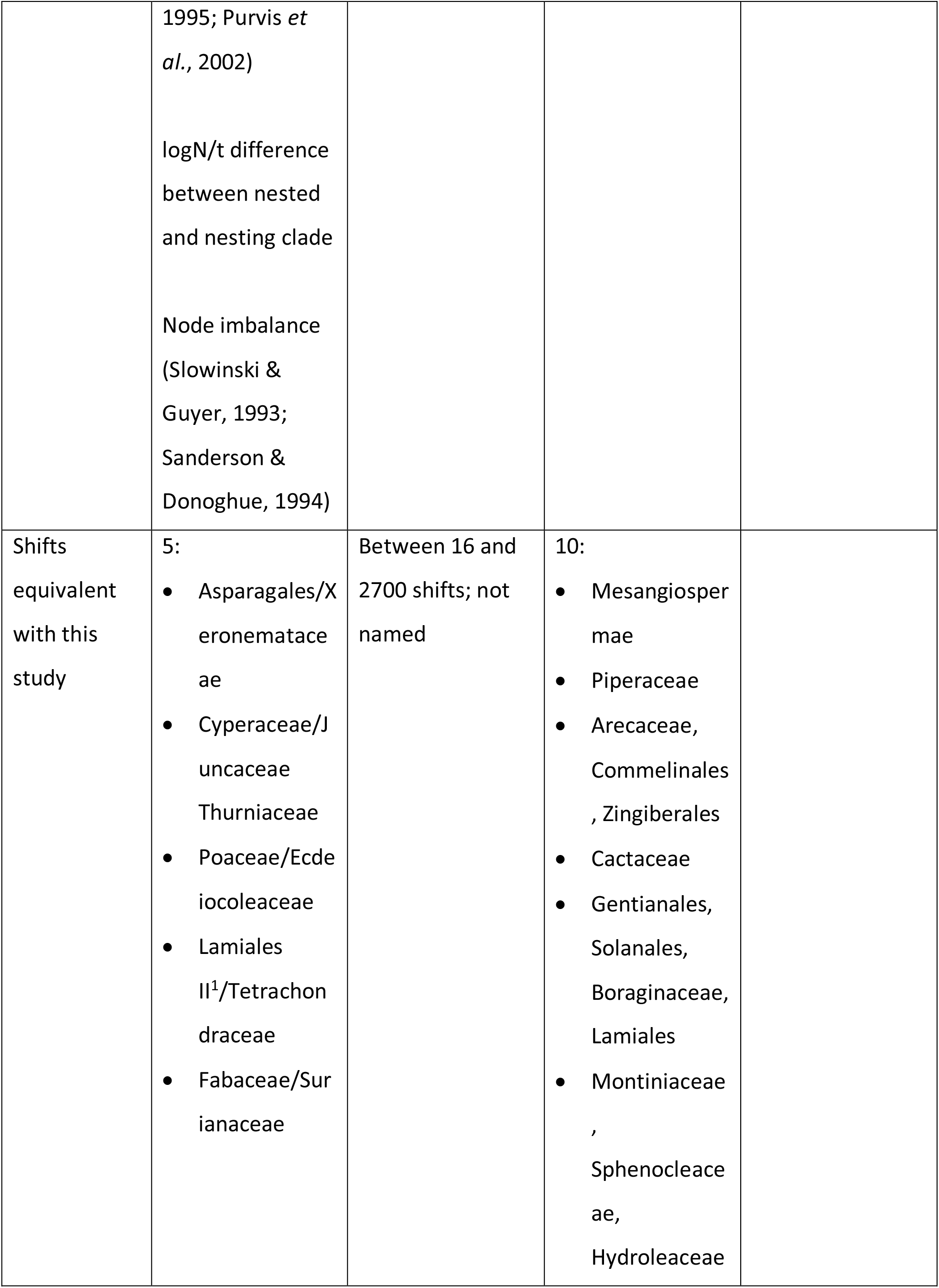
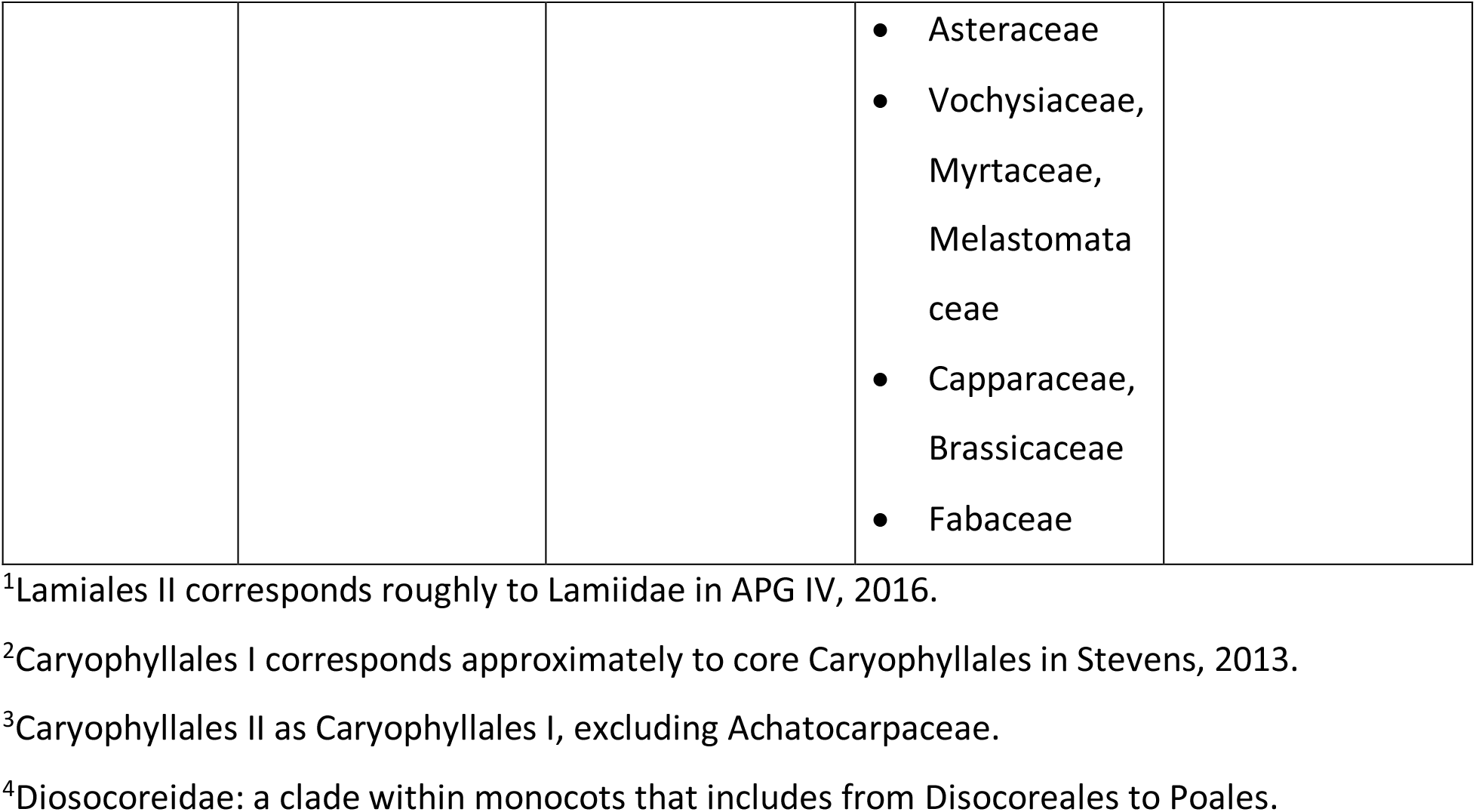
Published studies identifying angiosperm diversification shifts. Comparative summay of published studies in which shifts in the rate of diversification within angiosperms have been identified.

### Phylogenetic placement of radiations

The unequal distribution of species richness among lineages in different biological groups has for long attracted the interest of evolutionary biologists. Angiosperms are an emblematic example, in which clades exhibit pronounced differences in the number of extant species they contain. This study conclusively confirms previous suggestions (Magallón & Sanderson, 2001; Davies *et al*., 2004; Tank *et al*., 2015) that angiosperm megadiversity is not directly associated with the origin of angiosperms as a whole, but rather, results from several independent diversification shifts within the clade, and goes beyond by identifying particular phylogenetic regions in which diversification shifts have taken place, documenting a complex pattern of temporally overlapping radiations and depletions in different lineages that result in present-day distribution of species richness across angiosperms.

Most core shifts subtend species-rich clades identified as distinct botanical orders and families that, while containing wide disparity in morphology and ecological function, are nevertheless each characterized by a distinctive combination of vegetative attributes and/or reproductive groundplans (e.g., Fabaceae, Asteraceae, Orchidaceae, Poaceae; Table 1). The placement of diversification shifts associated with distinct groundplans suggests that diversification shifts are associated with the evolution of integrated morphological combinations that achieve particular complex functions (O’Meara *et al*., 2016). Nevertheless, as discussed above, the sparse taxonomic sample in this study, in which massive clades are represented by only a few species, precludes detecting radiations that might have taken place within those clades.

The finding of shifts on adjacent branches is technically consistent with the fact that shifts identified in different configurations by the CPP in BAMM are not independent from each other, but most likely represent a shift detected with moderate marginal probabilities on adjacent branches (Rabosky, 2017).

### Have there been diversification decreases in angiosperm evolution?

The extraordinary present-day diversity of angiosperms makes it easy to overlook that some angiosperm lineages may be in decline. Our results document that, while some lineages within angiosperms represent exceptional evolutionary radiations, others are decreasing. Diversification through time plots show that lineages within angiosperms underwent independent radiations and depletions at different times, which overlapped and masked each other, suggesting that different lineages were predominant at different times through angiosperm history (Fig. 3).

Our results show pronounced variations in the rates of diversification among angiosperm lineages (Table S3). Many high rate clades derive from distinct diversification shifts, and are scattered across the phylogeny (Fig. 2). We observe that lineages with low diversification rates occupy two distinct types positions in the phylogeny: There are low diversification grades that subtend large clades with high diversification rates. These grades correspond to the depauperons discussed by Donoghue and Sanderson (2015). There are also isolated low diversification branches (or small clades) embedded within a speciose clade characterized by high diversification rate. We hypothesize that low diversification lineages in these two distinct phylogenetic positions result from differential evolutionary situations, and are characterized by distinct diversification dynamics. The low diversification grades have been explained as branches that differentiated before the evolution or during the assembly of traits or conditions associated with the increased diversification of the speciose sister clade (Donoghue & Sanderson, 2015). Thus, the rates of low diversification grades possibly represent retained plesiomorphic conditions. On the other hand, isolated low diversification branches are probably outcomes of diversification shifts towards decreasing rates that affected that particular branch. In fact, the single decreasing core diversification shift detected in this study (core shift 15, Fig. 2, Table 1) is associated with an isolated low diversification clade (i.e., Montiniaceae, Sphenocleaceae, Hydroleaceae), within a high diversification, speciose clade (i.e., Lamiidae). Although only one isolated low diversification clade was identified as resulting from a core shift, many others can be identified in the phylorate plot (e.g., Plocospermataceae, Roridulaceae, Barbeyaceae-Dirachmaceae, Goupiaceae; Fig. 2).

The relatively few detected diversification decreases probably do not reflect the paucity of lineages in decline, but rather, as the natural ultimate outcome of decreasing diversification is extinction, lineages undergoing an evolutionary depletion persist shortly. Detection of recent diversification decreases is more likely, appearing as depauperate lineages on their way to extinction. Plocospermataceae, the sister group to the remainder of Lamiales, is a possible example (Fig. 2). Lineages that underwent an ancient diversification decrease but survive to the present are unexpected and difficult to explain (Strathmann & Slatkin, 1983; Magallón & Sanderson, 2001). These lineages might be in decline from former megadiversity and ultimate demise is taking longer, or they might have recovered after a drastic decrease. The clade containing Monitiniaceae, Hydroleaceae and Spehnocleaceae (Fig. 2, shift 15) is a possible example.

### Are angiosperms in decline?

Previous work (Vamosi & Vamosi, 2011) has suggested that extrinsic factors can set limits to angiosperm diversification. The long-term diversification trajectory of angiosperms modelled here includes a stable to slightly increasing trend towards the present. This sustained diversification trajectory indicates that angiosperms as a whole are not undergoing a diversification decline, but rather that species-richness will continue to accumulate. The trends of increasing speciation and extinction (Fig. 2) imply a higher species turnover. Although angiosperms are today the most diverse group of plants in terrestrial ecosystems, where they display exceptional morphological, functional and ecological complexity and innovation, the estimated diversification trajectory indicates that angiosperm evolutionary expansion remains unmitigated. The existence of limits to species accumulation in angiosperm as a whole remains to be evaluated, but our results suggest that if such limits exist, they have not yet been reached. These results are congruent with the finding of a non-equilibrium phase in the acquisition of floral structure diversity associated with increasing diversification rates (O’Meara *et al*., 2016).

Angiosperm DTT plots were here modelled considering a sample of lineages that diverged tens of million of years ago, and as such, cannot predict potential changes in trajectory in response to drastic environmental changes (e.g., climate change) that were not modelled in the simulation. These plots could be complemented with graphs estimated with methods that can directly incorporate the fossil record (Stadler, 2011), or information about relevant paleoenvironmental variables, such as temperature (Condamine *et al*., 2013; Sauquet & Magallón, in review).

### Are there phylogenetic regions or times in which diversification increases or decreases are concentrated?

The origin of angiosperm megadiversity cannot be traced to a few events or to a restricted time interval, but rather, results from many independent diversification shifts through most of its evolutionary history and across its phylogenetic spectrum. Namely, a concentration of diversification shifts in response to major global events such as the K/T mass extinction of the end-Eocene cooling event is not observed. While floristic shifts are detected in local paleofloras (Nichols & Johnson, 2008; Barreda *et al*., 2012), the absence of a distinct concentration of diversification shifts, or marked changes in direction in DTT plots around any particular time suggests that responses in angiosperm composition to global events most likely took place at lower phylogenetic scales.

### Perspectives and Emerging Questions

In this study, we detected diversification shifts at a deep macroevolutionary level. The identified diversification shifts represent clues to recognizing possible causes and ultimate drivers of megadiversity, which likely combined intrinsic attributes, ecological functions and abiotic conditions. Most variables that have been postulated as drivers of angiosperm diversification are complex structures and functions resulting from additive integration of simpler attributes through the phylogenetic history of different lineages (O’Meara *et al*., 2016). The clades here detected as resulting from diversification increases lack shared potential “key attributes” that could commonly underlie their respective diversifications. Furthermore, each of these clades includes many intrinsic attributes deployed in a great variety of extrinsic conditions that could potentially be related with increased diversification (Sauquet & Magallón, in review). While this study is not intended to recognize the causal factors underlying angiosperm diversification, it provides a critical framework (Uyeda *et al*., 2017) by pointing to particular phylogenetic regions to investigate for diversification associated to intrinsic or extrinsic factors (Bouchenak-Khelladi *et al*., 2015). A further improved understanding of angiosperm radiations, and of the causes that drive them, should necessarily be based on a more precise phylogenetic location of incremental and decremental diversification shifts, derived from a much denser taxonomic sampling, namely species-level phylogenies, and ideally, direct integration of fossil information.

## Acknowledgements

We thank H. Sauquet, A. Benítez-Villaseñor, R. Hernández-Gutiérrez, A. López-Martínez and S. Ramírez-Barahona for comments. G. Ortega-Leite, A. Wong-León, J.C. Montero-Rojas, D. Martínez-Almaguer and A. Luna provided technical assistance. LLSR thanks the Consejo Nacional de Ciencia y Tecnología México for a scholarship (CONACyT 262540), and the Posgrado en Ciencias Biológicas, Universidad Nacional Autónoma de México (UNAM) for support. SLGA thanks the Dirección General de Asuntos del Personal Académico, UNAM, for postdoctoral funding.

## Author Contributions

SM designed research; SM and LLSR performed research; SLGA and LLSR contributed data; SM and LLSR analysed data; SM, with contributions of LLSR, wrote the paper.

## Supporting Information

**Fig S1.** Phylorate plots of mean diversification rate. Phylorate plots showing the mean rate of diversification for each time interval (3 Ma) in each branch, averaged across the configurations in the posterior distribution resulting from diversification analyses conducted with a different magnitude for the prior of expected number of shifts: A = 0.1; B = 1; C = 5; D = 10; E = 50; F = 100.

**Fig. S2.** Distribution of shift Marginal Odds Ratio (MOR). Distribution of MOR of shifts detected in diversification analyses conducted with different prior magnitudes for expected number of shifts: A = 0.1; B = 1; C = 5; D = 10; E = 50; F = 100. The magnitude of MORs varies among analyses, but in all cases, their distribution corresponds to a hollow curve in which few shifts have high MORs, and most have low MORs.

**Table S1.** Species list and GenBank accessions.

**Table S2.** Sampling fraction associated to each terminal.

**Table S3.** Thirty core angiosperm rate shifts, including shifts in adjacent branches in a distinct phylogenetic region.

**Table S4.** Thirty core angiosperm rate shifts, including shifts in adjacent branches in a distinct phylogenetic region.

**Table S5.** Values of Marginal Odds Ratio (MOR) and associated parameters.

