## Supplementary Materials for "Thirty clues to the exceptional diversification of flowering plants"

Figure S1: page 2

Figure S2: page 3

Table S1: page 4

Table S2: page 39

Table S3: separate file

Table S4: page 62

Table S5: page 67

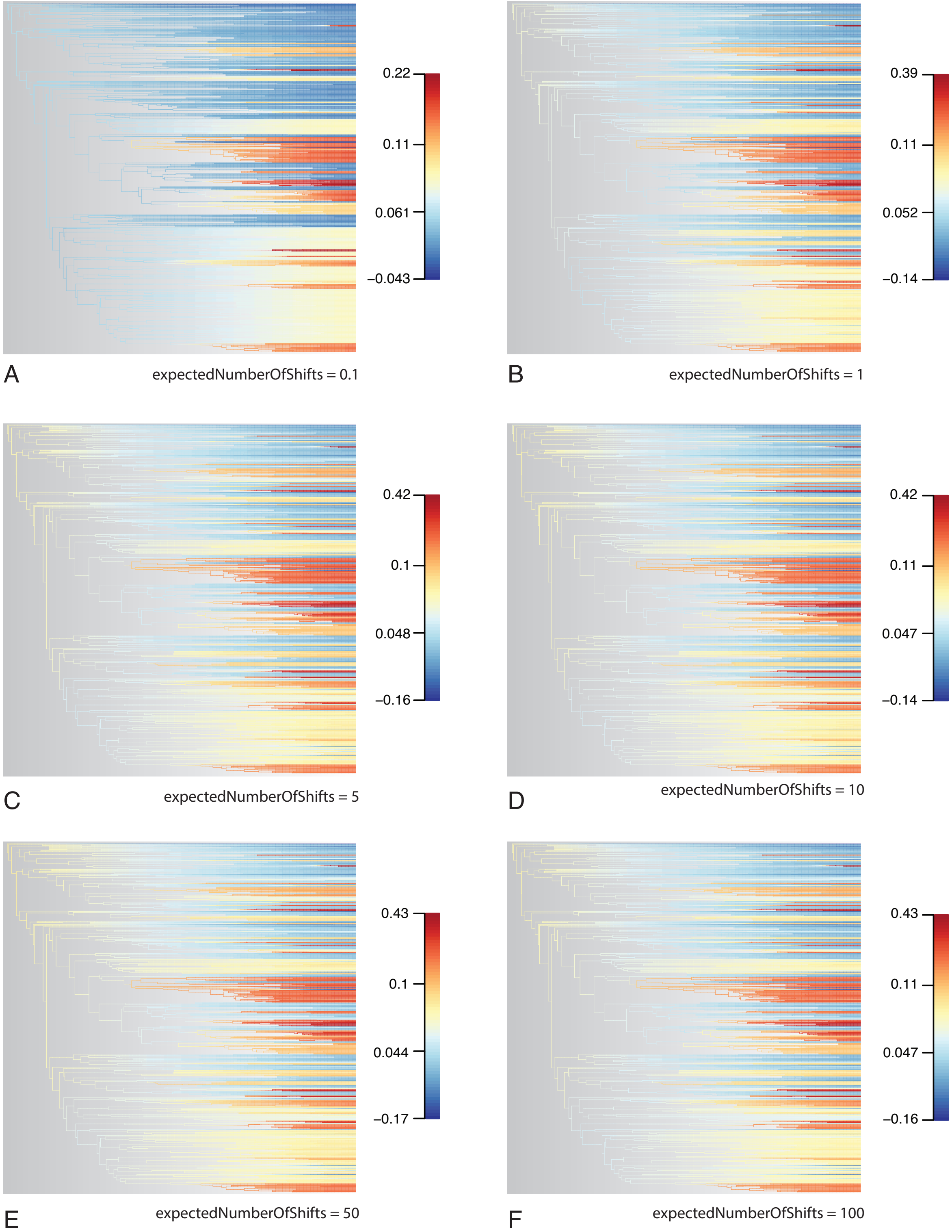
0.22
0.11
0.061
−0.043
A
expectedNumberOfShifts = 0.1
0.39
0.11
0.052
−0.14
B
expectedNumberOfShifts = 1
0.42
0.1
0.048
−0.16
C
expectedNumberOfShifts = 5
0.42
0.11
0.047
−0.14
D
expectedNumberOfShifts = 10
0.43
0.1
0.044
−0.17
E
expectedNumberOfShifts = 50
0.43
0.11
0.047
−0.16
F
expectedNumberOfShifts = 100

Figure S1

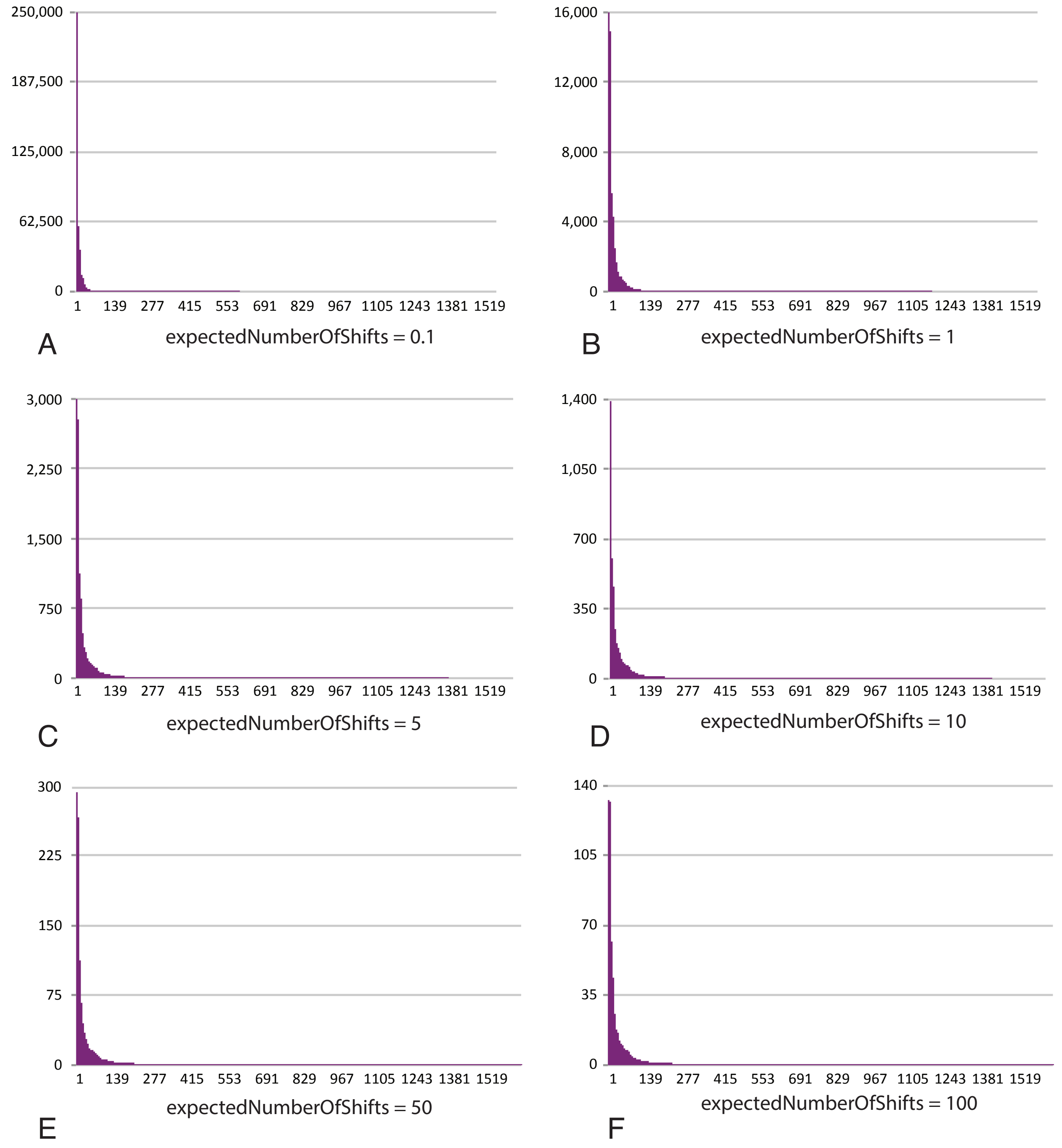
250,000
187,500
125,000
62,500
0
1 139 277 415 553 691 829 967 1105 1243 1381 1519
A
expectedNumberOfShifts = 0.1
16,000
12,000
8,000
4,000
0
1 139 277 415 553 691 829 967 1105 1243 1381 1519
B
expectedNumberOfShifts = 1
3,000
2,250
1,500
750
0
1 139 277 415 553 691 829 967 1105 1243 1381 1519
C
expectedNumberOfShifts = 5
1,400
1,050
700
350
0
1 139 277 415 553 691 829 967 1105 1243 1381 1519
D
expectedNumberOfShifts = 10
300
225
150
75
0
1 139 277 415 553 691 829 967 1105 1243 1381 1519
E
expectedNumberOfShifts = 50
140
105
70
35
0
1 139 277 415 553 691 829 967 1105 1243 1381 1519
F
expectedNumberOfShifts = 100

Figure S2

**Supplementary Table 1.** Species list and GenBank accessions. List of species included in diversification analyses, including the order and family to which they belong, and GenBank accession.

| Order | Family | Species | *rbcL* | *atpB* | *matK* | 18S | 26S |
|---|---|---|---|---|---|---|---|
| Amborellales | Amborellaceae | *Amborella trichopoda* | L12628.2 | AJ235389.1 | AF543721.1 | U42497.1 | AF479238.1 |
| Nymphaeales | Hydatellaceae | *Trithuria submersa* | DQ915188.1 | AJ419142.1 | | | |
| Nymphaeales | Hydatellaceae | *Trithuria austinensis* | | | JQ284130.1 | | |
| Nymphaeales | Nymphaeaceae | *Nymphaea odorata* | M77034.1 | AJ235544.1 | DQ185549.1 | AF206973.1 | |
| Nymphaeales | Nymphaeaceae | *Nymphaea sp.* | | | | | AY095465.1 |
| Nymphaeales | Nymphaeaceae | *Nuphar advena* | DQ354691.1 | DQ354691.1 | | | |
| Nymphaeales | Nymphaeaceae | *Nuphar lutea* | | | AF117100.1 | | |
| Nymphaeales | Nymphaeaceae | *Nuphar japonica* | | | | D85295.1 | |
| Nymphaeales | Nymphaeaceae | *Nuphar sp.* | | | | | DQ008660.2 |
| Nymphaeales | Nymphaeaceae | *Cabomba caroliniana* | M77027.1 | AF187058.1 | DQ185527.1 | AF096691.1 | AF479239.1 |
| Nymphaeales | Nymphaeaceae | *Brasenia schreberi* | M77031.1 | AF209544.1 | DQ185529.1 | AF206874.1 | DQ008661.2 |
| Austrobaileyales | Austrobaileyaceae | *Austrobaileya scandens* | L12632.2 | AJ235403.1 | DQ401347.1, 77743621 | AF206858.1 | AY095452.1 |
| Austrobaileyales | Trimeniaceae | *Trimenia moorei* | 7580491 | AY116653.1 | DQ401360.1 | | AY095470.1 |
| Austrobaileyales | Schisandraceae | *Illicium floridanum* | DQ182334.1 | | AF543738.1 | | |
| Austrobaileyales | Schisandraceae | *Illicium parviflorum* | 37194806 | U86385.2 | | L75832.1 | |
| Austrobaileyales | Schisandraceae | *Illicum verum* | | | | | EU161362.1 |
| Austrobaileyales | Schisandraceae | *Schisandra chinensis* | AF238061.1 | AF239790.1 | | AF094561.1 | |
| Austrobaileyales | Schisandraceae | *Schisandra rubriflora* | | | AF543750.1 | | |
| Austrobaileyales | Schisandraceae | *Schisandra sphenanthera* | | | | | DQ008658.1 |
| Austrobaileyales | Schisandraceae | *Kadsura japonica* | AF197595.1 | AF197607.1 | DQ185525.1, 77743625 | AF293763.1 | DQ008657.1 |
| Chloranthales | Chloranthaceae | *Hedyosmum arborescens* | L12649.2 | AJ235491.1 | DQ401339.1 | AF206925.1 | AF479226.1 |
| Chloranthales | Chloranthaceae | *Ascarina lucida* | 9623108 | AF239775 | | | |
| Chloranthales | Chloranthaceae | *Ascarina sp.* | | | 89242557 | | |
| Chloranthales | Chloranthaceae | *Ascarina rubricaulis* | | | | | 66969244 |
| Chloranthales | Chloranthaceae | *Sarcandra chloranthoides* | AY236833.1 | | AJ966796.1 | | DQ008655.1 |
| Chloranthales | Chloranthaceae | *Sarcandra grandiflora* | | AJ235593.1 | | 7595532 | |
| Chloranthales | Chloranthaceae | *Sarcandra glabra* | | | | AF094536.1 | |
| Chloranthales | Chloranthaceae | *Chloranthus japonicus* | L12640.2 | AJ235431.2 | | | AF479245.1 |
| Chloranthales | Chloranthaceae | *Chloranthus brachystachys* | | | AF543733.1 | | |
| Chloranthales | Chloranthaceae | *Chloranthus spicatus* | | | | D29787.1 | |
| Canellales | Canellaceae | *Cinnamodendron ekmanii* | | AJ235435.1 | AF465290.1 | AF206887.1 | |
| Canellales | Canellaceae | *Cinnamodendron sp.* | | | | | AY095458.1 |
| Canellales | Canellaceae | *Canella winterana* | AY572265.1 | AF528847.1 | DQ882240.1 | AF206879.1 | AY095455.1 |

| | | | | | | | |
|---|---|---|---|---|---|---|---|
| Canellales | Winteraceae | *Takhtajania perrieri* | AF206824.1 | AY572287.1 | DQ401371.1 | AF207032.1 | DQ008645.1 |
| Canellales | Winteraceae | *Tasmannia insipida* | L01957.2 | AF093424.1 | | AF207035.1 | AY095469.1 |
| Canellales | Winteraceae | *Tasmannia lanceolata* | | | DQ882241.1 | | |
| Canellales | Winteraceae | *Drymis winteri* | L01905.2 | AF093425.1 | | U42823.1 | AF036491.1 |
| Canellales | Winteraceae | *Drymis granadensis* | | | DQ887676.1 | | |
| Piperales | Aristolochiaceae | *Saruma henryi* | 2924746 | 33327768 | 111154467 | 472412 | 66969232 |
| Piperales | Aristolochiaceae | *Asarum canadense* | L14290.1 | U86383.1 | | | DQ008643.1 |
| Piperales | Aristolochiaceae | *Asarum yakusimense* | | | DQ882197.1 | | |
| Piperales | Aristolochiaceae | *Asarum tamaensis* | | | | DQ472350.1 | |
| Piperales | Aristolochiaceae | *Lactoris fernandeziana* | L08763.1 | AJ235515.2 | DQ882195.1 | U42783.1 | AY095463.1 |
| Piperales | Aristolochiaceae | *Thottea tomentosa* | AF197598.1 | AF197609.1 | AB060738.1 | DQ007406.1 | DQ008642.1 |
| Piperales | Aristolochiaceae | *Thottea corymbosa* | | | 111154473 | | |
| Piperales | Aristolochiaceae | *Aristolochia macrophylla* | L12630.2 | AF528845.1 | AB060742.1 | AF206855.1 | AY095450.1 |
| Piperales | Aristolochiaceae | *Aristolochia manshuriensis* | | | 111154481 | | |
| Piperales | Piperaceae | *Piper arborescens* | AY572253.1 | | | | |
| Piperales | Piperaceae | *Piper capense* | 125991626 | | | | |
| Piperales | Piperaceae | *Piper betle* | | AF528856.1 | | AF206992.1 | AY095467.1 |
| Piperales | Piperaceae | *Piper nigrum* | | | AB040153.2 | | |
| Piperales | Piperaceae | *Peperomia caliginigaudes* | AY572269.1 | AY572291.1 | | AY572313.1 | |
| Piperales | Piperaceae | *Peperomia sp.* | 7105714 | | | | |
| Piperales | Piperaceae | *Peperomia graveolens* | | | DQ212722.1 | | |
| Piperales | Piperaceae | *Peperomia incana* | | | 78155761 | | |
| Piperales | Piperaceae | *Peperomia obtusifolia* | | | | | DQ008641.2 |
| Piperales | Saururaceae | *Houttuynia cordata* | AB205610.1 | AY572283.1, D89555 | AF543737.1 | AF206929.1 | DQ008640.1 |
| Piperales | Saururaceae | *Saururus cernuus* | L14294.1 | AF187061.1 | AF543749.1 | | AY095468.1 |
| Piperales | Saururaceae | *Saururus chinensis* | | | | AY572303.1 | |
| Piperales | Saururaceae | *Anemopsis californica* | AF197597 | AF197608.1 | DQ882198.1 | AF197576.1 | DQ008639.1 |
| Magnoliales | Magnoliaceae | *Magnolia tripetala* | AF206791.1 | AJ235526.1 | | AF206956.1 | |
| Magnoliales | Magnoliaceae | *Magnolia denudata* | | | AF123465.1 | | AF479244.1 |
| Magnoliales | Magnoliaceae | *Liriodendron chinense* | L12654.1 | | AF123481.1 | AJ235981.1 | AY095464.1 |
| Magnoliales | Magnoliaceae | *Liriodendron tulipifera* | | AJ235522.1 | | | |
| Magnoliales | Degeneriaceae | *Degeneria vitensis* | L12643.1 | AJ235451.1 | AB055549.1 | AF206898.1 | DQ008637.1 |
| Magnoliales | Myristicaceae | *Myristica fragrans* | AF206798 | AJ235539 | AJ966803 | AF206968 | |
| Magnoliales | Myristicaceae | *Myristica maingayi* | | | 32483787 | | |
| Magnoliales | Myristicaceae | *Mauloutchia chapelieri* | AF197594 | AF197606 | AY220451 | DQ007409 | DQ008638 |
| Magnoliales | Himantandraceae | *Galbulimima belgraveana* | L12646.2 | AJ235478.1 | AY220441.1 | AF206916.1 | AY095459.1 |
| Magnoliales | Eupomatiaceae | *Eupomatia bennettii* | L12644.2 | AJ235473.1 | DQ401341.1 | AF469771.1 | DQ008636.1 |
| Magnoliales | Annonaceae | *Cananga odorata* | AY841602.1 | DQ007418.1 | AY220438.1 | AF469770.1 | DQ008635.2 |
| Magnoliales | Annonaceae | *Asimina triloba* | AY743441 | AF209532.1 | AF543725.1 | AF206850.1 | AY095451.1 |
| Magnoliales | Annonaceae | *Annona muricata* | AY743440.1 | AJ235393.2 | AF543722.1 | AF206850.1 | DQ008634.2 |

| | | | | | | | |
|---|---|---|---|---|---|---|---|
| Laurales | Calycanthaceae | *Idiospermum australiense* | L12651.2 | AJ235500.1 | AY525342.1 | AF206937.1 | DQ008633.1 |
| Laurales | Calycanthaceae | *Chimonanthus praecox* | L12639.2 | D89558.1 | AY525340.1 | AF503352.1 | DQ008632.1 |
| Laurales | Calycanthaceae | *Calycanthus floridus* | L14291.1 | AJ235422.1 | AF543730.1 | U38318.1 | |
| Laurales | Calycanthaceae | *Calycanthus occidentalis* | | | | | AY095454.1 |
| Laurales | Siparunaceae | *Siparuna glycycarpa* | AF129016.1 | | | | |
| Laurales | Siparunaceae | *Siparuna brasiliensis* | | DQ007421.1 | DQ401375.1 | | |
| Laurales | Siparunaceae | *Siparuna dicipiens* | | | | DQ007411.1 | |
| Laurales | Siparunaceae | *Siparuna decipines* | | | | | DQ008631.1 |
| Laurales | Gomortegaceae | *Gomortega keule* | AF206773.1 | D89560.1 | | AF206918.1 | AY095460.1 |
| Laurales | Atherospermataceae | *Doryphora aromatica* | L77211.2 | | | | |
| Laurales | Atherospermataceae | *Doryphora sassafras* | | AF293858.1 | AF542568.1 | AF293754.1 | DQ008630.1 |
| Laurales | Atherospermataceae | *Atherosperma moschatum* | 4416440 | 6467921 | 89242567 | 6467908 | 66969216 |
| Laurales | Atherospermataceae | *Daphnandra repandula* | 4176744 | | | | |
| Laurales | Atherospermataceae | *Daphnandra micrantha* | | 15705375 | 89242569 | 6467909 | 66969217 |
| Laurales | Lauraceae | *Cryptocarya obovata* | L28950.1 | | | | |
| Laurales | Lauraceae | *Cryptocarya meissneriana* | | AF197602.1 | | AF293757.1 | DQ008627.1 |
| Laurales | Lauraceae | *Cryptocarya alba* | | | AJ247158.1 | | |
| Laurales | Lauraceae | *Cryptocarya subtriplinervia* | | | 59932887 | | |
| Laurales | Lauraceae | *Cinnamomum camphora* | L12641.2 | AJ235436.1 | AJ247154.1 | AF206888.1 | DQ008625.1 |
| Laurales | Lauraceae | *Sassafras albidum* | AF206819.1 | AF209668.1 | | U52031.1 | AF264140.1 |
| Laurales | Lauraceae | *Sassafras tzumu* | | | AF244391.1 | | |
| Laurales | Lauraceae | *Laurus nobilis* | AY337731.1 | AJ235518.1 | AF244407.1 | AF197580.1 | DQ008626.1 |
| Laurales | Lauraceae | *Laurus azorica* | | | 60495411 | | |
| Laurales | Hernandiaceae | *Hernandia ovigera* | L12650.2 | DQ007419.1 | | DQ007407.1 | |
| Laurales | Hernandiaceae | *Hernandia nymphaeifolia* | | | AJ247165.2 | | AY095462.1 |
| Laurales | Hernandiaceae | *Gyrocarpus sp.* | L12647.2 | | | | |
| Laurales | Hernandiaceae | *Gyrocarpus americanus* | | AJ235487.1 | DQ401370.1 | AF206923.1 | DQ008624.1 |
| Laurales | Monimiaceae | *Peumus boldus* | AF206807.1 | AF209650.1 | AJ247183.2 | AF206988.1 | AY095466.1 |
| Laurales | Monimiaceae | *Hortonia floribunda* | AF040663.1 | DQ007420.1 | AY437811.1 | DQ007408.1 | AF264143.1 |
| Laurales | Monimiaceae | *Hedycarya arborea* | L12648.2 | AJ235490.1 | AM396509.1 | AF206924.1 | DQ008623.2 |
| Acorales | Acoraceae | *Acorus gramineus* | D28866.1 | AF197616.1 | DQ182341.1 | AF197584.1 | AF036490.1 |
| Alismatales | Araceae | *Orontium aquaticum* | AJ005632.1 | AF197610.1 | AF543744.1 | AF293753.1 | DQ008652.2 |
| Alismatales | Araceae | *Lemna minor* | 209417450 | NC_10109.1 | | | |
| Alismatales | Araceae | *Lemna aequinoctialis* | | | 17220918 | | |
| Alismatales | Araceae | *Xanthosoma sagittifolium* | 7240497 | | | | |
| Alismatales | Araceae | *Xanthosoma mafaffa* | | 89242541 | | | |
| Alismatales | Araceae | *Xanthosoma helleborifolium* | | | 209417770 | | |

| | | | | | | | |
|---|---|---|---|---|---|---|---|
| Alismatales | Araceae | *Spathiphyllum wallisii* | AJ235807.1 | AJ235606.2 | 209417664 | AF207023.1 | AY095473.1 |
| Alismatales | Araceae | *Spathiphyllum floribundum* | | | AF542575.1 | | |
| Alismatales | Tofieldiaceae | *Tofieldia calyculata* | AB183410.1 | AJ235627.2 | AB183403.1 | AF207043.1 | DQ008653.1 |
| Alismatales | Tofieldiaceae | *Pleea tenuifolia* | AJ131774.1 | AJ235564.2 | AB183407.1 | AF206995.1 | AY095472.1 |
| Alismatales | Juncaginaceae | *Triglochin maritima* | AB088811.1 | AF197601.1 | AF542566.1, 47604605 | AF197586.1 | DQ008650.2 |
| Alismatales | Potamogetonaceae | *Potamogeton distinctus* | AB088809.1 | | AB002581.1 | | |
| Alismatales | Potamogetonaceae | *Potamogeton berchtoldii* | | AF197600.1 | | DQ007410.1 | DQ008649.1 |
| Alismatales | Alismataceae | *Alisma plantago-aquatica* | 166423 | DQ007417 | | 6467915 | 66969239 |
| Alismatales | Alismataceae | *Alisma canaliculatum* | | | 10862963 | | |
| Alismatales | Hydrocharitaceae | *Najas guadalupensis* | DQ859169.1 | | | | |
| Alismatales | Hydrocharitaceae | *Najas gracilima* | | DQ401333.1 | | | |
| Alismatales | Hydrocharitaceae | *Najas minor* | | | HM240480.1 | EF526347.1 | EF526396.1 |
| Alismatales | Hydrocharitaceae | *Hydrocharis dubia* | AB004892.1 | | AB002572.1 | AY952398.1 | |
| Alismatales | Hydrocharitaceae | *Hydrocharis morsus-ranae* | | DQ401334.1 | | | |
| Petrosaviales | Petrosaviaceae | *Japonolirion osense* | AB088835 | AY147598 | AB040161 | AF206942 | |
| Petrosaviales | Petrosaviaceae | *Petrosavia sakuraii* | AB088839 | | AB040156 | | |
| Petrosaviales | Petrosaviaceae | *Petrosavia stellaris* | | AF209649 | | AF206987 | |
| Dioscoreales | Nartheciaceae | *Metanarthecium luteoviride* | AB088837.1 | AF308041.1 | AB040163.1 | AF309410.1 | |
| Dioscoreales | Taccaceae | *Tacca chantieri* | AJ235810.1 | AF308025.1 | | | AY095474.1 |
| Dioscoreales | Taccaceae | *Tacca sp.* | | | AB088792.1 | | |
| Dioscoreales | Taccaceae | *Tacca leontopetaloides* | | | | EU420999.1 | |
| Dioscoreales | Dioscoreaceae | *Dioscorea maciba* | AY904800.1 | | | | |
| Dioscoreales | Dioscoreaceae | *Dioscorea mcvaughii* | | AF308004.1 | | | |
| Dioscoreales | Dioscoreaceae | *Dioscorea alata* | | | AB040208.1 | | |
| Dioscoreales | Dioscoreaceae | *Dioscorea schimperiana* | | | | AF309393.1 | |
| Dioscoreales | Dioscoreaceae | *Dioscorea macrostachya* | | | | | AF205123.1 |
| Dioscoreales | Burmanniaceae | *Burmannia biflora* | AF206742.1 | AF209548.1 | | AF168827.1 | |
| Dioscoreales | Burmanniaceae | *Burmannia madagascariensis* | | | AY956485.1 | | AF290589.1 |
| Pandanales | Stemonaceae | *Croomia pauciflora* | AY298827.1 | | AY437815.1 | AF168835.1 | DQ008647.1 |
| Pandanales | Stemonaceae | *Croomia japonica* | 17224679 | AF308039.1 | | | |
| Pandanales | Cyclanthaceae | *Carludovica palmata* | AF197596.1 | AY465545.1 | AB088793.1 | AF293756.1 | DQ008648.1 |
| Pandanales | Velloziaceae | *Acanthochlamys bracteata* | HQ845619.1 | | AY952413.1 | AY952411.1 | |
| Pandanales | Velloziaceae | *Xerophyta retinervis* | EU213532.1 | JN017040.1 | EU214302.1 | | AF205878.1 |
| Pandanales | Velloziaceae | *Xerophyta humilis* | | | | EF418586.1 | |
| Pandanales | Velloziaceae | *Barbacenia elegans* | AJ131946.1 | AJ235406.2 | | AF206861.1 | |
| Liliales | Campynemataceae | *Campynema linearis* | Z77264 | AJ417573 | JN417414.1 | GQ497570.1 | AF364029 |
| Liliales | Smilacaceae | *Smilax glauca* | AF206822.1 | AF209677.1 | | 1280407 | |
| Liliales | Smilacaceae | *Smilax china* | | | AB040204.1 | | |
| Liliales | Smilacaceae | *Smilax bonanox* | | | | | AF293852.1 |
| Liliales | Liliaceae | *Lilium superbum* | L12682.2 | AF209618.1 | | AF206952.1 | |
| Liliales | Liliaceae | *Lilium alexandrae* | | | AB030849.1 | | |
| Liliales | Liliaceae | *Lilium michauxii* | | | | | AF205126.1 |

| | | | | | | | |
|---|---|---|---|---|---|---|---|
| Liliales | Philesiaceae | *Philesia buxifolia* | Z77302.1 | | AY624479.1 | | |
| Liliales | Philesiaceae | *Philesia magellanica* | | AY465551.1 | | | |
| Liliales | Philesiaceae | *Lapageria rosea* | Z77301 | AJ235517 | AY624480 | X63304 | |
| Liliales | Melanthiaceae | *Trillium grandiflorum* | D28164.1 | | | | |
| Liliales | Melanthiaceae | *Trillium erectum* | | AJ417585.1 | | AF207048.1 | |
| Liliales | Melanthiaceae | *Trillium viridescens* | | | AB017416.1 | | |
| Liliales | Melanthiaceae | *Trillium gracilie* | | | | | AF205128.1 |
| Liliales | Colchicaceae | *Colchicum speciosum* | L12673.2 | AF209569.1 | AB040181.2 | | |
| Liliales | Colchicaceae | *Colchicum autumnale* | | | | U42072.1 | AF331971.1 |
| Liliales | Alstroemeriaceae | *Bomarea hirtella* | Z77255 | AJ235413 | | AF206871 | |
| Liliales | Alstroemeriaceae | *Bomarea pauciflora* | | | EU159966.1 | | |
| Liliales | Alstroemeriaceae | *Alstroemeria aurea* | AY120359 | AY465546 | | | |
| Liliales | Alstroemeriaceae | *Alstroemeria sp.* | | | AY624481 | | |
| Liliales | Alstroemeriaceae | *Alstroemeria pulchra* | | | | | AF290586 |
| Asparagales | Asparagales | *Cypripedium calceolus* | AB176549.1 | AJ235448.2 | AY557208.1 | AF069208.1 | |
| Asparagales | Asparagales | *Cypripedium kentuckiensis* | | | | | AF205119.1 |
| Asparagales | Asparagales | *Phalaenopsis equestris* | 3560803 | | | | |
| Asparagales | Asparagales | *Phalaenopsis aphrodite* | | NC_007499.1 | | | |
| Asparagales | Asparagales | *Phalaenopsis chibae* | | | 76667833 | | |
| Asparagales | Asparagales | *Oncidium excavatum* | 3560783 | 8452707 | | | |
| Asparagales | Asparagales | *Oncidium durangense* | | | 223370962 | | |
| Asparagales | Asparagales | *Oncidium ornithoglossum* | | | | 6273824 | |
| Asparagales | Boryaceae | *Borya septentrionalis* | AF206741 | AF209543 | HM640651 | AF206872 | |
| Asparagales | Blandfordiaceae | *Blandfordia punicea* | 1497826 | 8439256 | 71167184 | 7595389 | |
| Asparagales | Hypoxidaceae | *Molineria capitulata* | HM640538.1 | AF168901.1 | AB088783.1 | HM640770.1 | |
| Asparagales | Hypoxidaceae | *Rhodohypoxis millioides* | Z77280.1 | AJ235582.2 | AY368377.1 | AF207008.1 | |
| Asparagales | Hypoxidaceae | *Hypoxis hemerocallidea* | HM640539.1 | | HM640657.1 | HM640771.1 | |
| Asparagales | Hypoxidaceae | *Hypoxis hirsuta* | | | | | AF290585.1 |
| Asparagales | Tecophilaeaceae | *Tecophilaea cyanocrocus* | HM640544.1 | AJ235620.2 | HM640661.1 | AF207036.1 | |
| Asparagales | Ixioliriaceae | *Ixiolirion tataricum* | HM640543.1 | AY147621.1 | AJ579965.1 | HM640775.1 | |
| Asparagales | Iridaceae | *Alophia silvestris* | JQ670512.1 | | | | |
| Asparagales | Iridaceae | *Alophia veracruzana* | | | AJ579931.1 | | |
| Asparagales | Iridaceae | *Alophia drummondii* | | | | | AF203685.1 |
| Asparagales | Iridaceae | *Iris missouriensis* | 37722359 | 37720992 | | | |
| Asparagales | Iridaceae | *Iris colchica* | | | 58219817 | | |
| Asparagales | Iridaceae | *Iris tenax* | | | | JQ283937.1 | JQ283879.1 |
| Asparagales | Iridaceae | *Aristea glauca* | Z77282.1 | AF209531.1 | AJ579933.1 | AF206854.1 | |
| Asparagales | Iridaceae | *Gladiolus buckerveldii* | AF206772.1 | AF209592.1 | HQ394274.1 | L54062.1 | |
| Asparagales | Xanthorrhoeaceae | *Xanthorrhoea resinosa* | 37722369 | 37721004 | | | |

| | | | | | | | |
|---|---|---|---|---|---|---|---|
| Asparagales | Xanthorrhoeaceae | *Xanthorrhoea quadrangulata* | | | 89242571 | 2795854 | |
| Asparagales | Asphodelaceae | *Bulbine suculenta* | Z73684.1 | AJ235421.1 | | AF206876.1 | AY095471.1 |
| Asparagales | Asphodelaceae | *Bulbine frutescens* | | | AJ511414.1 | | |
| Asparagales | Amaryllidaceae | *Crinum asiaticum* | HM640488.1 | JQ273605.1 | | HM640718.1 | |
| Asparagales | Amaryllidaceae | *Crinum moorei* | | | AB017279.1 | | |
| Asparagales | Amaryllidaceae | *Crinum americanum* | | | | | AF293854.1 |
| Asparagales | Amaryllidaceae | *Nothoscordum bivalve* | Z69202.1 | | | | AF293853.1 |
| Asparagales | Amaryllidaceae | *Nothoscordum dialystemon* | | AJ235504.2 | JQ435522.1 | | |
| Asparagales | Amaryllidaceae | *Allium fistulosum* | 124484484 | | | | |
| Asparagales | Amaryllidaceae | *Allium textile* | | 37721008 | | | |
| Asparagales | Amaryllidaceae | *Allium grayi* | | | 5832656 | | |
| Asparagales | Amaryllidaceae | *Allium thunbergii* | | | | AF168825 | |
| Asparagales | Asparagaceae | *Asparagus officinalis* | L05028.2 | AJ235400.2 | | | DQ008646.1 |
| Asparagales | Asparagaceae | *Asparagus cochinchinensis* | | | AB029804.1 | | |
| Asparagales | Asparagaceae | *Asparagus falcatus* | | | | AF069205.1 | |
| Asparagales | Asparagaceae | *Agave ghiesbreghtii* | 17135864 | 14717923 | | 7595361 | |
| Asparagales | Asparagaceae | *Agave attenuata* | | | 89242609 | | |
| Asparagales | Asparagaceae | *Yucca glauca* | 37722389 | | | | |
| Asparagales | Asparagaceae | *Yucca schidigera* | | 69214423 | | | |
| Asparagales | Asparagaceae | *Yucca filamentosa* | | | 62511859 | 61741965 | |
| Asparagales | Asparagaceae | *Lomandra longifolia* | 7240316 | | | | |
| Asparagales | Asparagaceae | *Lomandra ordii* | | 16943762 | | | |
| Asparagales | Asparagaceae | *Lomandra obliqua* | | | 89242593 | | |
| Asparagales | Asparagaceae | *Lomandra hastilis* | | | | HM640750.1 | |
| Asparagales | Asparagaceae | *Nolina recurvata* | 7240393 | 14718150 | 5731683 | HM640703.1 | |
| unplaced Dasypogonaceae | Dasypogonaceae | *Dasypogon bromeliifolius* | AF206758.1 | AF168907.1 | AM114719.1 | AJ417898.1 | AF466384.1 |
| unplaced Dasypogonaceae | Dasypogonaceae | *Calectasia cyanea* | AJ286557.1 | | DQ888765.1 | | AY079521.1 |
| unplaced Dasypogonaceae | Dasypogonaceae | *Calectasia intermedia* | | AF168891.1 | | AF069209.1 | |
| Arecales | Arecaceae | *Chamaedorea seifrizii* | 17135901 | 7582347 | | 3342023 | |
| Arecales | Arecaceae | *Chamaedorea oblongata* | | | 77819970 | | |
| Arecales | Arecaceae | *Elaeis guineensis* | | 192910867 | 90967848 | | |
| Arecales | Arecaceae | *Elaeis oleifera* | | | | 12584295 | |
| Arecales | Arecaceae | *Caryota mitis* | M81811 | AF233082 | AY952414 | AF168831 | |
| Arecales | Arecaceae | *Trachycarpus fortunei* | AJ404752 | AY012403 | HQ619794 | AY012346 | AF290588 |
| Arecales | Arecaceae | *Phoenix dactylifera* | AM110195 | AY012411 | AB040211.1 | AY012354 | |
| Commelinales | Commelinaceae | *Tradescantia spathacea* | 90968377 | | 90967931 | | |
| Commelinales | Commelinaceae | *Tradescantia ohiensis* | | AF168950.1 | | AF069213.1 | |
| Commelinales | Philydraceae | *Philydrum lanuginosum* | U41596.2 | AY147607.1 | AY952429.1 | AY952390.1 | AF205519.1 |
| Commelinales | Pontederiaceae | *Pontederia cordata* | U41593.1 | AF209657.1 | AF434872.1 | AF206998.1 | AF205517.1 |
| Commelinales | Haemodoraceae | *Anigozanthos flavidus* | AJ404843.1 | AF387600.1 | AB088796.1 | AF069214.1 | AF290587.1 |
| Zingiberales | Strelitziaceae | *Strelitzia nicolai* | 18026911 | | | 3342042 | |

| | | | | | | | |
|---|---|---|---|---|---|---|---|
| Zingiberales | Strelitziaceae | *Strelitzia reginae* | | 42559029 | | | |
| Zingiberales | Strelitziaceae | *Strelitzia alba* | | | 33314647 | | |
| Zingiberales | Strelitziaceae | *Ravenala madagascariensis* | L05459 | AF168939 | AF434873 | AF069228 | |
| Zingiberales | Strelitziaceae | *Phenakospermum guyannense* | AF243845 | AF168938 | AF478911 | AF069227 | |
| Zingiberales | Musaceae | *Musa acuminata* | L05455.1 | AF168931.1 | | AF069226.1 | EU418632.1 |
| Zingiberales | Musaceae | *Musa beccarii* | | | AF434869.1 | | |
| Zingiberales | Musaceae | *Musa rosea* | | | 90967929 | | |
| Zingiberales | Zingiberaceae | *Zingiber gramineum* | L05465.1 | AF168953.1 | AB088799.1 | U42081.1 | |
| Zingiberales | Zingiberaceae | *Zingiber officinale* | | | | | AF205522.1 |
| Zingiberales | Marantaceae | *Maranta bicolor* | AF378768.1 | AF168927.1 | AY140302.1 | AF069225.1 | AY673056.1 |
| Zingiberales | Cannaceae | *Canna indica* | AF378763.1 | AF168892.1 | AM114724.1 | D29784.1 | |
| Zingiberales | Cannaceae | *Canna flaccida* | | | | | AF205521.1 |
| Poales | Typhaceae | *Sparganium americanum* | 343405 | | | | |
| Poales | Typhaceae | *Sparganium eurycarpum* | | 42559027 | | 532650 | |
| Poales | Typhaceae | *Sparganium glomeratum* | | | 62511851 | | |
| Poales | Typhaceae | *Typha latifolia* | DQ069503.1 | DQ069347.1 | DQ069587.1 | AF168880.1 | |
| Poales | Typhaceae | *Typha domingensis* | | | | | AY079520.1 |
| Poales | Bromeliaceae | *Vriesea psittacina* | 53831785 | | | | |
| Poales | Bromeliaceae | *Vriesea splendens* | | 89242555 | | | |
| Poales | Bromeliaceae | *Vriesea monstrum* | | | 53831317 | | |
| Poales | Bromeliaceae | *Puya raimondii* | AF206814 | AF209661 | FJ968207 | AF207001 | |
| Poales | Rapateaceae | *Stegolepis ligulata* | 53831743 | | 53831255 | | |
| Poales | Rapateaceae | *Stegolepis sp.* | | 89242553 | | | |
| Poales | Xyridaceae | *Xyris jupicai* | AY465698 | AY465541 | | | |
| Poales | Xyridaceae | *Xyris difformis* | | | | AF168881 | |
| Poales | Xyridaceae | *Xyris laxifolia* | | | | | AF466386 |
| Poales | Eriocaulaceae | *Eriocaulon microcephalus* | L10252.2 | | | | |
| Poales | Eriocaulaceae | *Eriocaulon compressum* | | EU832854.1 | | | |
| Poales | Eriocaulaceae | *Eriocaulon septangulare* | | | AY952430.1 | AY952402.1 | |
| Poales | Eriocaulaceae | *Eriocaulon decangulare* | | | | | AY079519.1 |
| Poales | Mayacaceae | *Mayaca fluviatilis* | AF036885.1 | AJ419148.1, AF168929.1 | | | AF293855.1 |
| Poales | Mayacaceae | *Mayaca aubletti* | | | | AF168859.1 | |
| Poales | Juncaceae | *Juncus effusus* | AY216612.1 | AJ235509.2 | AB088803.1 | AF206944.1 | AF205520.1 |
| Poales | Cyperaceae | *Rhynchospora nervosa* | Y12977.1 | AF209667.1 | | AF207009.1 | |
| Poales | Cyperaceae | *Rhynchospora alba* | | | JN894695.1 | | |
| Poales | Cyperaceae | *Rhynchospora latifolia* | | | | | AF205518.1 |
| Poales | Cyperaceae | *Cyperus alternifolius* | AM999802.1 | | AY952421.1 | AY952404.1 | |
| Poales | Cyperaceae | *Cyperus albostriatus* | | AF168906.1 | | | |
| Poales | Anarthriaceae | *Anarthria polyphylla* | AF148760.1 | | DQ257498.2 | | |
| Poales | Anarthriaceae | *Anarthria scabra* | | AJ419129.1 | | | |
| Poales | Anarthriaceae | *Anarthria humilis* | | | | | AF466389.1 |
| Poales | Restionaceae | *Restio tetraphyllus* | AF206816.1 | | AF164379.1 | AF207006.1 | AF486829.1 |
| Poales | Restionaceae | *Restio paludosus* | | AJ419152.1 | | | |
| Poales | Centrolepidaceae | *Centrolepis strigosa* | AJ286558.1 | AJ419131.1 | DQ257507.2 | | AF466388.1 |
| Poales | Flagellariaceae | *Flagellaria indica* | AF206769 | AJ419141 | AB040214 | AF206913 | |
| Poales | Joinvilleaceae | *Joinvillea plicata* | EF423010 | EF422979 | | | |

| | | | | | | | |
|---|---|---|---|---|---|---|---|
| Poales | Joinvilleaceae | *Joinvillea ascendens* | | | AF164380 | AF168855 | |
| Poales | Ecdeiocoleaceae | *Ecdeiocolea monostachya* | AF148773 | AJ419136 | DQ257530 | GQ497573.1 | AF466387 |
| Poales | Poaceae | *Oryza sativa* | D00207.1 | AB037543 | EU434287 | AF069218 | |
| Poales | Poaceae | *Oryza longistaminata* | | | | | AY097328.1 |
| Poales | Poaceae | *Hordeum vulgare* | 61378650 | 11582 | | | |
| Poales | Poaceae | *Hordeum murinum* | | | 27544095 | | |
| Poales | Poaceae | *Hordeum bulbosum* | | | | X53792.1 | |
| Poales | Poaceae | *Danthonia californica* | EF423004 | EF422972 | | | |
| Poales | Poaceae | *Danthonia spicata* | | | AF164409 | | |
| Poales | Poaceae | *Danthonia californica* | | | | | EU400689 |
| Poales | Poaceae | *Saccharum officinarum* | 144583483 | NC_006084.1 | 194021757 | 472232 | |
| Poales | Poaceae | *Zea mays* | 18035 | NC_001666.2 | 62868807 | 471990 | 28022.2 |
| Ceratophyllales | Ceratophyllaceae | *Ceratophyllum demersum* | D89473.1 | AJ235430.2 | AF543732.1 | U42517.1 | AY095456.1 |
| Ranunculales | Eupteleaceae | *Euptelea polyandra* | L12645.2 | AF528850.1 | DQ401348.1 | L75831.1 | AF389249.1 |
| Ranunculales | Eupteleaceae | *Euptelea pleiosperma* | | | 121491023 | | |
| Ranunculales | Papaveraceae | *Eschscholzia californica* | 2687476 | 29726129 | GU266597.1 | 115490334 | |
| Ranunculales | Papaveraceae | *Hypecoum imberbe* | 2687482 | 14718079 | GU266596.1 | 6706966 | 22595036 |
| Ranunculales | Papaveraceae | *Dicentra eximia* | L37917.2 | | DQ182345.1 | L37908.1 | AF389262.1 |
| Ranunculales | Papaveraceae | *Dicentra chrysantha* | | AJ235454.2 | | | |
| Ranunculales | Circaeasteraceae | *Kingdonia uniflora* | AF093719.1 | AF092115.1 | FJ626519.1 | AF094537.1 | AF389245.1 |
| Ranunculales | Circaeasteraceae | *Circaeaster agrestis* | FJ626607.1 | AF092116.1 | GU266594.1 | AF094538.1 | AF389246.1 |
| Ranunculales | Lardizabalaceae | *Decaisnea insignis* | 229464427 | | 229464285 | | |
| Ranunculales | Lardizabalaceae | *Decaisnea fargesii* | | 25990326 | | 904097 | 22595027 |
| Ranunculales | Lardizabalaceae | *Sargentodoxa cuneata* | 229464429 | 6017831 | 89242583 | 1479997 | 66969208 |
| Ranunculales | Lardizabalaceae | *Lardizabala biternata* | D85693.1 | L37929.1 | AY437809.1 | L37910.1 | DQ008618.1 |
| Ranunculales | Lardizabalaceae | *Akebia quinata* | L12627.2 | AF209523.1 | AB069851.1 | L37905.1 | AF389253.1 |
| Ranunculales | Menispermaceae | *Menispermum dauricum* | 224830669 | | | | |
| Ranunculales | Menispermaceae | *Menispermum canadense* | | 6017807 | GU266604.1 | 2924752 | 22595030 |
| Ranunculales | Menispermaceae | *Tinospora sinensis* | 229464423 | | 134039086 | | |
| Ranunculales | Menispermaceae | *Tinospora smilacina* | | 224830593 | | | |
| Ranunculales | Menispermaceae | *Tinospora caffra* | | | | 904164 | 22595031 |
| Ranunculales | Menispermaceae | *Cocculus trilobus* | L12642.2 | AF197614.1 | | AF197581.1 | DQ008617.1 |
| Ranunculales | Menispermaceae | *Cocculus laurifolius* | | | AF542588.2 | | |
| Ranunculales | Menispermaceae | *Cissampelos pareira* | AF197590.1 | AF197613.1 | AJ966802.1 | AF293758.1 | DQ008616.1 |
| Ranunculales | Ranunculaceae | *Glaucidium palmatum* | AF093723.1 | AF093375.1 | AB069850.1 | AF094545.1 | AF389267.1 |
| Ranunculales | Ranunculaceae | *Hydrastis canadensis* | L75849.2 | AF093382.1 | AB069849.1 | L75828.1 | AF389268.1 |
| Ranunculales | Ranunculaceae | *Xanthorhiza simplicissima* | L12669.2 | AF093394.1 | AB069848.1 | L75839.1 | AF389270.1 |
| Ranunculales | Ranunculaceae | *Ranunculus acris* | AY395557.1 | | AY954199.1 | | |
| Ranunculales | Ranunculaceae | *Ranunculus macranthus* | | DQ069346.1 | | | |

| | | | | | | | |
|---|---|---|---|---|---|---|---|
| Ranunculales | Ranunculaceae | *Ranunculus sceleratus* | | | | EF526357.1 | |
| Ranunculales | Ranunculaceae | *Ranunculus keniensis* | | | | | AF389269.1 |
| Ranunculales | Berberidaceae | *Mahonia bealei* | L12657.2 | AF197611.1 | | AF293755.1 | DQ008613.1 |
| Ranunculales | Berberidaceae | *Mahonia japonica* | | | AF542585.1 | | |
| Ranunculales | Berberidaceae | *Caulophyllum thalictroides* | AF190442.1 | AF092108.1 | AB069831.1 | L54064.1 | AF389240.1 |
| Ranunculales | Berberidaceae | *Nandina domestica* | DQ923117.1 | DQ923117.1 | AB069830.1 | L37911.1 | AF389241.1 |
| Ranunculales | Berberidaceae | *Podophyllum peltatum* | AF197591.1 | AF092109.1 | AF542586.1 | L24413.1 | DQ008614.1 |
| Sabiales | Sabiaceae | *Sabia swinhoei* | 229464451 | 6017829 | | 1479995 | 22595045 |
| Sabiales | Sabiaceae | *Sabia japonica* | | | 121491027 | | |
| Sabiales | Sabiaceae | *Meliosma veitchiorum* | AF206793.1 | AF209626.1 | AJ581449.1 | AF206961.1 | AF389271.1 |
| Sabiales | Sabiaceae | *Meliosma cuneifolia* | | | 121491029 | | |
| Proteales | Nelumbonaceae | *Nelumbo lutea* | DQ182337.1 | AF528853.1 | | AF094556.1 | AF389259.1 |
| Proteales | Nelumbonaceae | *Nelumbo nucifera* | 229464449 | | AF543740.1 | | |
| Proteales | Platanaceae | *Platanus occidentalis* | AF081073.1 | AF528858.1 | AF543747.1 | U42794.1 | AF274662.1 |
| Proteales | Proteaceae | *Petrophile biloba* | 4098553 | | | | |
| Proteales | Proteaceae | *Petrophile circinata* | | 3850921 | | | |
| Proteales | Proteaceae | *Petrophile canescens* | | | 166156327 | 15428397 | 66969198 |
| Proteales | Proteaceae | *Roupala macrophylla* | 6017866 | | | 6706972 | 22595038 |
| Proteales | Proteaceae | *Roupala montana* | | 194267336 | 166156338 | | |
| Proteales | Proteaceae | *Grevillea robusta* | AF197589.1 | | | AF197577.1 | |
| Proteales | Proteaceae | *Grevillea baileyana* | | AF060434.1 | | | |
| Proteales | Proteaceae | *Grevillea banksii* | | | AF542583.2 | | DQ008612.1 |
| Trochodendrales | Trochodendraceae | *Trochodendron aralioides* | L01958 | AF093423 | GQ998807 | AH001800 | AF479205 |
| Trochodendrales | Trochodendraceae | *Tetracentron sinense* | L12668 | AF093422 | AF274633, 121491011 | U42814 | AF274670 |
| Buxales | Didymelaceae | *Didymeles perrieri* | AF061994.1 | AF092119.1 | DQ401354.1 | AF094541.1 | AF389247.1 |
| Buxales | Buxaceae | *Pachysandra procumbens* | AF061993.1 | | | AF094533.1 | DQ008607.1 |
| Buxales | Buxaceae | *Pachysandra terminalis* | | AF528854.1 | AF542581.2 | | |
| Buxales | Buxaceae | *Buxus sempervirens* | DQ182333.1 | AF092110.1 | AF543728.1 | X16599.1 | AF389243.1 |
| Gunnerales | Gunneraceae | *Gunnera hamiltonii* | AF093724.1 | AF093374.1 | | | |
| Gunnerales | Gunneraceae | *Gunnera perpensa* | | | AY042596.1 | | |
| Gunnerales | Gunneraceae | *Gunnera tinctoria* | | | 121491015 | | |
| Gunnerales | Gunneraceae | *Gunnera manicata* | | | | U43787.1 | AF389250.1 |
| Gunnerales | Myrothamnaceae | *Myrothamnus flabellifolius* | AF060707.1 | AF093386.1 | AM396507.1 | AF094555.1 | AF479223.1 |
| Gunnerales | Myrothamnaceae | *Myrothamnus moschata* | | | AF542591.2 | | |
| Dilleniales | Dilleniaceae | *Dillenia indica* | L01903.2 | | | | |
| Dilleniales | Dilleniaceae | *Dillenia retusa* | | AF095732.1 | | | AF479096.1 |
| Dilleniales | Dilleniaceae | *Hibbertia volubilis* | AF093721 | AF092120 | | AF094542 | |
| Dilleniales | Dilleniaceae | *Hibbertia cuneiformis* | | | HQ896421 | | HQ843451 |
| Dilleniales | Dilleniaceae | *Tetracera asiatica* | AJ235796 | AJ235622 | AY042665 | AJ235982 | AF479097 |
| Santalales | Coulaceae | *Ochanostachys amentacea* | DQ790146.1 | | DQ790183.1 | DQ790116.1 | |
| Santalales | Coulaceae | *Miquartia guianensis* | DQ790148.1 | | DQ790185.1 | L24396.1 | |

| | | | | | | | |
|---|---|---|---|---|---|---|---|
| Santalales | Coulaceae | *Coula edulis* | DQ790147.1 | | DQ790184.1 | | |
| Santalales | Olacaceae | *Heisteria parvifolia* | AJ131771 | AJ235492 | DQ790199 | | DQ790232 |
| Santalales | Olacaceae | *Heisteria concinna* | | | | L24146 | |
| Santalales | Olacaceae | *Ximenia americana* | GQ997898 | GQ997862 | GQ997871 | L24428 | DQ790220 |
| Santalales | Santalaceae | *Santalum album* | L26077.1 | AJ235592.2 | AY042650.1 | L24416.1 | AY957453.1 |
| Santalales | Santalaceae | *Osyris lanceolata* | L11196.2 | AF209641.1 | | L24409.1 | AF389274.1 |
| Santalales | Santalaceae | *Osyris quadripartita* | | | AY042623.1 | | |
| Santalales | Opiliaceae | *Opilia sp.* | AJ131773.1 | AJ235550.2 | AY042621.1 | | |
| Santalales | Opiliaceae | *Opilia amantaceae* | | | | U42790.1 | AF479095.1 |
| Santalales | Loranthaceae | *Gaiadendron punctatum* | 532000 | | 111145144 | 532585 | 112408972 |
| Santalales | Schoepfiaceae | *Schoepfia schreberi* | L11205.2 | AF209671.1 | AY957454.1, 111145148 | AF207017.1, 532648 | AF389261.1 |
| Santalales | Misodendraceae | *Misodendrum linearifolium* | L26074.1 | | DQ787438.1 | L24397.2 | DQ790211.1 |
| Santalales | Misodendraceae | *Misodendrum punctulatum* | | | DQ787443.1 | | |
| Berberidopsidales | Berberidopsidaceae | *Berberidopsis corallina* | AJ235773.1 | AJ235409.2 | EU002171.1 | AF206866.1 | AF389242.1 |
| Berberidopsidales | Aextoxicaceae | *Aextoxicon punctatum* | X83986 | AJ235384.2 | DQ182342.1 | AF206839.1 | AF389239.1 |
| Caryophyllales | Nepenthaceae | *Nepenthes alata* | L01936.2 | AJ235542.2 | | | |
| Caryophyllales | Nepenthaceae | *Nepenthes madagascariensis* | | | AF315883.1 | | |
| Caryophyllales | Nepenthaceae | *Nepenthes sp.* | | | | U42787.1 | AF389260.1 |
| Caryophyllales | Droseraceae | *Drosera capensis* | L01909.2 | AY096110.1 | AY096122 | U42532.1 | AF389248.1 |
| Caryophyllales | Droseraceae | *Drosera regia* | | | AF204848.1 | | |
| Caryophyllales | Drosophyllaceae | *Drosophyllum lusitanicum* | L01907 | AY096113 | AY042580 | AY096119 | HQ843447 |
| Caryophyllales | Ancistrocladaceae | *Ancistrocladus korupensis* | AF206733 | AF209526 | | AF206846 | HQ843441 |
| Caryophyllales | Ancistrocladaceae | *Ancistrocladus abbreviatus* | | | AF315939 | | |
| Caryophyllales | Dioncophyllaceae | *Triphyophyllum peltatum* | Z97637.1 | AF209693.1 | AF315940.1 | AF207049.1 | AF479091.1 |
| Caryophyllales | Frankeniaceae | *Frankenia pulverulenta* | Z97638 | AJ235476 | | AF206914 | HQ843448 |
| Caryophyllales | Frankeniaceae | *Frankenia laevis* | | | AF204862 | | |
| Caryophyllales | Tamaricaceae | *Tamarix pentandra* | Z97650.1 | AF209684.1 | AY042663 | AF207033.1 | AF479083.1 |
| Caryophyllales | Tamaricaceae | *Tamarix gallica* | | | AF204861.1 | | |
| Caryophyllales | Polygonaceae | *Fagopyrum statice* | 3983621 | | | | |
| Caryophyllales | Polygonaceae | *Fagopyrum esculentum* | | EU254477.1 | | | |
| Caryophyllales | Polygonaceae | *Polygonum aviculare* | AF297127.1 | | EF438020 | | |
| Caryophyllales | Polygonaceae | *Polygonum sachalinense* | | AJ235569.2 | | | AF479085.1 |
| Caryophyllales | Polygonaceae | *Polygonum alpinum* | | | AF204858.1 | | |
| Caryophyllales | Polygonaceae | *Polygonum sp.* | | | | AF206996.1 | |
| Caryophyllales | Plumbaginaceae | *Limonium gibertii* | AJ786659 | | | | |
| Caryophyllales | Plumbaginaceae | *Limonium arborescens* | | AF209620 | | AF206953 | HQ843453 |
| Caryophyllales | Plumbaginaceae | *Limonium latifolium* | | | AY514861 | | |
| Caryophyllales | Plumbaginaceae | *Plumbago auriculata* | M77701.1 | | EU002187 | U42795.1 | AF036492.1 |
| Caryophyllales | Plumbaginaceae | *Plumbago zeylanica* | | AJ235565.2 | | | |
| Caryophyllales | Plumbaginaceae | *Plumbago indica* | | | AF204857.1 | | |
| Caryophyllales | Rhabdodendraceae | *Rhabdodendron amazonicum* | Z97649.1 | AJ235578.2 | | AF207007.1 | HQ843460.1 |

| | | | | | | | |
|---|---|---|---|---|---|---|---|
| Caryophyllales | Rhabdodendraceae | *Rhabdodendron macrophyllum* | | | AY042642.1 | | |
| Caryophyllales | Simmondsiaceae | *Simmondsia chinensis* | AF093732 | AF093401 | AF204863, AY042657 | AF094562 | HQ843463.1 |
| Caryophyllales | Physenaceae | *Physena sp.* | Y13116 | | | | |
| Caryophyllales | Physenaceae | *Physena madagascariensis* | | HQ843260 | | HQ843434 | HQ843458 |
| Caryophyllales | Asteropeiaceae | *Asteropeia micraster* | AF206737.1 | AF209533.1 | AY042549.1 | AF206857.1 | AF479090.1 |
| Caryophyllales | Caryophyllaceae | *Stellaria media* | AF206823.1 | AF209680.1 | AY936299 | AF207027.1 | AF479084.1 |
| Caryophyllales | Caryophyllaceae | *Stellaria nemorum* | | | AY936298.1 | | |
| Caryophyllales | Achatocarpaceae | *Phaulothamnus spinescens* | M97887 | HQ843259 | AF542594 | HQ843433 | HQ843457 |
| Caryophyllales | Amaranthaceae | *Spinacia oleracea* | AJ400848 | AF528861 | NC_002202 | L24420 | HQ843464 |
| Caryophyllales | Amaranthaceae | *Beta vulgaris* | DQ074969 | DQ067451 | DQ116790 | | |
| Caryophyllales | Amaranthaceae | *Celosia argentea* | AY270072.1 | AF209559.1 | | AF206883.1 | |
| Caryophyllales | Amaranthaceae | *Celosia trigyna* | | | AY514811.1 | | |
| Caryophyllales | Amaranthaceae | *Celosia cristata* | | | | | HQ843444.1 |
| Caryophyllales | Stegospermataceae | *Stegnosperma halimifolium* | M62571 | | HQ878442 | | HQ843465 |
| Caryophyllales | Limeaceae | *Limeum aethiopicum* | AF132095 | | | | |
| Caryophyllales | Limeaceae | *Limeum sp.* | | AF093385 | | AF094554 | |
| Caryophyllales | Limeaceae | *Limeum africanum* | | | AY042608 | | HQ843452 |
| Caryophyllales | Lophiocarpaceae | *Corbichonia decumbens* | AF132096.1 | GQ497648.1 | FN825760.1 | GQ497577.1 | |
| Caryophyllales | Barbeuiaceae | *Barbeuia madagascarensis* | GQ497673.1 | | 15340899 | | |
| Caryophyllales | Aizoaceae | *Lampranthus filicaulis* | 125857478 | | | | |
| Caryophyllales | Aizoaceae | *Lampranthus blandus* | | | FN597631.1 | | |
| Caryophyllales | Aizoaceae | *Delosperma echinatum* | AJ235778 | AJ235452 | AY042575 | | |
| Caryophyllales | Aizoaceae | *Delosperma napiforme* | | | | HQ843428 | HQ8434461 |
| Caryophyllales | Gisekiaceae | *Gisekia pharnacioides* | M97890 | | | | |
| Caryophyllales | Gisekiaceae | *Gisekia africana* | | | AY042591 | | HQ843449 |
| Caryophyllales | Sarcobataceae | *Sarcobatus vermiculatus* | AF132088 | HQ843262 | AY042652 | GQ497586.1 | HQ843462 |
| Caryophyllales | Phytolaccaceae | *Phytolacca americana* | M62567.1 | AF528855.1 | DQ401362.1 | AF094557.1 | HQ843459.1 |
| Caryophyllales | Phytolaccaceae | *Rivina humilis* | M62569 | HQ843261 | AY042646 | HQ843438 | HQ843461 |
| Caryophyllales | Nyctaginaceae | *Mirabilis jalapa* | M62565.1 | AF209629.1 | FN868307.1 | U42788.1 | HQ843454, AF479086.1 |
| Caryophyllales | Nyctaginaceae | *Bougainvillea glabra* | M88340.1 | AJ235415.2 | | AF206873.1 | |
| Caryophyllales | Nyctaginaceae | *Bougainvillea sp.* | | | AY042560 | | |
| Caryophyllales | Nyctaginaceae | *Bougainvillea spectabilis* | | | | | HQ843443 |
| Caryophyllales | unassigned Hypertellis | *Hypertelis spergulacea* | FN824420.1 | | FN825700.1 | | |
| Caryophyllales | Molluginaceae | *Mollugo verticillata* | M62566.1 | AF209631.1 | AY936330.1 | U42828.1 | AF479088.1 |
| Caryophyllales | Montiaceae | *Claytonia perfoliata* | AF132093 | | | | |
| Caryophyllales | Montiaceae | *Claytonia virginica* | | HQ843256 | | HQ843427 | HQ843445 |
| Caryophyllales | Montiaceae | *Claytonia caroliniana* | | | AY764099 | | |
| Caryophyllales | Talinaceae | *Talinum paniculatum* | GQ436529 | HQ843263 | GQ434150 | HQ843439 | HQ843466 |
| Caryophyllales | Portulacaceae | *Portulaca grandiflora* | M62568.1 | AF209659.1 | | AF207000.1 | AF479093.1 |

| | | | | | | | |
|---|---|---|---|---|---|---|---|
| Caryophyllales | Portulacaceae | *Portulaca oleracea* | | | AF204867.1, DQ855850 | | |
| Caryophyllales | Cactaceae | *Opuntia dillenii* | AY875233 | | | | |
| Caryophyllales | Cactaceae | *Opuntia microdasys* | | HQ843258 | | HQ843432 | HQ843456 |
| Caryophyllales | Cactaceae | *Opuntia quimilo* | | | AY015279 | | |
| Caryophyllales | Cactaceae | *Pereskia aculeata* | AF206805.1 | AF209648.1 | DQ855863.1 | AF206986.1 | AF479092.1 |
| Caryophyllales | Halophytaceae | *Halophytum ameghinoi* | AJ403024 | HQ843257 | AY042599 | HQ843429 | HQ843450 |
| Caryophyllales | Didieraceae | *Alluaudia procera* | 336270 | | | GQ497645.1 | |
| Caryophyllales | Didieraceae | *Alluaudia ascendens* | | | AY042541 | | HQ843440 |
| Caryophyllales | Basellaceae | *Anredera cordifolia* | HM849777.1 | | HM851012.1 | | |
| Caryophyllales | Basellaceae | *Anredera baselloides* | | HQ620741.1 | | | |
| Caryophyllales | Basellaceae | *Anredera brachystachys* | | | FN597626.1 | | |
| Caryophyllales | Basellaceae | *Basella alba* | M62564 | HQ8432541 | AY042553 | HQ843426 | HQ843442 |
| Cornales | Hydrostachyaceae | *Hydrostachys multifida* | U17879.1 | | AB038179.1 | | AY260021.1 |
| Cornales | Hydrostachyaceae | *Hydrostachys imbricata* | | AJ236230.1 | | AJ235983.1 | |
| Cornales | Nyssaceae | *Nyssa ogeche* | L11228.2 | | U96886.1 | U52032.1 | AF297545.1 |
| Cornales | Nyssaceae | *Nyssa sylvatica* | | AJ235545.2 | | | |
| Cornales | Nyssaceae | *Nyssa sp.* | | | 85679045 | | |
| Cornales | Nyssaceae | *Camptotheca acuminata* | L11211.2 | AF209554.1 | U96888.1 | U42789.1 | AY260010.1 |
| Cornales | Cornaceae | *Cornus mas* | L11216.2 | AF528848.1 | AJ429275.1 | | AF297535.1 |
| Cornales | Cornaceae | *Cornus officinalis* | | | | U52033.1 | |
| Cornales | Cornaceae | *Alangium chinense* | L11209.2 | | | AF206843.1 | AY260009.1 |
| Cornales | Cornaceae | *Alangium sp.* | | AJ235386.2 | | | |
| Cornales | Cornaceae | *Alangium platanifolium* | | | U96880.1 | | |
| Cornales | Grubbiaceae | *Grubbia tormentosa* | Z83141.1 | JF298839.1 | | | AY260020.1 |
| Cornales | Grubbiaceae | *Grubbia rosmarinifolia* | | | AJ429276.1 | | |
| Cornales | Curtisiaceae | *Curtisia dentata* | L11222.2 | JF298838.1 | U96901.1 | L16007.1 | AY260012.1 |
| Cornales | Hydrangeaceae | *Fendlera rupicola* | AF206766.1 | AJ236234.1 | AY254063.1 | AJ235986.1 | AY260041.1 |
| Cornales | Hydrangeaceae | *Philadelphus lewisii* | L11198.2 | AJ236231.1 | | U42782.1 | AF389252.1 |
| Cornales | Hydrangeaceae | *Philadelphus hirsutus* | | | U96881.1 | | |
| Cornales | Hydrangeaceae | *Hydrangea macrophylla* | L11187.2 | AF528852.1 | | U42781.1 | |
| Cornales | Hydrangeaceae | *Hydrangea paniculata* | | | AB236029.1 | | |
| Cornales | Hydrangeaceae | *Hydrangea arborescens* | | | | | AY260032.1 |
| Cornales | Loasaceae | *Eucnide lobata* | U17874.1 | | | | |
| Cornales | Loasaceae | *Eucnide bartonoides* | | AJ236227.1 | | AJ235988.1 | |
| Cornales | Loasaceae | *Eucnide urens* | | | AF503315.1 | | AY260031.1 |
| Cornales | Loasaceae | *Mentzelia decapetala* | 643626 | | | | 30230599 |
| Cornales | Loasaceae | *Mentzelia lindleyi* | | 6688707 | | 6688701 | |
| Cornales | Loasaceae | *Mentzelia reflexa* | | | 33867458 | | |
| Cornales | Loasaceae | *Petalonyx nitidus* | AF299086.1 | AJ236232.1 | | AJ235989.1 | AY260028.1 |
| Cornales | Loasaceae | *Petalonyx crenatus* | | | AF503295.1 | | |
| Ericales | Marcgraviaceae | *Marcgravia rectiflora* | Z83148.1 | AJ235529.1 | | | AY727937.1 |
| Ericales | Marcgraviaceae | *Marcgravia sp.* | | | AJ429289.1 | | |
| Ericales | Tetrameristaceae | *Tetramerista sp.* | Z80199.1 | AJ235623.2 | AJ429304.1 | AF207039.1 | AF479153.1 |
| Ericales | Balsaminaceae | *Impatiens repens* | Z80197.1 | AJ235503.2 | | | AF479154.1 |

| | | | | | | | |
|---|---|---|---|---|---|---|---|
| Ericales | Balsaminaceae | *Impatiens capensis* | | | AJ429280.1 | | |
| Ericales | Balsaminaceae | *Impatiens pallida* | | | | L24148.1 | |
| Ericales | Lecythidaceae | *Couroupita guianensis* | Z80181 | AJ236224 | | AJ235993 | AY727950 |
| Ericales | Lecythidaceae | *Barringtonia asiatica* | Z80174.1 | AY725929.1 | DQ924095.1 | | AY727949.1 |
| Ericales | Lecythidaceae | *Barringtonia racemosa* | AF088853.1 | | | AY289647.1 | |
| Ericales | Fouquieriaceae | *Fouquieria fasciculata* | AY725862 | AY725924 | | | AY727940 |
| Ericales | Fouquieriaceae | *Fouquieria splendens* | | | U96903 | L49280 | |
| Ericales | Fouquieriaceae | *Fouquieria diguetii* | | | 22795826 | | |
| Ericales | Polemoniaceae | *Cobaea scandens* | Z83143.1 | AJ235440.2 | L48568.1 | L49277.1 | AY727944.1 |
| Ericales | Polemoniaceae | *Gilia aggregata* | Z83144.1 | | | | |
| Ericales | Polemoniaceae | *Gilia capitata* | | AJ236220.1 | | | AF479155.1 |
| Ericales | Polemoniaceae | *Gilia sinuata* | | | L34198.1 | | |
| Ericales | Polemoniaceae | *Gilia reticulata* | | | | DQ080013.1 | |
| Ericales | Polemoniaceae | *Polemonium reptans* | L11687 | | | | |
| Ericales | Polemoniaceae | *Polemonium pauciflorum* | | AY725925 | | | |
| Ericales | Polemoniaceae | *Polemonium pulcherrimum* | | | 22797432 | | |
| Ericales | Polemoniaceae | *Polemonium californicum* | | | | L49294 | |
| Ericales | Polemoniaceae | *Polemonium caeruleum* | | | | | AY056516 |
| Ericales | Polemoniaceae | *Phlox longifolia* | AF206809.1 | AJ236221.1 | | | |
| Ericales | Polemoniaceae | *Phlox gracilis* | | | L34203.1 | | |
| Ericales | Polemoniaceae | *Phlox hoodii* | | | | L49293.1 | |
| Ericales | Polemoniaceae | *Phlox divaricada* | | | | | AF148281.1 |
| Ericales | Sapotaceae | *Manilkara zapota* | AF213793.1 | AJ235528.1 | DQ924092.1 | L49288.1 | AY727946.1 |
| Ericales | Ebenaceae | *Lissocarpa benthamii* | EU980793.1 | DQ923969.1 | DQ924077.1 | | AY727956.1 |
| Ericales | Ebenaceae | *Euclea crispa* | EU980789.1 | DQ923966.1 | DQ924073.1 | GU476413.1 | |
| Ericales | Ebenaceae | *Diospyros lotus* | EU980703.1 | DQ923924.1 | AB174992.1 | | AY727957.1 |
| Ericales | Ebenaceae | *Diospyros digyna* | | AF213768.1 | | | |
| Ericales | Ebenaceae | *Diospyros cauliflora* | | | DQ924001.1 | | |
| Ericales | Ebenaceae | *Diospyros virginiana* | | | | L49279.1 | |
| Ericales | Primulaceae (Myrsinaceae) | *Anagallis arvensis* | M88343.1 | | | | |
| Ericales | Primulaceae | *Anagallis tenella* | | AJ235390.2 | AJ581446.1 | AF206845.1 | AF479149.1 |
| Ericales | Primulaceae | *Maesa myrsinoides* | Z80203 | | | | |
| Ericales | Primulaceae | *Maesa tenera* | | AF213781 | 22797100 | | |
| Ericales | Primulaceae | *Maesa japonica* | | | | | AY727959 |
| Ericales | Primulaceae | *Clavija domingensis* | AF213818 | | | | |
| Ericales | Primulaceae | *Clavija eggersiana* | | AJ235437 | | | AF479151 |
| Ericales | Primulaceae | *Clavija costaricana* | | | JQ588865.1 | | |
| Ericales | Primulaceae | *Clavija integrifolia* | | | | AJ235998 | |
| Ericales | Primulaceae | *Primula sieboldii* | U96657 | AF213787 | | | |
| Ericales | Primulaceae | *Primula forrestii* | | | 87197564 | | |
| Ericales | Primulaceae | *Primula sp.* | | | | L49295 | |
| Ericales | Primulaceae | *Primula elatior* | | | | | AY727960 |
| Ericales | Primulaceae | *Androsace erecta* | AF395004.1 | | | | |
| Ericales | Primulaceae | *Androsace sp.* | | AF213775.1 | | | |
| Ericales | Primulaceae | *Androsace sempervivoides* | | | AY647535.1 | | |

| | | | | | | | |
|---|---|---|---|---|---|---|---|
| Ericales | Primulaceae | *Androsace spinulifera* | | | | AF206847.1 | AF479150.1 |
| Ericales | Pentaphylacaceae | *Ternstroemia stahlii* | AF206827.1 | AF209687.1 | | AF207038.1 | AY727955.1 |
| Ericales | Pentaphylacaceae | *Ternstroemia longipes* | | | AF380110.1 | | |
| Ericales | Pentaphylacaceae | *Eurya chinensis* | AF089714.1 | | | | |
| Ericales | Pentaphylacaceae | *Eurya sp.* | | AF420969.1 | | | |
| Ericales | Pentaphylacaceae | *Eurya japonica* | | | AF380081.1 | | AY727953.1 |
| Ericales | Pentaphylacaceae | *Eurya emarginata* | | | | AJ235995.1 | |
| Ericales | Theaceae | *Camellia sinensis* | AF380037.1 | AY725933.1 | AF380077.1 | AB120309.1 | AY727975.1 |
| Ericales | Symplocaceae | *Symplocos zizyphoides* | AY725865.1 | AY725934.1 | | | AY727978.1 |
| Ericales | Symplocaceae | *Symplocos wikstroemiifolia* | | | AY336346.1 | | |
| Ericales | Symplocaceae | *Symplocos paniculata* | | | | U43297.1 | |
| Ericales | Diapensiaceae | *Galax aphylla* | Z80184.1 | | | | |
| Ericales | Diapensiaceae | *Galax urceolata* | | AY725936.1 | L48576.1 | L49281.1 | AY727983.1 |
| Ericales | Styracaceae | *Styrax chinensis* | AF396157.1 | | | | |
| Ericales | Styracaceae | *Styrax japonica* | Z80189 | | | | AF479156.1 |
| Ericales | Styracaceae | *Styrax officinalis* | | AF420984.1 | DQ924099.1 | | |
| Ericales | Styracaceae | *Styrax americana* | | | | U43296.1 | |
| Ericales | Styracaceae | *Halesia carolina* | Z80190.1 | DQ923988.1 | DQ924097.1 | | AY727981.1 |
| Ericales | Styracaceae | *Halesia tetraptera* | | | | L49284.1 | |
| Ericales | Sarraceniaceae | *Sarracenia flava* | L01952.2 | AJ235594.2 | | | |
| Ericales | Sarraceniaceae | *Sarracenia purpurea* | | | AJ429296.1 | U42804.1 | AY260044.1 |
| Ericales | Roridulaceae | *Roridula gorgonias* | L01950.2 | AJ236180.1 | AJ429294.1 | AF207010.1 | AY727965.1 |
| Ericales | Actinidiaceae | *Actinidia chinensis* | L01882.2 | AJ235382.2 | U61324.1 | AF419792.1 | |
| Ericales | Actinidiaceae | *Actinidia arguta* | | | | | AY727964.1 |
| Ericales | Clethraceae | *Clethra alnifolia* | L12609.2 | | AJ429281.1 | AF419793.1 | |
| Ericales | Clethraceae | *Clethra arborea* | | AF420965.1 | | | |
| Ericales | Clethraceae | *Clethra cf. ferruginea* | | | | | AY727968.1 |
| Ericales | Cyrillaceae | *Cyrilla racemiflora* | L01900.2 | AJ235449.2 | AJ429282.1 | U43294.1 | AY727969.1 |
| Ericales | Ericaceae | *Enkianthus campanulatus* | L12616.2 | AF420968.1 | U61344.2 | AF419802.1 | AY727970.1 |
| Ericales | Ericaceae | *Arbutus canariensis* | L12597 | | U61345 | | |
| Ericales | Ericaceae | *Arbutus unedo* | | AF266738 | | AF206853 | DQ067894 |
| Ericales | Ericaceae | *Vaccinium uliginosum* | AF421107.1 | AF420987.1 | | | |
| Ericales | Ericaceae | *Vaccinium scoparium* | | | AF419716.1 | | |
| Ericales | Ericaceae | *Vaccinium macrocarpon* | | | | L49297.2 | |
| Ericales | Ericaceae | *Vaccinium myrtillus* | | | | | AY727974.1 |
| Ericales | Ericaceae | *Rhododendron hippophaeoides* | L01949.2 | | | AF419807.1 | |
| Ericales | Ericaceae | *Rhododendron impeditum* | | AY725932.1 | | | AY727973.1 |
| Ericales | Ericaceae | *Rhododendron wadanum* | | | AB012746.1 | | |
| unassigned Icacinaceae | Icacinaceae | *Icacina mannii* | 17135962 | AF209603 | HQ384577 | 7595455 | |
| Garryales | Eucommiaceae | *Eucommia ulmoides* | L01917 | HQ384789 | AJ429317 | HQ384682 | |
| Garryales | Garryaceae | *Garrya elliptica* | L01919.2 | AJ235479.2 | AJ429319.1 | U4250.1 | AF479181.1 |
| Garryales | Garryaceae (Aucubaceae) | *Aucuba japonica* | L11210.2 | AJ235402.2 | AJ429318.1 | U42522.1 | AY727931.1 |
| unassigned Oncothecaceae | Oncothecaceae | *Oncotheca balansae* | AJ131950 | AJ235549 | AJ581439 | AF206976 | |
| unassigned Vahliaceae | Vahliaceae | *Vahlia capensis* | L11208.2 | AJ236217.1 | AJ429316.1 | U42813.1 | AF479182.1 |

| | | | | | | | |
|---|---|---|---|---|---|---|---|
| Solanales | Montiniaceae | *Montinia caryophyllacea* | L11194.2 | AY100852.1 | AJ429359.1 | U42808.1 | AF479175.1 |
| Solanales | Sphenocleaceae | *Sphenoclea zeylanica* | L18798 | HQ384778 | AJ429360 | GQ497585.1 | |
| Solanales | Hydroleaceae | *Hydrolea ovata* | L14293.1 | AJ236184.1 | AJ429356.1 | AJ236014.1 | AF479177.1 |
| Solanales | Convolvulaceae | *Cuscuta subinclusa* | 32468854 | | | | |
| Solanales | Convolvulaceae | *Cuscuta sandwichiana* | | 21633448 | | | |
| Solanales | Convolvulaceae | *Cuscuta japonica* | | | 165910534 | | |
| Solanales | Convolvulaceae | *Cuscuta gronovii* | | | | 532575 | |
| Solanales | Convolvulaceae | *Cuscuta cuspidata* | | | | | AF148273.1 |
| Solanales | Convolvulaceae | *Ipomea alba* | AY100963.1 | AY100754.1 | | | |
| Solanales | Convolvulaceae | *Ipomoea purpurea* | | EU118126 | | | |
| Solanales | Convolvulaceae | *Ipomea batatas* | | | AJ429355.1 | | |
| Solanales | Convolvulaceae | *Ipomea hederacea* | | | | U38310.1 | |
| Solanales | Convolvulaceae | *Ipomea lacunosa* | | | | | AF146016.1 |
| Solanales | Solanaceae | *Petunia hybrida* | X04976.1 | | | | |
| Solanales | Solanaceae | *Petunia axillaris* | | AJ236182.1 | | AJ236020.1 | AF479174.1 |
| Solanales | Solanaceae | *Petunia scheideana* | | | AB262070.1 | | |
| Solanales | Solanaceae | *Solanum dulcamara* | 241993465 | | | | |
| Solanales | Solanaceae | *Solanum chilense* | | 237783986 | | | |
| Solanales | Solanaceae | *Solanum oedipus* | | | 209864945 | | |
| Solanales | Solanaceae | *Solanum petrophilum* | | | | 21360 | |
| Solanales | Solanaceae | *Atropa belladonna* | 237784030 | 237783992 | 156628783 | | |
| Solanales | Solanaceae | *Nolana spathulata* | U08616.1 | | | | |
| Solanales | Solanaceae | *Nolana humifusa* | | AF209638.1 | | AJ236017.1 | AF148272.1 |
| Solanales | Solanaceae | *Nolana albescens* | | | AB036647.1 | | |
| Solanales | Solanaceae | *Nicotiana tabacum* | Z00044.2 | AF035909.1 | | AJ236016.1 | AF479172.1 |
| Solanales | Solanaceae | *Nicotiana acaulis* | | | AB039985.1 | | |
| Gentianales | Rubiaceae | *Luculia gratissima* | 211573234 | HQ384771 | AJ429325 | | |
| Gentianales | Rubiaceae | *Coffea arabica* | X83631 | AJ235441 | DQ401346 | EU650386.1 | EU650384 |
| Gentianales | Rubiaceae | *Galium aparine* | X81091 | | HQ384560 | HQ384689 | |
| Gentianales | Rubiaceae | *Galium septentrionale* | | X81669 | | | |
| Gentianales | Rubiaceae | *Mitchella repens* | AF190440 | AF209630 | HQ593366.1 | U42802 | AF148279 |
| Gentianales | Loganiaceae | *Strychnos nux-vomica* | L14410 | AJ235613 | Z70193 | | |
| Gentianales | Loganiaceae | *Spigelia marilandica* | L14007 | AF209679 | | AJ236024 | |
| Gentianales | Loganiaceae | *Spigelia sp.* | | | EF077198 | | |
| Gentianales | Gelsemiaceae | *Gelsemium sempervirens* | L14397 | AJ236193 | HQ384556 | AJ236025 | |
| Gentianales | Apocynaceae | *Nerium oleander* | AJ002886.1 | AJ236189.1 | Z98173.1 | AF107572.1 | AF479178.1 |
| Gentianales | Gentianaceae | *Gentiana procera* | L14398 | | | | |
| Gentianales | Gentianaceae | *Gentiana saponaria* | | JF298870 | | | JF321128 |
| Gentianales | Gentianaceae | *Gentiana purpurea* | | | AJ429323 | | |
| Gentianales | Gentianaceae | *Gentiana asclepiadea* | | | | HQ448773 | |
| Gentianales | Gentianaceae | *Exacum affine* | L11684 | AJ236195 | FJ014087, 4456077 | AJ236023 | AF479180 |
| Boraginales | Boraginaceae | *Borago officinalis* | L11680.1 | AJ235414.2 | AJ429308.1 | AF107580.1 | AF479179.1 |
| Boraginales | Boraginaceae | *Hydrophyllum capitatum* | HQ384926 | HQ384784 | HQ384574 | HQ384691 | |
| Boraginales | Boraginaceae | *Ehretia acuminata* | GQ997264 | | GQ997226 | HQ384690 | |
| Boraginales | Boraginaceae | *Ehretia cymosa* | | 24940181 | | | |
| Lamiales | Plocospermataceae | *Plocosperma buxifolium* | Z68829 | HQ384756 | AJ429315 | HQ384684.1 | |
| Lamiales | Oleaceae | *Olea europaea* | DQ673304.1 | AJ236163.1 | AJ429335.1 | L49289.1 | AF479171.1 |
| Lamiales | Oleaceae | *Syringa vulgaris* | DQ673303 | HQ384751 | HQ384544 | HQ384685.1 | |

| | | | | | | | |
|---|---|---|---|---|---|---|---|
| Lamiales | Oleaceae | *Jasminum suavissimum* | 7240290 | | | 6688488 | |
| Lamiales | Oleaceae | *Jasminum polyanthum* | | 8452687 | | | |
| Lamiales | Oleaceae | *Jasminum nudiflorum* | | | 32811725 | | |
| Lamiales | Tetrachondraceae | *Polypremum procumbens* | AJ011989.1 | HQ384749 | AJ429351.1 | | |
| Lamiales | unplaced Peltanthera | *Peltanthera floribunda* | HQ384900 | HQ384748 | AJ429330 | | AY423082 |
| Lamiales | Calceolariaceae | *Calceolaria sp.* | HQ384899 | HQ384746 | | | |
| Lamiales | Calceolariaceae | *Calceolaria pennellii* | | | 44845004 | | |
| Lamiales | Calceolariaceae | *Calceolaria integrifolia* | | | | GQ497569.1 | |
| Lamiales | Calceolariaceae | *Calceolaria arachnoidea* | | | | | 41323876 |
| Lamiales | Gesneriaceae | *Titanotrichum oldhamii* | AF206829 | AJ236171 | | AJ236054 | AY423085 |
| Lamiales | Gesneriaceae | *Rhynchoglossum notonianum* | AF170244.1 | AJ236170 | | AJ236052 | AF479164 |
| Lamiales | Gesneriaceae | *Rhynchoglossum gardneri* | | | FN773555 | | |
| Lamiales | Stilbaceae | *Halleria lucida* | AF026828 | HQ384732.1 | | | |
| Lamiales | Stilbaceae | *Halleria tetragona* | | | 57901546 | | |
| Lamiales | Plantaginaceae | *Antirrhinum majus* | L11688 | AJ235395 | AJ235395 | AJ236047 | 41323870 |
| Lamiales | Plantaginaceae | *Veronica arguta* | AY034025.1 | | | | |
| Lamiales | Plantaginaceae | *Veronica anagallis-aquatica* | | AJ236169.1 | | AF207052.1 | AF479169.1 |
| Lamiales | Plantaginaceae | *Veronica arvensis* | | | AF052003.1 | | |
| Lamiales | Plantaginaceae | *Plantago lanceolata* | L36454.1 | AF209656.1 | | AJ236046.1 | |
| Lamiales | Plantaginaceae | *Plantago argentea* | | | AJ429344.1 | | |
| Lamiales | Plantaginaceae | *Plantago virginica* | | | | | AF148277.1 |
| Lamiales | Scrophulariaceae | *Myoporum mauritianum* | L36445.1 | AJ236166.1 | | AJ236036.1 | AF479170.1 |
| Lamiales | Scrophulariaceae | *Myoporum montanum* | | | AF531808.1 | | |
| Lamiales | Scrophulariaceae | *Verbascum thapsus* | L36452.1 | AJ236177.1 | AF052002.1 | AF207051.1 | AF479168.1 |
| Lamiales | Scrophulariaceae | *Scrophularia sp.* | L36449.1 | | | | |
| Lamiales | Scrophulariaceae | *Scrophularia californica* | HQ384892 | AJ236175.1 | | AJ236031.1 | |
| Lamiales | Scrophulariaceae | *Scrophularia peregrina* | | | AF375185.1 | | |
| Lamiales | Scrophulariaceae | *Scrophularia canina* | | | | | AY423080.1 |
| Lamiales | Martyniaceae | *Martynia annua* | HQ384889.1 | HQ384730 | HQ384524 | | |
| Lamiales | Pedaliaceae | *Sesamum indicum* | HQ384882 | HQ384721 | AJ429340 | AJ236041 | |
| Lamiales | Bignoniaceae | *Catalpa sp.* | 336744 | | | | |
| Lamiales | Bignoniaceae | *Catalpa aff. speciosa* | | HQ384724 | HQ384519 | | |
| Lamiales | Bignoniaceae | *Catalpa bignonioides* | | | | AF107579 | |
| Lamiales | Bignoniaceae | *Campsis radicans* | AF102642 | AJ236168 | AF531775 | AF107578 | |
| Lamiales | Byblidaceae | *Byblis liniflora* | L01891.2 | HQ384703 | AJ429354.1 | L15195.2, HQ384686 | |
| Lamiales | Byblidaceae | *Byblis gigantea* | | AJ236181.1 | | | AF479141.1 |
| Lamiales | Lentibulariaceae | *Pinguicula moranensis* | HQ384871 | HQ384704 | HQ384501 | HQ384687 | |
| Lamiales | Lentibulariaceae | *Utricularia biflora* | L13190 | AJ235636 | | | |
| Lamiales | Lentibulariaceae | *Utricularia alpina* | | | AF531822 | | |
| Lamiales | Lentibulariaceae | *Utricularia sp.* | | | | AJ236044 | |
| Lamiales | Acanthaceae | *Thunbergia usambarica* | L12596.1 | | | | |

| | | | | | | | |
|---|---|---|---|---|---|---|---|
| Lamiales | Acanthaceae | *Thunbergia coccinea* | | AJ235625.2 | | | |
| Lamiales | Acanthaceae | *Thunbergia alata* | | | AF531811.1 | AF107569.1 | |
| Lamiales | Acanthaceae | *Acanthus montanus* | L12592 | HQ384715 | HQ384511 | | |
| Lamiales | Acanthaceae | *Acanthus ebracteatus* | | | | AY289642 | |
| Lamiales | Acanthaceae | *Justicia americana* | L14401 | AJ236178 | | AF107568 | AF479165 |
| Lamiales | Acanthaceae | *Justicia arborescens* | | | JQ586382.1 | | |
| Lamiales | Acanthaceae | *Barleria prionitis* | L01886 | AJ236179 | | AF107567 | AF479166 |
| Lamiales | Acanthaceae | *Barleria rotundifolia* | | | JF270654.1 | | |
| Lamiales | Thomandersiaceae | *Thomandersia laurifolia* | AY919280 | HQ384719 | HQ384515 | | |
| Lamiales | Verbenaceae | *Verbena bonariensis* | L14412.1 | | | | |
| Lamiales | Verbenaceae | *Verbena bracteata* | HQ384875 | HQ384711 | HQ384506 | AJ236042.1 | |
| Lamiales | Verbenaceae | *Verbena scabrido-glandulosa* | | AJ235639.2 | | | AF479167.1 |
| Lamiales | Verbenaceae | *Verbena rigida* | | | AJ429353.1 | | |
| Lamiales | Lamiaceae | *Lamium amplexicaule* | AB266225 | AJ236165 | | L49287 | |
| Lamiales | Lamiaceae | *Lamium album* | | | AJ429332 | | |
| Lamiales | Lamiaceae | *Callicarpa dichotoma* | L14393 | AF209551 | | AJ236048 | |
| Lamiales | Lamiaceae | *Callicarpa stapfii* | | | FM163265 | | |
| Lamiales | Paulowniaceae | *Paulownia tomentosa* | L36447.1 | AJ236174.1 | AJ429339.1 | AJ236039.1 | JF321129.1 |
| Lamiales | Orobanchaceae | *Pedicularis foliosa* | HQ384873 | | | | |
| Lamiales | Orobanchaceae | *Pedicularis coronata* | | AF209646.1 | | | |
| Lamiales | Orobanchaceae | *Pedicularis iwatensis* | | | AB280514.1 | | |
| Lamiales | Orobanchaceae | *Pedicularis racemosa* | | | | U59959.1 | |
| Lamiales | Phrymaceae | *Phryma leptostachya* | U28881 | HQ384710 | AJ429341 | HQ384688 | |
| Lamiales | Mazaceae | *Mazus stachydifolius* | EU348860.1 | | | | |
| Lamiales | Mazaceae | *Mazus reptans* | | HQ384705.1 | | | |
| Lamiales | Mazaceae | *Mazus rugosus* | | | FN773547.1 | | |
| Lamiales | Mazaceae | *Mazus pumilus* | | | | GU359050.1 | AF148278.1 |
| Aquifoliales | Aquifoliaceae | *Ilex cornuta* | GQ997347.1 | GQ997300.1 | | | |
| Aquifoliales | Aquifoliaceae | *Ilex opaca* | | | GQ248140.1, EF590403 | AF161010.1 | AF479203.1 |
| Aquifoliales | Phyllonomaceae | *Phyllonoma laticuspis* | L11201 | AJ236216 | | U42546 | |
| Aquifoliales | Phyllonomaceae | *Phyllonoma ruscifolia* | | | AJ429377 | | |
| Aquifoliales | Helwingiaceae | *Helwingia japonica* | GQ436764 | AF209596 | AJ430195 | U42524 | |
| Aquifoliales | Helwingiaceae | *Helwingia sp.* | | | | | GQ983586 |
| Aquifoliales | Stemonuraceae | *Gomphandra javanica* | AJ402954 | GQ983598 | GQ983651 | | |
| Aquifoliales | Stemonuraceae | *Irvingbaileya sp.* | AF156733 | AJ236219 | | AJ235999 | AF479202 |
| Aquifoliales | Cardiopteridaceae | *Citronella suaveolens* | | GQ983606 | GQ983648 | | |
| Aquifoliales | Cardiopteridaceae | *Gonocaryum litorale* | AJ235779 | AJ235484 | GQ983654 | AF206919 | AF479201 |
| Aquifoliales | Cardiopteridaceae | *Cardiopteris quinqueloba* | AJ402936 | | AJ429310 | | |
| Aquifoliales | Cardiopteridaceae | *Cardiopteris sp.* | | GQ983618 | | GQ983562 | GQ983581 |
| Escalloniales | Escalloniaceae | *Polyosma cunninghamii* | AF299091 | AJ419680 | AJ429368 | | |

| | | | | | | | |
|---|---|---|---|---|---|---|---|
| Escalloniales | Escalloniaceae | *Polyosma sp.* | | | | GQ983575 | |
| Escalloniales | Escalloniaceae | *Anopterus macleayanus* | Y10673 | | GQ983639 | | |
| Escalloniales | Escalloniaceae | *Anopterus glandulosus* | | AJ419683 | | | |
| Escalloniales | Escalloniaceae | *Eremosyne pectinata* | L47969 | AJ236215 | AJ429364 | U42807 | |
| Escalloniales | Escalloniaceae | *Escallonia calcottiae* | AJ419693 | | | | |
| Escalloniales | Escalloniaceae | *Escallonia rubra* | | AJ318974 | AJ429365 | | |
| Escalloniales | Escalloniaceae | *Escallonia coquimbensis* | | | | U42544 | GQ983585 |
| Escalloniales | Escalloniaceae | *Valdivia gayana* | 39725410 | 18077606 | GQ983642 | | |
| Escalloniales | Escalloniaceae | *Forgesia racemosa* | AJ575923 | AJ419678 | GQ983661 | | |
| Asterales | Pentaphragmataceae | *Pentaphragma ellipiticum* | L18794 | AJ318980 | AJ429387 | | |
| Asterales | Pentaphragmataceae | *Pentaphragma sp.* | | | | GQ983574 | |
| Asterales | Rousseaceae | *Roussea simplex* | AF084477.1 | AJ235586.2 | AJ429389.1 | U42548.1 | AF479243.1 |
| Asterales | Rousseaceae | *Carpodetus serratus* | 2385356 | 15425561 | 22795876 | GQ983563 | |
| Asterales | Rousseaceae | *Cuttsia viburnea* | 2385361 | 15425565 | GQ983640 | GU476392.1 | |
| Asterales | Rousseaceae | *Abrophyllum ornans* | 1304273 | 15422201 | GQ983653 | | |
| Asterales | Campanulaceae | *Pseudonemacladus oppositifolius* | AJ318992 | EU437600 | GQ983646 | | |
| Asterales | Campanulaceae | *Dialypetalum sp.* | AJ318991 | AJ318972 | GQ983649 | | |
| Asterales | Campanulaceae | *Lobelia sp.* | L01931.2 | | | | |
| Asterales | Campanulaceae | *Lobelia angulata* | | AJ235524.1 | | | |
| Asterales | Campanulaceae | *Lobelia cardinalis* | | | GQ248149.1, 197257900 | | |
| Asterales | Campanulaceae | *Lobelia erinus* | | | | U42785.1 | |
| Asterales | Campanulaceae | *Lobelia puberula* | | | | | AF148276.1 |
| Asterales | Campanulaceae | *Cyphia elata* | EU713371 | | | | |
| Asterales | Campanulaceae | *Cyphia rogersii* | | AJ318970 | | | |
| Asterales | Campanulaceae | *Cyphia decora* | | | GQ983652 | | |
| Asterales | Campanulaceae | *Campanula trachelium* | DQ356118 | AJ235423 | | | |
| Asterales | Campanulaceae | *Trachelium caeruleum* | | | EU713328 | | |
| Asterales | Campanulaceae | *Campanula ramosa* | L13861.1 | AJ235423.2 | | | |
| Asterales | Campanulaceae | *Campanula elatines* | | | AJ430387.1 | | |
| Asterales | Campanulaceae | *Campanula ramulosa* | | | | U42510.1 | |
| Asterales | Campanulaceae | *Campanula trachelium* | | | | | AF479191.1 |
| Asterales | Phellinaceae | *Phelline billardieri* | AJ238346.1 | | | AF206989.1 | |
| Asterales | Phellinaceae | *Phelline comosa* | X69748 | AJ235557.1 | | | AF479188.1 |
| Asterales | Phellinaceae | *Phelline lucida* | | | AJ429388.1 | | |
| Asterales | Argophyllaceae | *Argophyllum sp.* | 1304277 | AJ318965 | 22795870 | | |
| Asterales | Argophyllaceae | *Argophyllum laxum* | | | | | GQ983580 |
| Asterales | Argophyllaceae | *Corokia cotoneaster* | L11221.2 | AJ235445.2 | AY491646.1 | U42523.1 | AF479187.1 |
| Asterales | Alseuosmiaceae | *Platyspermation crassifolium* | AJ419700 | AJ419689 | | | |
| Asterales | Alseuosmiaceae | *Crispiloba disperma* | 1304285 | 15425563 | GQ983650 | | |
| Asterales | Alseuosmiaceae | *Wittsteinia vacciniacea* | X87399 | AJ318986 | | | |
| Asterales | Alseuosmiaceae | *Wittsteinia panderi* | | | GQ983647 | | |

| | | | | | | | |
|---|---|---|---|---|---|---|---|
| Asterales | Alseuosmiaceae | *Alseuosmia macrophylla* | X87377 | 6686770 | AJ429378 | 7595364 | |
| Asterales | Alseuosmiaceae | *Alseuosmia sp.* | | | | | GQ983579 |
| Asterales | Stylidaceae | *Donatia fascicularis* | AF307913.1 | | AJ429384.1 | | |
| Asterales | Stylidaceae | *Donatia sp.* | | AJ236203.1 | | AJ236012.1 | AF479189.1 |
| Asterales | Stylidaceae | *Forstera bidwillii* | AJ225055 | | GQ983645 | | |
| Asterales | Stylidaceae | *Forstera bellidifolia* | | AJ318976 | | | |
| Asterales | Stylidaceae | *Stylidium graminifolium* | L18790 | AJ236201 | | AJ236011 | GQ983592 |
| Asterales | Stylidaceae | *Stylidium majus* | | | AF542576 | | |
| Asterales | Menyanthaceae | *Nymphoides peltata* | EF173110 | | GQ983659 | | |
| Asterales | Menyanthaceae | *Nymphoides geminata* | | AJ236204 | | 6688858 | 19919627 |
| Asterales | Menyanthaceae | *Villarsia calthifolia* | L11685 | | | | |
| Asterales | Menyanthaceae | *Villarsia capitata* | | AJ318984 | | | |
| Asterales | Menyanthaceae | *Villarsia sp.* | | | GQ983655 | | |
| Asterales | Menyanthaceae | *Nephrophyllidium crista-galli* | EF173095 | | GQ983644 | | |
| Asterales | Menyanthaceae | *Fauria crista-galli* | | AJ318975 | | | |
| Asterales | Menyanthaceae | *Menyanthes trifoliata* | L14006.2 | AJ235533.2 | AJ429386.1 | AJ236009.1 | AF479185.1 |
| Asterales | Goodeniaceae | *Dampiera spicigera* | X87383 | AJ318971 | | | |
| Asterales | Goodeniaceae | *Dampiera diversifolia* | | | GQ983658 | | |
| Asterales | Goodeniaceae | *Goodenia ovata* | X87386 | AJ318977 | GQ983643 | | |
| Asterales | Goodeniaceae | *Scaevola aemula* | EU017199.1 | EU017162.1 | EU385394 | AJ236008.1 | |
| Asterales | Goodeniaceae | *Scaevola auriculata* | | | | | GQ983591.1 |
| Asterales | Calyceraceae | *Boopis anthemoides* | L13860.1 | | | | |
| Asterales | Calyceraceae | *Boopis graminea* | | AJ236199.1 | AJ429382.1 | AF107583.1 | AF479184.1 |
| Asterales | Calyceraceae | *Acicarpha tribuloides* | 1304279 | 15422203 | 22795872 | | |
| Asterales | Calyceraceae | *Moschopsis rosulata* | X87390 | AJ318979 | GQ983662 | | |
| Asterales | Asteraceae | *Barnadesia caryophylla* | 289491 | 14717953 | | | AF479246.1 |
| Asterales | Asteraceae | *Barnadesia arborea* | | | GQ983656 | | |
| Asterales | Asteraceae | *Barnadesia sp.* | | | | 4558886 | |
| Asterales | Asteraceae | *Gerbera jamesonii* | L13643 | AJ236200 | AF456789 | 4558887 | |
| Asterales | Asteraceae | *Echinops exaltatus* | 290607 | | 32364845 | | |
| Asterales | Asteraceae | *Echinops bannaticus* | | 15425573 | | | |
| Asterales | Asteraceae | *Tragopogon pratensis* | AY395563.1 | | | | |
| Asterales | Asteraceae | *Tragopogon dubius* | | AJ236197.1 | AJ633258.1 | U42502.1 | AF036493.1 |
| Asterales | Asteraceae | *Lactuca sativa* | 78675147 | 78675147 | 78675147 | 471875 | |
| Asterales | Asteraceae | *Cichorium intybus* | 289848 | 8452630 | 54021386 | | |
| Asterales | Asteraceae | *Tagetes erecta* | L13637 | | | | |
| Asterales | Asteraceae | *Tagetes sp.* | | AJ236206 | | 1777651 | |
| Asterales | Asteraceae | *Tagetes patula* | | | AF151515 | | |
| Asterales | Asteraceae | *Guizotia abyssinica* | NC_010601 | NC_010601 | AM411125 | | |
| Asterales | Asteraceae | *Helianthus annuus* | AF097517.1 | AJ236205.1 | DQ383815.1 | AF107577.1 | AF479183.1 |
| Bruniales | Columelliaceae | *Columellia oblonga* | Y10675. | AJ419676 | AJ429362 | GQ983564 | |
| Bruniales | Columelliaceae | *Desfontainia spinosa* | Z29670.1 | AJ419677.1 | AJ429363.1 | GQ983565.1 | GQ983582.1 |
| Bruniales | Bruniaceae | *Brunia albiflora* | AY490988 | | | | |
| Bruniales | Bruniaceae | *Brunia laevis* | | GQ983608 | | | |

| | | | | | | | |
|---|---|---|---|---|---|---|---|
| Bruniales | Bruniaceae | *Brunia alopecuroides* | | | AY490959 | | |
| Bruniales | Bruniaceae | *Berzelia lanuginosa* | L14391 | AF095731 | AY490955.1 | U42508 | |
| Apiales | Pennantiaceae | *Pennantia corymbosa* | AJ494842 | AJ494840 | AY188404 | GU476405 | AY189099 |
| Apiales | Pennantiaceae | *Pennantia cunninghamii* | | | | GQ983573 | |
| Apiales | Torricelliaceae | *Aralidium pinnatifidum* | AF299087 | | APU58627 | | AY189036 |
| Apiales | Torricelliaceae | *Toricellia tiliifolia* | AF299089 | | AY188410 | | AY189113 |
| Apiales | Torricelliaceae | *Melanophylla alnifolia* | U50254.1 | AF209625.1 | | AF206960.1 | AY189085.1 |
| Apiales | Torricelliaceae | *Melanophylla sp.* | | | AJ429373.1 | | |
| Apiales | Griseliniaceae | *Griselinia littoralis* | AF307916.1 | AJ236213.1 | AJ429372.1 | | |
| Apiales | Griseliniaceae | *Griselinia lucida* | | | | AF206922.1 | AF479197.1 |
| Apiales | Pittosporaceae | *Pittosporum japonicum* | L11202 | | | 532222 | |
| Apiales | Pittosporaceae | *Pittosporum verrucosum* | | AF528857 | | | |
| Apiales | Pittosporaceae | *Pittosporum tobira* | | | U58624 | | 37778912 |
| Apiales | Pittosporaceae | *Sollya heterophylla* | U50262.1 | AJ236214.1 | U58625.1 | AJ236001.1 | AY189109.1 |
| Apiales | Araliaceae | *Hydrocotyle vulgaris* | GQ983666 | GQ983599 | DQ133792 | | |
| Apiales | Araliaceae | *Hydrocotyle sibthorpioides* | | | | X16605 | |
| Apiales | Araliaceae | *Hydrocotyle verticillata* | | | | | AY189078 |
| Apiales | Araliaceae | *Aralia spinosa* | 289043 | GQ983605 | 22795866 | | 37778827 |
| Apiales | Araliaceae | *Panax quinquefolius* | U50250.1 | AJ236210.1 | | | AF479193.1, 37778890 |
| Apiales | Araliaceae | *Panax ginseng* | 51235292 | | AB044903.2 | | |
| Apiales | Araliaceae | *Panax zingiberensis* | | | | AB085764.1 | |
| Apiales | Araliaceae | *Cussonia spicata* | 1292997 | GQ983596 | 2281238 | | 37778849 |
| Apiales | Araliaceae | *Polyscias guilfoylei* | U50251 | GQ983620 | U58616 | | AY189102.1 |
| Apiales | Araliaceae | *Tetraplasandra hawaiensis* | U50257 | | U58593 | | 37778904 |
| Apiales | Araliaceae | *Tetraplasandra oahuensis* | | GQ983616 | | | |
| Apiales | Araliaceae | *Pseudopanax arboreus* | 1293017 | | 2281268 | | 37778895 |
| Apiales | Araliaceae | *Pseudopanax colensoi* | | GQ983637 | | | |
| Apiales | Araliaceae | *Hedera helix* | L01924.2 | AJ235489.2 | U58612.1, 18073960 | U42500.1 | AY189073.1 |
| Apiales | Araliaceae | *Schefflera arboricola* | U50255 | | 2281278 | | 37778900 |
| Apiales | Araliaceae | *Schefflera delavayi* | | GQ983613 | | | |
| Apiales | Araliaceae | *Tetrapanax papyriferus* | U50256 | GQ983624 | GQ434267 | | 37778903 |
| Apiales | Myodocarpaceae | *Myodocarpus involucratus* | AY188430 | | | | AY189095 |
| Apiales | Myodocarpaceae | *Myodocarpus fraxinifolius* | | GQ983630 | | GQ983570 | |
| Apiales | Myodocarpaceae | *Myodocarpus simplicifolius* | | | AF271754 | | |
| Apiales | Myodocarpaceae | *Delarbrea michieana* | U50243.1 | AJ236211.1 | U58608.1 | AF107573.1 | AF479196.1 |
| Apiales | Apiaceae | *Platysace lanceolata* | AY188434 | GQ983628 | GQ983657 | | AY189101.1 |
| Apiales | Apiaceae | *Mackinlaya confusa* | AY188426 | GQ983615 | AF271741 | GQ983568 | |
| Apiales | Apiaceae | *Mackinlaya macrosciadia* | | | | | AY189083 |
| Apiales | Apiaceae | *Azorella selago* | 34559276 | | 14276784 | | 37778838 |

| | | | | | | | |
|---|---|---|---|---|---|---|---|
| Apiales | Apiaceae | *Azorella caespitosa* | | GQ983635 | | GQ983561 | |
| Apiales | Apiaceae | *Arctopus echinatus* | AY188414 | | AF271761 | | AY189037.1 |
| Apiales | Apiaceae | *Arctopus dregei* | | GQ983621 | | | |
| Apiales | Apiaceae | *Sanicula gregari* | L11170 | | U58589 | | 37778899 |
| Apiales | Apiaceae | *Heteromorpha trifoliata* | HTU50227 | | HTU58565 | | 37778866 |
| Apiales | Apiaceae | *Heteromorpha arborescens* | | GQ983611 | | | |
| Apiales | Apiaceae | *Daucus carota* | 1374996 | 113200887 | 2281160 | | 37778850 |
| Apiales | Apiaceae | *Coriandrum sativum* | 336618 | GQ983603 | 2281158 | | AY189056.1 |
| Apiales | Apiaceae | *Angelica lucida* | 1292957 | | | | 37778823 |
| Apiales | Apiaceae | *Angelica sylvestris* | | GQ983632 | DQ133783 | | |
| Apiales | Apiaceae | *Angelica gigas* | | | | 200116396 | |
| Apiales | Apiaceae | *Anethum graveolens* | 156595776 | 156573702 | 156573664 | | |
| Apiales | Apiaceae | *Apium graveolens* | L01885.2 | AJ235396.2 | AJ429370.1 | AF206852.1 | AF479195.1 |
| Paracryphiales | Paracryphiaceae | *Quintinia verdonii* | AF299092 | | AJ429366 | | |
| Paracryphiales | Paracryphiaceae | *Quintinia quatrefagesii* | | AJ318983 | | GQ983576 | GQ983590 |
| Paracryphiales | Paracryphiaceae | *Sphenostemon lobosporus* | AJ403005 | GQ983631.1 | GQ983660.1 | | |
| Paracryphiales | Paracryphiaceae | *Paracryphia alticola* | AJ402983 | AJ419679 | AJ429367 | GQ983571 | GQ983589 |
| Dipsacales | Adoxaceae | *Viburnum acerifolia* | AF446927.1 | | | AJ236007.1 | |
| Dipsacales | Adoxaceae | *Viburnum opulus* | | AJ235640.2 | | | |
| Dipsacales | Adoxaceae | *Viburnum rhytidophyllum* | | | AJ429391.1 | | |
| Dipsacales | Adoxaceae | *Viburnum erosum* | | | | | JF321130.1 |
| Dipsacales | Adoxaceae | *Sambucus racemosa* | 17863820 | | 20530881 | | |
| Dipsacales | Adoxaceae | *Sambucus caerulea* | | GQ983634 | | | |
| Dipsacales | Adoxaceae | *Sambucus canadensis* | | | | 11066013 | |
| Dipsacales | Adoxaceae | *Sinadoxa corydalifolia* | AF446929 | GQ983638 | AF446899 | | |
| Dipsacales | Adoxaceae | *Tetradoxa omeiensis* | AF446931 | GQ983607 | AF446901 | | |
| Dipsacales | Adoxaceae | *Adoxa moschatellina* | AF446930.1 | GQ983610.1 | AF446900.1 | GQ983560.1 | GQ983578.1 |
| Dipsacales | Caprifoliaceae (Diervillaceae) | *Weigela hortensis* | AF446938 | GQ983609 | AF446908 | | |
| Dipsacales | Caprifoliaceae (Diervillaceae) | *Diervilla sessilifolia* | AF446937 | GQ983617 | AF446907 | GQ983566 | GQ983583 |
| Dipsacales | Caprifoliaceae | *Heptacodium miconioides* | AF446936 | GQ983604 | AF446906 | | |
| Dipsacales | Caprifoliaceae | *Lonicera japonica* | EU201192 | GQ983602 | | | GQ983587 |
| Dipsacales | Caprifoliaceae | *Lonicera orientalis* | | | 22796641 | | |
| Dipsacales | Caprifoliaceae | *Lonicera maackii* | | | | 1857127 | |
| Dipsacales | Caprifoliaceae | *Symphoricarpos albus* | L11682 | | | 1777647 | 19919683 |
| Dipsacales | Caprifoliaceae | *Symphoricarpos sp.* | | GQ983633 | | | |
| Dipsacales | Caprifoliaceae | *Symphoricarpos occidentalis* | | | GU168654 | | |
| Dipsacales | Caprifoliaceae | *Triosteum perfoliatum* | AF446935 | GQ983597 | GQ284972 | | |
| Dipsacales | Caprifoliaceae | *Leycesteria formosa* | AF446992 | GQ983636 | GQ284975 | | |
| Dipsacales | Caprifoliaceae (Linnaeaceae) | *Linnaea borealis* | AF446941 | GQ983619 | AF446911 | | |
| Dipsacales | Caprifoliaceae (Linnaeaceae) | *Abelia triflora* | AF206727.1 | AJ236209.1 | | AJ236004.1 | AF479200.1 |

| | | | | | | | |
|---|---|---|---|---|---|---|---|
| Dipsacales | Caprifoliaceae (Linnaeaceae) | *Abelia grandiflora* | | | AF446909.1 | | |
| Dipsacales | Caprifoliaceae (Linnaeaceae) | *Kolkwitzia amabilis* | AJ420877 | GQ983600 | AF446912 | | |
| Dipsacales | Caprifoliaceae (Linnaeaceae) | *Dipelta floribunda* | AJ420876 | | | | |
| Dipsacales | Caprifoliaceae (Linnaeaceae) | *Dipelta yunnanensis* | | GQ983629 | 20530893 | GQ983567 | GQ983584 |
| Dipsacales | Caprifoliaceae (Morinaceae) | *Morina longifolia* | AF446945 | GQ983601 | AF446915 | GQ983569 | |
| Dipsacales | Caprifoliaceae (Morinaceae) | *Zabelia tyaihyoni* | GQ983786 | GQ983627 | GQ983641 | | |
| Dipsacales | Caprifoliaceae (Dipsacaceae) | *Triplostegia glandulifera* | AF446949 | GQ983612 | AF446919 | GQ983577 | GQ983593 |
| Dipsacales | Caprifoliaceae (Dipsacaceae) | *Scabiosa columbaria* | AF446948.1 | GQ983595 | AF446918.1 | | |
| Dipsacales | Caprifoliaceae (Dipsacaceae) | *Scabiosa sp.* | | AJ236207.1 | | AJ236006.1 | AF479198.1 |
| Dipsacales | Caprifoliaceae (Dipsacaceae) | *Pterocephalus hookeri* | AF446946 | GQ983623 | AF446916 | | |
| Dipsacales | Caprifoliaceae (Dipsacaceae) | *Dipsacus sativus* | L13864.1 | AF209577.1 | 22795882 | | AF479231.1 |
| Dipsacales | Caprifoliaceae (Dipsacaceae) | *Dipsacus mitis* | | | AF446917.1 | | |
| Dipsacales | Caprifoliaceae (Dipsacaceae) | *Dipsacus sp.* | | | | U43150.1 | |
| Dipsacales | Caprifoliaceae (Valerianaceae) | *Patrinia triloba* | AF446951 | GQ983625 | AF446921 | GQ983572 | |
| Dipsacales | Caprifoliaceae (Valerianaceae) | *Nardostachys chinensis* | GQ436316 | | | | |
| Dipsacales | Caprifoliaceae (Valerianaceae) | *Valeriana jatamansi* | | GQ983614 | | | |
| Dipsacales | Caprifoliaceae (Valerianaceae) | *Nardostachys grandiflora* | | | GU188967 | | |
| Dipsacales | Caprifoliaceae (Valerianaceae) | *Valerianella locusta* | AF446954 | GQ983622 | DQ354186 | | |
| Dipsacales | Caprifoliaceae (Valerianaceae) | *Centranthus ruber* | 18873584 | GQ983626 | 20530909 | | |
| Dipsacales | Caprifoliaceae (Valerianaceae) | *Valeriana officinalis* | L13934.1 | AJ235637.2 | | AJ236003.1 | AF479199.1 |
| Dipsacales | Caprifoliaceae (Valerianaceae) | *Valeriana hirtella* | | | AJ429396.1 | | |
| Saxifragales | Peridiscaceae | *Soyauxia talbotii* | AM111356 | AM111357 | | AM111355 | DQ400572 |
| Saxifragales | Peridiscaceae | *Soyauxia sp.* | | | DQ241372 | | |
| Saxifragales | Peridiscaceae | *Peridiscus lucidus* | AY380356 | AY788274 | DQ411570 | AY674625 | DQ400571 |
| Saxifragales | Paeoniaceae | *Paeonia californica* | AF274593.1 | AF274683.1 | AF033589.1 | AF274603.1 | AF274658.1 |
| Saxifragales | Altingiaceae | *Liquidambar styraciflua* | AF119181.1 | AF092104.1 | AF133219.1 | U42553.1 | AF479217.1 |
| Saxifragales | Altingiaceae | *Altingia excelsa* | AF206732.1 | AF092103.1 | AF304520.1 | | AF479208.1 |
| Saxifragales | Altingiaceae | *Altingia sp.* | | | | U42552.1 | |
| Saxifragales | Daphniphyllaceae | *Daphniphyllum sp.* | L01900.2 | AF092118.1 | AF274612.1 | U42531.1 | AF479215.1 |
| Saxifragales | Cercidiphyllaceae | *Cercidiphyllum japonicum* | L11673.1 | AF092112.1 | AM396508.1, 209171822 | AF094534.1 | AF274639.1 |
| Saxifragales | Hamamelidaceae | *Rhodoleia henryi* | AF081072.1 | | | | |
| Saxifragales | Hamamelidaceae | *Rhodoleia championii* | | AF274674.1 | AF128833.1 | AF274599.1 | AF274664.1 |
| Saxifragales | Hamamelidaceae | *Exbucklandia populnea* | AF081071.1 | AF093379.1 | AF128831.1 | AF094550.1 | AF274647.1 |
| Saxifragales | Hamamelidaceae | *Disanthus cercidifolius* | AF081069.1 | AF093378.1 | AF128826.1 | AF094549.1 | AF274645.1 |
| Saxifragales | Hamamelidaceae | *Hamamelis mollis* | L01922.2 | | | | |
| Saxifragales | Hamamelidaceae | *Hamamelis helix* | | AF092105.1 | | | |
| Saxifragales | Hamamelidaceae | *Hamamelis virginiana* | | | AF013046.1 | AF094551.1 | AF036495.1 |

| | | | | | | | |
|---|---|---|---|---|---|---|---|
| Saxifragales | Hamamelidaceae | *Corylopsis pauciflora* | AF060710.1 | AF093377.1 | | AF094548.1 | |
| Saxifragales | Hamamelidaceae | *Corylopsis sinensis* | | | AF013038.1 | | AF274642.1 |
| Saxifragales | Pterostemonaceae | *Pterostemon rotundifolius* | L11203.2 | AJ235573.2 | AF274630.1 | U42547.1 | AF274663.1 |
| Saxifragales | Iteaceae | *Itea virginica* | 13539648 | | 149798891 | U42545.1 | 19919657 |
| Saxifragales | Iteaceae | *Itea ilicifolia* | | 6017805 | | | |
| Saxifragales | Iteaceae | *Choristylis rhamnoides* | 4586133 | 9799477 | 9864090 | 9502207 | 9799438 |
| Saxifragales | Grossulariaceae | *Ribes aureum* | L11204.2 | AF528859.1 | L34153.1 | L28143.1 | AF274665.1 |
| Saxifragales | Saxifragaceae | *Saxifraga cernua* | 459082 | | 845665 | | |
| Saxifragales | Saxifragaceae | *Saxifraga retusa* | | 8452772 | | | |
| Saxifragales | Saxifragaceae | *Saxifraga mertensiana* | | | | 1777719 | |
| Saxifragales | Saxifragaceae | *Saxifraga integrifolia* | | | | | 9799464 |
| Saxifragales | Saxifragaceae | *Sullivantia oregana* | U06219.1 | AF209682.1 | L34113.1 | U42812.1 | AF479212.1 |
| Saxifragales | Saxifragaceae | *Heuchera micrantha* | L01925.2 | | | L28139.1 | AF479211.1 |
| Saxifragales | Saxifragaceae | *Heuchera sanguinea* | | EU002163.1 | 157689191 | | |
| Saxifragales | Saxifragaceae | *Heuchera hirsutissima* | | | L34125.1 | | |
| Saxifragales | Crassulaceae | *Crassula marnierana* | L01899.1 | AJ235447.2 | AF115600.1 | U42525.1 | AF479210.1 |
| Saxifragales | Crassulaceae | *Kalanchoe daigremontiana* | 7240298 | 8439429 | | 1777669 | 9799449 |
| Saxifragales | Crassulaceae | *Kalanchoe zimbabwensis* | | | 13568519 | | |
| Saxifragales | Crassulaceae | *Sedum rubrotinctum* | L01956.2 | | | U42528.1 | |
| Saxifragales | Crassulaceae | *Sedum nudum* | | AJ235600.2 | | | AF274667.1 |
| Saxifragales | Crassulaceae | *Sedum aizoon* | | | AB038187.1 | | |
| Saxifragales | Crassulaceae | *Dudleya viscida* | L11182.2 | AJ235461.2 | AF274614.1 | U42526.1 | AF274646.1 |
| Saxifragales | Aphanopetalaceae | *Aphanopetalum resinosum* | AF274596.1 | AF274675.1 | AF274607.1, EF179066.1 | AF274600.1 | AF274637.1 |
| Saxifragales | Tetracarpaeaceae | *Tetracarpaea tasmanica* | L11207.2 | AF209688.1 | L34154.1 | U42549.1 | AF274669.1 |
| Saxifragales | Penthoraceae | *Penthorum sedoides* | L11197.2 | AJ235555.2 | AF274628.1, 148470644 | U42825.1 | AF274661.1 |
| Saxifragales | Haloragaceae | *Myriophyllum exalbescens* | 7240338 | 8452699 | | 1777686 | 9799453 |
| Saxifragales | Haloragaceae | *Myriophyllum drummondii* | | | 148470523 | | |
| Saxifragales | Haloragaceae | *Haloragis serra* | U26325.2 | | | | |
| Saxifragales | Haloragaceae | *Haloragis aspera* | | AJ235488.2 | AF274616.1 | | |
| Saxifragales | Haloragaceae | *Haloragis glauca* | | | 148470559 | | |
| Saxifragales | Haloragaceae | *Haloragis erecta* | | | | AF094547.1 | AF274648.1 |
| Vitales | Vitaceae | *Leea guineensis* | AJ235783.1 | AJ235520.2 | AF274621.1 | AF206951.1 | AF274653.1 |
| Vitales | Vitaceae | *Vitis aestivalis* | L01960.2 | | AF274635 | | AF479207.1 |
| Vitales | Vitaceae | *Vitis vinifera* | | DQ424856.1 | DQ424856.1 | | |
| Vitales | Vitaceae | *Vitis sp.* | | | | AF207053.1 | |
| Geraniales | Geraniaceae | *Pelargonium cotyledonis* | L14703 | AF035911 | EU922531 | AF206982 | |
| Geraniales | Geraniaceae | *Geranium tuberosum* | DQ452887 | | | | |
| Geraniales | Geraniaceae | *Geranium sanguineum* | | AF035906 | | | AF479129 |
| Geraniales | Geraniaceae | *Geranium palmatum* | | | EU922315 | | |
| Geraniales | Geraniaceae | *Geranium sp.* | | | | U42541 | |
| Geraniales | Vivianiaceae | *Viviania marifolia* | L14707.2 | AF209696.1 | | AF207054.1 | AF479130.1 |
| Geraniales | Melianthaceae | *Melianthus major* | 125991586 | 8439469 | | | |
| Geraniales | Melianthaceae | *Melianthus comosus* | | | 157689199 | | |

| | | | | | | | |
|---|---|---|---|---|---|---|---|
| Geraniales | Greyiaceae | *Greyia radlkoferi* | L11185.2 | AF209594.1 | AF542592.1 | U43151.1 | AF479227.1 |
| Myrtales | Combretaceae | *Terminalia catappa* | U26338 | AF209686 | GU135057 | AF207037 | |
| Myrtales | Combretaceae | *Terminalia boivinii* | | | | | AF479147 |
| Myrtales | Lythraceae | *Lythrum hyssopyfolia* | L10218.1 | | HM850986.1 | | |
| Myrtales | Lythraceae | *Lythrum salicaria* | | AF209621.1 | | AF206955.1 | AF479240.1 |
| Myrtales | Lythraceae | *Lythrum flagellare* | | | 157689197 | | |
| Myrtales | Onagraceae | *Fuchsia procumbens* | AM235668 | AJ235477 | AJ581440 | AM235543 | AM235615 |
| Myrtales | Onagraceae | *Oenothera speciosa* | 255755669 | | | 472042 | |
| Myrtales | Onagraceae | *Oenothera parviflora* | | NC_010362.1 | 10362.1 | | |
| Myrtales | Onagraceae | *Oenothera macrocarpa* | | | | | 133918150 |
| Myrtales | Onagraceae | *Clarkia xantiana* | L01896 | AF209564 | | U67930 | AF479148 |
| Myrtales | Penaeaceae | *Olinia cymosa* | 7240565 | | | | |
| Myrtales | Penaeaceae | *Olinia emarginata* | | GQ497640.1 | | | |
| Myrtales | Penaeaceae | *Olinia ventosa* | | | 28569321 | 133918022 | 133918094 |
| Myrtales | Crypteroniaceae | *Crypteronia paniculata* | AF215545, 24899637 | | AY151566 | AM235482 | EU002152 |
| Myrtales | Melastomataceae | *Mouriri cyphocarpa* | U26327 | AF209634 | | AF206965 | |
| Myrtales | Melastomataceae | *Mouriri myrtilloides* | | | JQ588330.1 | | |
| Myrtales | Melastomataceae | *Clidemia rubra* | AF215535 | | | | |
| Myrtales | Melastomataceae | *Clidemia petiolaris* | | AJ235439 | | AM235516 | AM235588 |
| Myrtales | Melastomataceae | *Clidemia septuplinervia* | | | GQ981968 | | |
| Myrtales | Melastomataceae | *Clidemia dentata* | | | 157689177 | | |
| Myrtales | Vochysiaceae | *Qualea sp.* | 433100 | 14718196 | | 7595522 | |
| Myrtales | Vochysiaceae | *Qualea grandiflora* | | | 17226036 | | |
| Myrtales | Vochysiaceae | *Vochysia hondurensis* | U26340.2 | | AY572446.1 | | |
| Myrtales | Vochysiaceae | *Vochysia rufescens* | | AJ235644.2 | | | |
| Myrtales | Vochysiaceae | *Vochysia tucanorum* | | | | AM235540.1 | AM235612.1 |
| Myrtales | Myrtaceae | *Heteropyxis natalensis* | U26326 | AF209597 | AF368208 | AF206927 | AM235609 |
| Myrtales | Myrtaceae | *Metrosideros nervulosa* | AJ235785 | AJ235535 | | | |
| Myrtales | Myrtaceae | *Metrosideros diffusa* | | | AY521542 | | |
| Myrtales | Myrtaceae | *Metrosideros excelsa* | | | | AM235534 | AM235606 |
| Myrtales | Myrtaceae | *Eucalyptus melliodora* | 240253140 | | | | |
| Myrtales | Myrtaceae | *Eucalyptus grandis* | | NC_014570.1 | | | |
| Myrtales | Myrtaceae | *Eucalyptus curtisii* | | | 41324109 | | |
| Myrtales | Myrtaceae | *Eucalyptus lehmannii* | | | | 133918060 | 133918132 |
| Myrtales | Myrtaceae | *Myrtus communis* | 15530016 | JF268426 | 164606883 | | 157689142 |
| Crossosomatales | Staphyleaceae | *Staphylea trifolia* | AY646111.1 | AJ235611.2 | | AJ235978.1 | AF479133.1 |
| Crossosomatales | Staphyleaceae | *Staphylea colchica* | | | EU002189.1, 157689209 | | |
| Crossosomatales | Guamatelaceae | *Guamatela tuerckheimii* | DQ443463.1 | DQ443453.1 | DQ443460.1 | | |
| Crossosomatales | Stachyuraceae | *Stachyurus praecox* | 7414637 | 8452777 | | AF207025.1 | |
| Crossosomatales | Stachyuraceae | *Stachyurus chinensis* | | | 121491005 | | |
| Crossosomatales | Crossosomataceae | *Crossoma bigelovii* | AY101844.1 | | DQ443456.1 | AF193942.1 | |

| | | | | | | | |
|---|---|---|---|---|---|---|---|
| Crossosomatales | Crossosomataceae | *Crossoma californicum* | | AF209571.1 | | | AF479131.1 |
| Crossosomatales | Aphloiaceae | *Aphloia theiformis* | AF206735 | AF209528.1 | HQ680692.1 | AF206851 | AF479132 |
| Crossosomatales | Geissolomataceae | *Geissoloma marginatum* | X83990.1 | HQ680710.1 | HQ680697.1 | | AF222378.1 |
| Crossosomatales | Ixerbaceae | *Ixerba brexioides* | 4530131 | 14718088 | 157689193 | 4530133 | |
| Crossosomatales | Strasburgeriaceae | *Strasburgeria robusta* | AJ403007 | AF502597 | HQ680701.1 | AF502596 | |
| Picramniales | Picramniaceae | *Picramnia polyantha* | AF127025 | | | | |
| Picramniales | Picramniaceae | *Picramnia pentandra* | | AJ235559 | | | |
| Sapindales | Biebersteiniaceae | *Biebersteinia orphanidis* | AF035920 | AF035921 | | GQ497568.1 | |
| Sapindales | Nitrariaceae | *Nitraria sphaerocarpa* | 88174765 | | | | |
| Sapindales | Nitrariaceae | *Nitraria retusa* | | GQ497651.1 | 157689201 | | 157689143 |
| Sapindales | Nitrariaceae | *Nitraria praevisa* | | | | GQ497579.1 | |
| Sapindales | Burseraceae | *Bursera inaguensis* | L01890 | AF035899 | | AF206877 | AY177421 |
| Sapindales | Burseraceae | *Bursera fagaroides* | | | AY594462.1 | | |
| Sapindales | Anacardiaceae | *Schinus molle* | 7261037 | 4063563 | 195540515 | 7595535 | |
| Sapindales | Anacardiaceae | *Rhus copallina* | U00440 | AF035912 | | | |
| Sapindales | Anacardiaceae | *Rhus transvaalensis* | | | EU214283 | | |
| Sapindales | Anacardiaceae | *Rhus typhina* | | | | GU476470 | |
| Sapindales | Sapindaceae | *Cupaniopsis anacardioides* | L13182.2 | AF035903.1 | AY724283.1 | AF206896.1 | AF479139.1 |
| Sapindales | Sapindaceae | *Aesculus pavia* | U39277.2 | AF035894.1 | | AF206838.1 | AF479138.1 |
| Sapindales | Sapindaceae | *Aesculus glabra* | | | AY968671.1 | | |
| Sapindales | Meliaceae | *Trichilia emetica* | U39082.2 | AJ235629.2 | AY128202.1 | AF207045.1 | AY128171.1 |
| Sapindales | Meliaceae | *Swietenia macrophylla* | U39080.2 | AJ235616.2 | AY128200.1 | AF207031.1 | AF479241.1 |
| Sapindales | Rutaceae | *Citrus paradisi* | AJ238407.1 | AJ238408.1 | | | |
| Sapindales | Rutaceae | *Citrus trifoliata* | | | HM163960.1 | | |
| Sapindales | Rutaceae | *Citrus aurantium* | | | | U38312.1 | AY177420.1 |
| Sapindales | Simaroubaceae | *Ailanthus altissima* | U02726 | AF035895 | EF489111 | AF206842 | |
| Huerteales | Gerrardiniaceae | *Gerrardina foliosa* | AY757086.1 | AY757085.1 | FM179924.1 | | |
| Huerteales | Tapisciaceae | *Tapiscia sinensis* | AF206825 | AF209685 | EU002190, 195540518 | AF207034 | AF479146 |
| Huerteales | Dipentodontaceae | *Dipentodon sinicus* | 27448213 | | 22795884 | 27462221 | |
| Huerteales | Dipentodontaceae | *Perrottetia ovata* | AY935737 | AY935842 | AY935916 | AY929358 | AY935806 |
| Malvales | Neuradaceae | *Neurada procumbens* | U06814 | AF209637 | | AF206970 | AF479225 |
| Malvales | Bixaceae | *Bixa orellana* | 2502007 | 4063529 | 195540524 | 7595388 | 19919670 |
| Malvales | Cistaceae | *Helianthemum kahiricum* | 226358319 | | | | |
| Malvales | Cistaceae | *Helianthemum grandiflorum* | | 4063549 | | 7595446 | |
| Malvales | Cistaceae | *Helianthemum scopulicola* | | | 71668271 | | |
| Malvales | Dipterocarpaceae | *Anisoptera marginata* | Y15144 | AF035918 | AJ581409 | AF206849 | |
| Malvales | Thymelaeaceae | *Thymelaea hirsuta* | Y15151 | AJ235626 | EU002191 | AF207041 | AF479234 |
| Malvales | Malvaceae | *Ochroma pyramidale* | AF206800.1 | AF035910.1 | AY321172.1 | AF206975.1 | AF479135.1 |
| Malvales | Malvaceae | *Bombax buonopozense* | AF022118.1 | | AY321171.1 | | |
| Malvales | Malvaceae | *Bombax ceiba* | | AJ233051.1 | | U42507.1 | AF479134.1 |
| Malvales | Malvaceae | *Durio zibethinus* | AF206764.1 | AF209580.1 | AB289826.1 | AF206905.1 | AF479136.1 |
| Malvales | Malvaceae | *Gossypium hirsutum* | 11562 | 4995178 | | 1777727 | |

| | | | | | | | |
|---|---|---|---|---|---|---|---|
| Malvales | Malvaceae | *Gossypium longicalyx* | | | 15187153 | | |
| Malvales | Malvaceae | *Sterculia lanceolata* | AY082362.1 | | | | |
| Malvales | Malvaceae | *Sterculia apetala* | | AJ233089.1 | | | AF479137.1 |
| Malvales | Malvaceae | *Sterculia tragacantha* | | | AY321178.1 | | |
| Malvales | Malvaceae | *Sterculia recordiana* | | | | AF207029.1 | |
| Brassicales | Tropaeolaceae | *Tropaeolum tricolor* | AF254059.1 | AF035917.1 | | | AF479144.1 |
| Brassicales | Tropaeolaceae | *Tropaeolum majus* | | | AY483224.1 | L28750.1 | |
| Brassicales | Caricaceae | *Carica papaya* | M95671.1 | AF035901.1 | AY042564.1, 45775521 | U42514.1 | AF479145.1 |
| Brassicales | Limnanthaceae | *Floerkea proserpinacoides* | L12679.2 | AF035904.1 | EU002178.1 | U42784.1 | AF479143.1 |
| Brassicales | Koeberliniaceae | *Koeberlinia spinosa* | L14600 | AF209612 | AY483222 | U42512 | |
| Brassicales | Bataceae | *Batis maritima* | M88341 | AF209538 | AY483219 | U42504 | |
| Brassicales | Resedaceae | *Reseda alba* | 7240435 | 14718202 | | 3265097 | |
| Brassicales | Resedaceae | *Reseda lutea* | | | 195540527 | | |
| Brassicales | Gyrostemonaceae | *Gyrostemon thesioides* | FJ212210 | | | | |
| Brassicales | Gyrostemonaceae | *Gyrostemon racemigerus* | | GQ497646.1 | | | |
| Brassicales | Gyrostemonaceae | *Gyrostemon tepperi* | | | AY483237 | AF070971 | |
| Brassicales | Tovariaceae | *Tovaria pendula* | FJ212209.1 | | AY483242.1 | | |
| Brassicales | Pentadiplandraceae | *Pentadiplandra brazzeana* | U38533.1 | | AY483239.1 | AF070972.1 | |
| Brassicales | Capparaceae | *Capparis spinosa* | AY167985 | AF035900 | EU371772 | EU090942 | |
| Brassicales | Capparaceae | *Capparis sandwichiana* | | | | | AF479140 |
| Brassicales | Brassicaceae | *Arabidopsis thaliana* | U91966.1 | AJ971660, NC_000932 | AF144348 | X16077 | X52320.1 |
| Brassicales | Brassicaceae | *Raphanus sativus* | GQ184382 | AJ277564 | GQ248193 | | AY366932 |
| Brassicales | Brassicaceae | *Brassica napus* | AF267640 | AF267641 | AB354273 | | |
| Brassicales | Brassicaceae | *Brassica oleracea* | | | | AF513990 | |
| Brassicales | Brassicaceae | *Brassica rapa* | | | | | EF470522 |
| Zygophyllales | Krameriaceae | *Krameria ixine* | EU644679 | AJ235514 | EU604050 | AF206948 | AF479116 |
| Zygophyllales | Zygophyllaceae | *Guaiacum guatemalense* | 2467134 | | | | |
| Zygophyllales | Zygophyllaceae | *Guaiacum sanctum* | | 8452659 | | AY674599 | |
| Zygophyllales | Zygophyllaceae | *Guaiacum officinale* | | | 89242613 | | |
| Zygophyllales | Zygophyllaceae | *Larrea tridentata* | AF200474 | AY935860 | AY935935 | AY929372 | AY935818 |
| Fabales | Quillajaceae | *Quillaja saponaria* | U06822 | GQ497659.1 | AY386843 | | |
| Fabales | Polygalaceae | *Polygala cruciata* | L01945.2 | AJ235568.1 | | | AF479233.1 |
| Fabales | Polygalaceae | *Polygala californica* | | | AY386842.1 | | |
| Fabales | Polygalaceae | *Polygala pauciflora* | | | | U42797.1 | |
| Fabales | Surianaceae | *Stylobasium spathulatum* | U06828 | AF209681 | EU604032 | AF207030 | |
| Fabales | Fabaceae | *Schotia afra* | AM235016 | | | | |
| Fabales | Fabaceae | *Schotia brachypetala* | | | EU362038 | X66779 | X66764 |
| Fabales | Fabaceae | *Cercis canadensis* | U74188 | DQ401328 | EU361912 | | |
| Fabales | Fabaceae | *Bauhinia galpinii* | AM234262 | | EU361875 | X66777 | X66755 |
| Fabales | Fabaceae | *Bauhinia sp.* | | AF209540 | | | |
| Fabales | Fabaceae | *Ceratonia siliqua* | U74203 | | EU361911 | X66778 | X66758 |
| Fabales | Fabaceae | *Albizia julibrissin* | Z70147 | AF209524 | AY386855 | GU476373 | X66754 |
| Fabales | Fabaceae | *Acacia cavenia* | Z70145 | | | | |
| Fabales | Fabaceae | *Acacia pulchella* | | EU811863 | | | |

| | | | | | | | |
|---|---|---|---|---|---|---|---|
| Fabales | Fabaceae | *Acacia anegadensis* | | | HM020706 | | |
| Fabales | Fabaceae | *Acacia fimbriata* | | | | X66780 | X66753 |
| Fabales | Fabaceae | *Mimosa speggazzinii* | Z70151 | | | X66686 | X66762 |
| Fabales | Fabaceae | *Mimosa tenuiflora* | | EU811854 | | | |
| Fabales | Fabaceae | *Mimosa pudica* | | | AY177668 | | |
| Fabales | Fabaceae | *Indigofera australis* | AF308711 | | | | |
| Fabales | Fabaceae | *Indigofera sokotrana* | | | GU951671 | | |
| Fabales | Fabaceae | *Indigofera gerardiana* | | | | X66689 | X66760 |
| Fabales | Fabaceae | *Glycine max* | EU717256 | AY935856 | AF142700 | X02623 | X66759 |
| Fabales | Fabaceae | *Erythrina crista-galli* | Z70170 | | AY386869 | AF525296 | |
| Fabales | Fabaceae | *Erythrina sousae* | | EU717512 | | | |
| Fabales | Fabaceae | *Lotus corniculatus* | U74213 | | HM049505 | 50880779 | |
| Fabales | Fabaceae | *Lotus japonicus* | | AP002983 | | | |
| Fabales | Fabaceae | *Pisum sativum* | 255957401 | 12147 | 52789062 | U43011 | |
| Fabales | Fabaceae | *Cicer arietinum* | 18032750 | NC_011163.1 | | 32968200 | |
| Fabales | Fabaceae | *Cicer cuneatum* | | | 125860457 | | |
| Fabales | Fabaceae | *Astragalus membranaceus* | EF685978 | | EF685992 | AF359594 | |
| Rosales | Rosaceae | *Spiraea vanhouttei* | L11206.2 | | | U42801.1 | |
| Rosales | Rosaceae | *Spiraea betulifolia* | | AJ235608.2 | | | AF479103.1 |
| Rosales | Rosaceae | *Spiraea cantoniensis* | | | AF288127.1 | | |
| Rosales | Rosaceae | *Prunus persica* | AF206813.1 | AF209660.1 | AF288117.1 | L28749.1 | AY935820.1 |
| Rosales | Rosaceae | *Photinia fraseri* | L11200.2 | AF209653.1 | | U42800.1 | AF479101.1 |
| Rosales | Rosaceae | *Photinia serrulata* | | | AF288111.1 | | |
| Rosales | Barbeyaceae | *Barbeya oleoides* | U60314.1 | AF209535.1 | JF317418.1 | JF317358.1 | JF317379.1 |
| Rosales | Elaeagnaceae | *Elaeagnus angustifolia* | U17038.1 | | | | |
| Rosales | Elaeagnaceae | *Elaeagnus sp.* | | AJ235462.2 | | | |
| Rosales | Elaeagnaceae | *Elaeagnus umbellata* | | | AY257529.1 | L24090.1 | |
| Rosales | Elaeagnaceae | *Elaeagnus bockii* | | | | | JF317385.1 |
| Rosales | Dirachmaceae | *Dirachma socotrana* | AJ225789.1 | | JF317423.1 | JF317364.1 | JF317383.1 |
| Rosales | Rhamnaceae | *Rhamnus lycioides* | 9968785 | | | | |
| Rosales | Rhamnaceae | *Rhamnus catharticus* | | 8452734 | | | |
| Rosales | Rhamnaceae | *Rhamnus cathartica* | | | 30421077 | 6689072 | |
| Rosales | Rhamnaceae | *Rhamnus utilis* | | | | | JF317393.1 |
| Rosales | Rhamnaceae | *Ceanothus sanguineus* | U06795.1 | AF209558.1 | | U42799.1 | AF479102.1 |
| Rosales | Rhamnaceae | *Ceanothus pumilus* | | | AF049841.1 | | |
| Rosales | Ulmaceae | *Zelkova serrata* | D86317.1, AF206835 | AF209699.1 | | U42819.1 | AF479099.1 |
| Rosales | Ulmaceae | *Zelkova schneideriana* | | | AF345328.1 | | |
| Rosales | Cannabaceae | *Celtis yunnanensis* | L12638.2 | | | U42818.1 | AF479098.1 |
| Rosales | Cannabaceae | *Celtis philippensis* | | AY263961.1 | AY263925.1 | | |
| Rosales | Cannabaceae | *Cannabis sativa* | 24634979 | JF317400.1 | 16224059 | JF317360.1 | 157689139 |
| Rosales | Cannabaceae | *Humulus lupulus* | AF206777.1 | AF209599.1 | AY257528.1 | AF206931.1 | AF223066.1 |
| Rosales | Moraceae | *Morus indica* | DQ226511.1 | DQ226511.1 | | | |
| Rosales | Moraceae | *Morus alba* | | | AY257531.1 | L24398.1 | |
| Rosales | Moraceae | *Morus nigra* | | | | | AF479232.1 |
| Rosales | Moraceae | *Ficus benjamina* | AF500350.1 | | | | |
| Rosales | Moraceae | *Ficus sp.* | | AF209587.1 | | AF206911.1 | |
| Rosales | Moraceae | *Ficus carica* | | | AY257530.1 | | |
| Rosales | Moraceae | *Ficus tikoua* | | | | | JF317386.1 |

| | | | | | | | |
|---|---|---|---|---|---|---|---|
| Rosales | Urticaceae | *Pilea depressa* | AF500359 | | | | |
| Rosales | Urticaceae | *Pilea cadierei* | | AF209654 | JF317431.1 | U42820 | |
| Rosales | Urticaceae | *Pilea fontana* | | | | | AY686776 |
| Rosales | Urticaceae | *Urtica dioica* | AF500361.1 | 8452793 | | | |
| Rosales | Urticaceae | *Urtica cannabina* | | | 195540529 | | |
| Rosales | Urticaceae | *Boehmeria nivea* | AJ235801 | | | AF206870 | AY686767 |
| Rosales | Urticaceae | *Boehmeria platanifolia* | | | AF353579 | | |
| Fagales | Nothofagaceae | *Nothofagus antarctica* | AY263939 | AY147106 | AY263924 | AY147111 | |
| Fagales | Nothofagaceae | *Nothofagus solandri* | | | 4586826 | | |
| Fagales | Fagaceae | *Fagus grandifolia* | AY263936.1 | AY147105.1 | | AF206910.1 | AY935813.1 |
| Fagales | Fagaceae | *Fagus sylvatica* | | | AB046507.1 | | |
| Fagales | Fagaceae | *Quercus rubra* | AB125026.1 | AF132888.1 | AY312058.1 | AF132892.1 | |
| Fagales | Fagaceae | *Quercus suber* | | | | | AY428812.1 |
| Fagales | Fagaceae | *Chrysolepis sempervirens* | AF061995.1 | AF209563.1 | | AF206886.1 | AF479107.1 |
| Fagales | Fagaceae | *Chrysolepis chrysophylla* | | | FJ185045.1 | | |
| Fagales | Casuarinaceae | *Casuarina cunninghamiana* | 436798 | | | | |
| Fagales | Casuarinaceae | *Casuarina litorea* | | 8439266 | | | |
| Fagales | Casuarinaceae | *Casuarina cristata* | | | 31788915 | | |
| Fagales | Casuarinaceae | *Casuarina equisetifolia* | | | | U42515 | |
| Fagales | Betulaceae | *Alnus incana* | 297535 | | | | |
| Fagales | Betulaceae | *Alnus sinuata* | | 37729409 | | 32815611 | |
| Fagales | Betulaceae | *Alnus japonica* | | | 18146885 | | |
| Fagales | Betulaceae | *Alnus glutinosa* | | | | | 19919547 |
| Fagales | Juglandaceae | *Juglans mandshurica* | AY263932.1 | AY263952.1 | | | |
| Fagales | Juglandaceae | *Juglans nigra* | | | AF118036 | AF206943.1 | AF479105.1 |
| Fagales | Myricaceae | *Myrica gale* | 436805 | | 31788949 | | |
| Fagales | Myricaceae | *Myrica esculenta* | | | | 218217868 | |
| Fagales | Myricaceae | *Morella cerifera* | AJ626759.1 | AJ235537.1 | AY491657.1 | AF206967.1 | AF479247.1 |
| Cucurbitales | Anisophylleaceae | *Anisophyllea fallax* | AY935742 | AY935849 | AY935923 | AY929365 | AY935807 |
| Cucurbitales | Anisophylleaceae | *Anisophyllea sororia* | | | 65332693 | | |
| Cucurbitales | Corynocarpaceae | *Corynocarpus laevigatus* | AF148994.1 | AJ235446.2 | AY491652.1 | AF206892.1 | AF479110.1 |
| Cucurbitales | Coriariaceae | *Coriaria ruscifolia* | AF148999.1 | AY968430.1 | AB016462.1 | AY968395.1 | AY968408.1 |
| Cucurbitales | Coriariaceae | *Coriaria myrtifolia* | | AF092117.1 | AF542600.2 | | |
| Cucurbitales | Tetramelaceae | *Tetrameles nudiflora* | L21943.1 | AF209689.1 | AY968458.1 | U41502.1 | AY968422.1 |
| Cucurbitales | Datiscaceae | *Datisca cannabina* | L21939.1 | AJ235450.2 | AB016467.1 | AF008952.1 | |
| Cucurbitales | Datiscaceae | *Datisca glomerata* | | | | | AY968411.1 |
| Cucurbitales | Begoniaceae | *Begonia sanguinea* | L01888.2 | | | | |
| Cucurbitales | Begoniaceae | *Begonia metallicaxsanguinea* | | AF209541.1 | | | AF479109.1 |
| Cucurbitales | Begoniaceae | *Begonia grandis* | | | AB016466.1 | | |
| Cucurbitales | Begoniaceae | *Begonia oxyloba* | | | 67772428 | | |
| Cucurbitales | Begoniaceae | *Begonia luxurians* | | | | AF534762.1 | |
| Cucurbitales | Cucurbitaceae | *Xerosicyos danguyi* | AY973026 | AJ235648 | AY968459 | AY973017 | AY968423 |
| Cucurbitales | Cucurbitaceae | *Dendrosicyos socotranus* | AY973022 | AY968433 | AY973018 | AY968397 | AY968412 |
| Cucurbitales | Cucurbitaceae | *Cucurbita pepo* | 17135916 | 14718023 | 111053130 | GQ856148 | AF479108.1 |
| Cucurbitales | Cucurbitaceae | *Cucumis melo* | DQ535800 | | DQ536659 | | |
| Cucurbitales | Cucurbitaceae | *Cucumis sativus* | | AF209572 | | | |
| Cucurbitales | Cucurbitaceae | *Cucumis anguria* | | | | 33330857 | |
| Cucurbitales | Cucurbitaceae | *Coccinia sessilifolia* | AY968520 | AY968427 | AY968446 | AY973011 | AY968404 |
| Celastrales | Lepidobotryaceae | *Lepidobotrys staudtii* | AJ402966 | AY935831 | AY935904 | AY929346 | |
| Celastrales | Lepidobotryaceae | *Ruptiliocarpon caracolito* | AJ402997 | AY788275 | 62902955 | AY929361 | |

| | | | | | | | |
|---|---|---|---|---|---|---|---|
| Celastrales | Celastraceae (Parnassiaceae) | *Parnassia fimbriata* | L01939.2 | | | | AF036496.1 |
| Celastrales | Celastraceae (Parnassiaceae) | *Parnassia glauca* | AY935729 | | AY935908 | | |
| Celastrales | Celastraceae (Parnassiaceae) | *Parnassia palustris* | | AJ235552.2 | AY935910.1 | AY929353.1 | |
| Celastrales | Celastraceae | *Siphonodon celastrineus* | AF206821.1 | AF209676.1 | AY935919.1 | AF207021.1 | |
| Celastrales | Celastraceae | *Siphonodon australe* | X83996 | | | | |
| Celastrales | Celastraceae | *Siphonodon australis* | | | | | AF222346.1 |
| Celastrales | Celastraceae | *Denhamia celastroides* | AJ402941 | AY788267 | | | |
| Celastrales | Celastraceae | *Denhamia pittosporoides* | | | EU329001 | EU328772 | |
| Celastrales | Celastraceae | *Denhamia viridissima* | | | | | 164370794 |
| Celastrales | Celastraceae | *Stackhousia minima* | AJ235795.1 | AJ235610.2 | EF135596 | AF207026.1 | AF479114.1 |
| Celastrales | Celastraceae | *Stackhousia monogyna* | | | AY935920.1 | | |
| Celastrales | Celastraceae | *Paxistima canbyi* | AY788198 | AY788273 | | AY674623 | 12082562 |
| Celastrales | Celastraceae | *Paxistima myrsinites* | | | 164652424 | | |
| Celastrales | Celastraceae | *Euonymus alatus* | AY788197 | | | X16600 | |
| Celastrales | Celastraceae | *Euonymus americanus* | | EU002160 | EU002170 | | 157689137 |
| Celastrales | Celastraceae | *Tripterygium regelii* | AY788193 | AY788260 | 164652380 | AY788161 | 12082560 |
| Celastrales | Celastraceae | *Celastrus scandens* | AY788195 | AY788264 | | | 12082567 |
| Celastrales | Celastraceae | *Celastrus orbiculatus* | | | EF135517 | AY788162 | |
| Celastrales | Celastraceae | *Maytenus senegalensis* | AY380353 | AY788272 | | | |
| Celastrales | Celastraceae | *Maytenus floribunda* | | | EU328961 | | 164370813 |
| Celastrales | Celastraceae | *Maytenus arbutifolia* | | | | AY674616 | |
| Celastrales | Celastraceae | *Plagiopteron suaveolens* | 257783293 | AJ235562 | 71891428 | AF206993 | 12082633 |
| Celastrales | Celastraceae | *Elaeodendron orientale* | AY380347 | AY788269 | EF135531 | AY674593 | |
| Celastrales | Celastraceae | *Elaeodendron vitiense* | | | | | 81250859 |
| Celastrales | Celastraceae | *Brexia madagascariensis* | L11176.1 | AJ235419.2 | AY935899.1 | U42543.1 | AF479112.1 |
| Oxalidales | Huaceae | *Hua gabonii* | FJ670185 | FJ669995 | FJ670056, 62902925 | AY929345 | AY935796 |
| Oxalidales | Huaceae | *Afrostyrax lepidophyllus* | AY935721 | AY935824 | AY935896 | AY929339 | AY935792 |
| Oxalidales | Connaraceae | *Rourea minor* | FJ707537.1 | FJ669994.1 | EF135591.1 | EF135603.1 | |
| Oxalidales | Connaraceae | *Connarus conchocarpus* | U06798.1 | | 157689179 | | |
| Oxalidales | Connaraceae | *Connarus championii* | | AY935852.1 | AY935928.1 | AY929368.1 | AY935810.1 |
| Oxalidales | Oxalidaceae | *Oxalis dillenii* | L01938.2 | AF209642.1 | | AF206978.1 | AF479230.1 |
| Oxalidales | Oxalidaceae | *Oxalis acetosella* | | FJ707531 | | | |
| Oxalidales | Oxalidaceae | *Oxalis latifolia* | | | EU002186.1 | | |
| Oxalidales | Oxalidaceae | *Dapania racemosa* | AY788196 | AY788266 | FJ670049.1 | AY674590 | |
| Oxalidales | Oxalidaceae | *Averrhoa carambola* | L14692.2 | AJ235404.2 | AY935924.1, 62902967 | AF206859.1 | AF479127.1 |
| Oxalidales | Brunelliaceae | *Brunellia acutangula* | FJ707536 | FJ669993 | EF135512 | FJ669718 | |
| Oxalidales | Cephalotaceae | *Cephalotus follicularis* | L01894 | AY788265 | FJ670045.1 | 1777658 | |

| | | | | | | | |
|---|---|---|---|---|---|---|---|
| Oxalidales | Cunoniaceae | *Eucryphia lucida* | L01918 | | 157689183 | U42533. | 2687434 |
| Oxalidales | Cunoniaceae | *Eucryphia milliganii* | | AJ235470 | | | |
| Oxalidales | Cunoniaceae | *Davidsonia pruriens* | AF291934.1 | AF209574.1 | U92846.1 | AF206897.1 | AY935812.1 |
| Oxalidales | Elaeocarpaceae | *Sloanea latifolia* | AF022131.1 | | | U42826.1 | |
| Oxalidales | Elaeocarpaceae | *Sloanea berteriana* | | AJ235603.2 | | | AF479126.1 |
| Oxalidales | Elaeocarpaceae | *Sloanea australis* | | | AY935938.1 | | |
| Oxalidales | Elaeocarpaceae | *Elaeocarpus sphaericus* | 17135933 | | | 7595426 | 27803696 |
| Oxalidales | Elaeocarpaceae | *Elaeocarpus reticulatus* | | 62902858 | 62902981 | | |
| Oxalidales | Elaeocarpaceae | *Crinodendron patagua* | AF291940.1 | | | | AY935811.1 |
| Oxalidales | Elaeocarpaceae | *Crinodendron hookerianum* | | AF209570.1 | AY491655.1 | AF206893.1 | |
| Malpighiales | Humiriaceae | *Sacoglottis sp.* | AB233890 | AB233682 | 118917529 | 119368155 | |
| Malpighiales | Humiriaceae | *Vantanea guianensis* | Z75679 | AY788261 | EF135600 | AY674639 | |
| Malpighiales | Humiriaceae | *Humiria balsaminifera* | L01926.2 | AJ235495.2 | EF135549.1 | AF206930.1 | AY935815.1 |
| Malpighiales | Ixonanthaceae | *Ochthocosmus longipedicellatus* | 257853507 | 257853485 | EF135573 | AY674621 | |
| Malpighiales | Irvingiaceae | *Irvingia malayana* | AF206782 | AF209605 | EF135553 | AF206939 | |
| Malpighiales | Irvingiaceae | *Klainedoxa gabonensis* | AY663630 | AY788232 | EF135556 | AY674610 | |
| Malpighiales | Clusiaceae | *Symphonia tanalensis* | AB233852.1 | AB233644.1 | AB233748.1 | AB233540.1 | |
| Malpighiales | Clusiaceae | *Garcinia xanthochymus* | 22003621 | | | | |
| Malpighiales | Clusiaceae | *Garcinia subelliptica* | | 119368193 | | 119368140 | |
| Malpighiales | Clusiaceae | *Garcinia hessii* | | | 149213077 | | |
| Malpighiales | Clusiaceae | *Clusia gundlachii* | Z75673 | AY788209 | EF135520 | AY674584 | |
| Malpighiales | Bonnetiaceae | *Archytaea multiflora* | AY380342 | AY788202 | | AY674574 | |
| Malpighiales | Bonnetiaceae | *Archytaea triflora* | | | HQ331545.1 | | |
| Malpighiales | Bonnetiaceae | *Bonnetia roraimae* | AJ402930 | | | | |
| Malpighiales | Bonnetiaceae | *Bonnetia sessilis* | | 257853479 | EF135509 | FJ707523.1 | |
| Malpighiales | Calophyllaceae | *Mesua sp.* | AF206794 | AF209627 | EF135567 | AF206962 | |
| Malpighiales | Calophyllaceae | *Calophyllum sp.* | Z75672 | | | | |
| Malpighiales | Calophyllaceae | *Calophyllum leleanii* | | | 47570919 | | |
| Malpighiales | Calophyllaceae | *Calophyllum soulattri* | | | | AY674580 | |
| Malpighiales | Calophyllaceae | *Mammea americana* | AY625029 | | 47570933 | | |
| Malpighiales | Calophyllaceae | *Mammea siamensis* | | 257852931 | | FJ669689.1 | |
| Malpighiales | Podostemaceae | *Podostemum ceratophyllum* | U68088 | AY788249 | EF135584 | AY674630 | |
| Malpighiales | Podostemaceae | *Marathrum rubrum* | U68085 | AF209624 | | AF206958 | |
| Malpighiales | Podostemaceae | *Marathrum cf.oxycarpum* | | 257852965 | | | |
| Malpighiales | Podostemaceae | *Marathrum schiedeanum* | | | AB038195 | | |
| Malpighiales | Hypericaceae | *Vismia baccifera* | 257783267 | | | | |
| Malpighiales | Hypericaceae | *Vismia rubescens* | | FJ669978.1 | | | |
| Malpighiales | Hypericaceae | *Vismia sp.* | | | EF135601.1 | AY674640.1 | |
| Malpighiales | Hypericaceae | *Hypericum perforatum* | HQ332081.1 | AF209602.1 | HQ331630.1, DQ168438 | AF206934.1 | AF479122.1 |
| Malpighiales | Hypericaceae | *Eliea articulata* | FJ670167 | FJ669976 | FJ670023 | FJ669698 | |
| Malpighiales | Hypericaceae | *Cratoxylum cochinchinense* | AB233891.1 | AB233683.1 | HQ331587.1 | AB233579.1 | |

| | | | | | | | |
|---|---|---|---|---|---|---|---|
| Malpighiales | Hypericaceae | *Cratoxylum ligustrinum* | | | 222079602 | | |
| Malpighiales | Achariaceae | *Erythrospermum phytolaccoides* | AJ418798 | 257852923 | EF135535 | FJ669684.1 | |
| Malpighiales | Achariaceae | *Hydnocarpus heterophylla* | AF206778 | AF209600 | | AF206932 | AF479121 |
| Malpighiales | Achariaceae | *Hydnocarpus sp.* | | | EF135551 | | |
| Malpighiales | Achariaceae | *Pangium edule* | AF206801 | AF209644 | FJ669998.1 | AF206979 | |
| Malpighiales | Achariaceae | *Kiggelaria africana* | AF206786 | | EF135555 | AF206945 | |
| Malpighiales | Achariaceae | *Kiggelaria sp.* | | AY788231 | | | |
| Malpighiales | Achariaceae | *Acharia tragodes* | AJ418795 | AF209520 | EF135500 | AF206837 | |
| Malpighiales | Passifloraceae | *Malesherbia linearifolia* | AF206792 | AF209622 | EF135562 | AF206957 | DQ123011 |
| Malpighiales | Passifloraceae | *Turnera subulata* | DQ123398 | | | | DQ123012 |
| Malpighiales | Passifloraceae | *Turnera ulmifolia* | | AJ235634 | EF135599 | U42817 | |
| Malpighiales | Passifloraceae | *Paropsia madagascariensis* | AF206802 | | EF135576 | AF206980 | |
| Malpighiales | Passifloraceae | *Passiflora coccinea* | | AJ235553 | | | |
| Malpighiales | Passifloraceae | *Passiflora biflora* | EU017122 | EU017086 | EU017067 | | |
| Malpighiales | Passifloraceae | *Passiflora foetida* | | | | GU476454 | DQ122966 |
| Malpighiales | Passifloraceae | *Passiflora standleyi* | | | | AF206981 | |
| Malpighiales | Goupiaceae | *Goupia glabra* | AJ235780 | AJ235485 | EF135544 | AF206920 | |
| Malpighiales | Violaceae | *Rinorea pubiflora* | AY935749 | AY935862 | AY935937 | AY929373 | AY935821 |
| Malpighiales | Violaceae | *Rinorea bengalensis* | | | 115112372 | | |
| Malpighiales | Violaceae | *Viola philippica* | AB354436 | | DQ842600 | 195183680 | |
| Malpighiales | Violaceae | *Viola pubescens* | | FJ669992 | | FJ669717 | EF564797 |
| Malpighiales | Violaceae | *Hymenanthera alpina* | Z75692 | AJ235499 | EF135552 | AF206933 | |
| Malpighiales | Violaceae | *Hybanthus concolor* | AY757141 | | EF135550 | | |
| Malpighiales | Violaceae | *Hybanthus elatus* | | AB354525 | | AB354525 | |
| Malpighiales | Violaceae | *Leonia glycycarpa* | 257783287 | AY788234 | EF135558 | AY674613 | |
| Malpighiales | Lacistemataceae | *Lozania pittieri* | AJ418804 | 257852955 | FJ670026.1 | FJ669702 | |
| Malpighiales | Lacistemataceae | *Lacistema aggregatum* | AF206787.1 | AF209613.1 | AY935933.1 | AF206949.1 | AY935816.1 |
| Malpighiales | Salicaceae | *Lunania sp.* | AY788182 | AY788236 | | AY674615 | |
| Malpighiales | Salicaceae | *Lunania parviflora* | | | EF135561 | | |
| Malpighiales | Salicaceae | *Casearia sylvestris* | AF206746.1 | AF209557.1 | | AF206882.1 | |
| Malpighiales | Salicaceae | *Casearia javitensis* | | | AY935927.1 | | AY935809.1 |
| Malpighiales | Salicaceae | *Scyphostegia borneensis* | AJ403000 | AY788254 | EF135594 | AY674635 | |
| Malpighiales | Salicaceae | *Poliothyrsis sp.* | AY788186 | AY788251 | | AY674631 | |
| Malpighiales | Salicaceae | *Poliothyrsis sinensis* | | | EF135586 | | |
| Malpighiales | Salicaceae | *Idesia polycarpa* | AF206781 | AF209604 | 118917619 | AB233623 | 19919558 |
| Malpighiales | Salicaceae | *Salix reticulata* | AJ235793 | AJ235590 | EF135592 | AF207011 | |
| Malpighiales | Salicaceae | *Populus tremuloides* | M58392.1 | AF209658.1 | | AF206999.1 | AF479118.1 |
| Malpighiales | Salicaceae | *Populus nigra* | | | AB038186.1 | | |
| Malpighiales | Salicaceae | *Dovyalis rhamnoides* | Z75677 | AY788217 | EF135529 | AY674592 | |
| Malpighiales | Salicaceae | *Flacourtia indica* | GU135218 | AB233725 | GU135055 | AB233621 | |
| Malpighiales | Salicaceae | *Abatia parviflora* | AF206726 | AF209519 | EF135498 | AF206836 | |
| Malpighiales | Salicaceae | *Prockia sp.* | AY788187 | AY788252 | | AY674632 | |
| Malpighiales | Salicaceae | *Prockia crucis* | | | EF135588 | | |
| Malpighiales | Elatinaceae | *Elatine triandra* | AY380349 | AY788219 | EF135532 | AY674594 | |
| Malpighiales | Elatinaceae | *Bergia texana* | AY380344 | | EF135506 | AY674577 | |
| Malpighiales | Malpighiaceae | *Byrsonima crassifolia* | L01892 | AY788206 | AF344535 | AY674579 | EU002150 |
| Malpighiales | Malpighiaceae | *Acridocarpus natalitius* | AF344455 | AY788200 | AF344525 | AY674573 | |
| Malpighiales | Malpighiaceae | *Tetrapterys glabrifolia* | AB233903 | AB233695 | AB233799 | AB233591 | |
| Malpighiales | Malpighiaceae | *Thryallis latifolia* | AF344516 | | AF344580 | | |

| | | | | | | | |
|---|---|---|---|---|---|---|---|
| Malpighiales | Malpighiaceae | *Thryallis longifolia* | | AY788258 | | AY674638 | |
| Malpighiales | Malpighiaceae | *Dicella nucifera* | AJ235802 | AJ235453 | AF344541 | AF206901 | |
| Malpighiales | Malpighiaceae | *Malpighia emarginata* | 14599571 | | 14599701 | | |
| Malpighiales | Malpighiaceae | *Malpighia coccigera* | | 8439434 | | 398579 | |
| Malpighiales | Phyllanthaceae | *Aporusa frutescens* | Z75674 | AY788201 | 51831167 | AY788147 | |
| Malpighiales | Phyllanthaceae | *Bischofia javanica* | AY663571.1 | AY830200.1 | EF135508.1 | AB233605.1 | EF135606.1 |
| Malpighiales | Phyllanthaceae | *Phyllanthus flexuosus* | AY663603 | AB233713.1 | | AB233609.1 | |
| Malpighiales | Phyllanthaceae | *Phyllanthus myrtifolius* | | | AY936616.1 | | |
| Malpighiales | Phyllanthaceae | *Phyllanthus calcinus* | | | | | EU002156.1 |
| Malpighiales | Phyllanthaceae | *Heywoodia lucens* | AY663587 | AY788224 | AM745937 | AY674602 | |
| Malpighiales | Phyllanthaceae | *Lachnostylis bilocularis* | 17066146 | AY830218 | | AY674611 | |
| Malpighiales | Phyllanthaceae | *Lachnostylis sp.* | | | 51831197 | | |
| Malpighiales | Phyllanthaceae | *Croizatia brevipetiolata* | AY663579 | AY788213 | FJ670033.1 | AY788148 | |
| Malpighiales | Picrodendraceae | *Podocalyx loranthoides* | AY663647 | AY788248 | EF135583 | AY674629 | |
| Malpighiales | Picrodendraceae | *Tetracoccus dioicus* | AY788190 | AY788256 | FJ670035.1 | AY788158 | |
| Malpighiales | Picrodendraceae | *Androstachys johnsonii* | AF206734.1 | AF209527.1 | AY552461.1 | AF206848.1 | AF479123.1 |
| Malpighiales | Picrodendraceae | *Petalostigma pubescens* | AY380357 | AY788245 | EF135579 | AY674626 | |
| Malpighiales | Picrodendraceae | *Dissiliaria muelleri* | AY380346 | 257853487 | EF135528 | FJ669707.1 | |
| Malpighiales | Picrodendraceae | *Micrantheum hexandrum* | AJ418816 | AY788237 | EF135569 | AY674617 | |
| Malpighiales | Picrodendraceae | *Austrobuxus megacarpus* | AY380343 | AY788204 | EF135504 | AY674576 | |
| Malpighiales | Putranjivaceae | *Drypetes roxburghii* | M95757 | | | U42534 | |
| Malpighiales | Putranjivaceae | *Drypetes madagascariensis* | | AY830256 | AY552457 | | |
| Malpighiales | Lohopyxidaceae | *Lophopyxis maingayi* | AY663643 | AY788235 | EF135560 | AY674614 | |
| Malpighiales | Ctenolophonaceae | *Ctenolophon englerianus* | AJ402940 | AY788215 | EF135524 | AY674589 | |
| Malpighiales | Erythroxylaceae | *Erythroxylum confusum* | L13183 | AJ235466 | | AF206909 | |
| Malpighiales | Erythroxylaceae | *Erythroxylum coca* | | | EF135536 | | EF135611 |
| Malpighiales | Erythroxylaceae | *Aneulophus africanus* | FJ670166 | FJ669973 | FJ670010 | FJ669694 | |
| Malpighiales | Rhizophoraceae | *Paradrypetes subintegrifolia* | 257783279 | AY788243 | FJ670039.1 | AY788154 | |
| Malpighiales | Rhizophoraceae | *Cassipourea lanceolata* | FJ670174 | FJ669988 | FJ670038 | FJ669713 | |
| Malpighiales | Rhizophoraceae | *Crossostylis grandiflora* | AF006760 | AB233721 | AB233825 | AB233617 | |
| Malpighiales | Rhizophoraceae | *Rhizophora stylosa* | AF127686 | | AF105092 | AY289627 | |
| Malpighiales | Rhizophoraceae | *Rhizophora racemosa* | | | | | AF224693 |
| Malpighiales | Rhizophoraceae | *Bruguiera gymnorhiza* | U26320 | AF209547 | EF135511 | AF206875 | |
| Malpighiales | Rhizophoraceae | *Carallia sp.* | AF206744 | | | | |
| Malpighiales | Rhizophoraceae | *Carallia brachiata* | | AJ235425 | EF135513 | 257853458 | |
| Malpighiales | Pandaceae | *Microdesmis pierlotiana* | AY663645 | | | AY674618 | |
| Malpighiales | Pandaceae | *Microdesmis puberula* | | AY788238 | EF135570 | | |
| Malpighiales | Pandaceae | *Panda oleosa* | AY663644 | AY788242 | FJ670032.1 | AY788153 | |
| Malpighiales | Pandaceae | *Galearia filiformis* | AJ418818 | AY788222 | EF135542 | AY674598 | |

| | | | | | | | |
|---|---|---|---|---|---|---|---|
| Malpighiales | Centroplacaceae | *Centroplacus glaucinus* | AY663646 | AY788207 | FJ670002.1 | AY674582 | |
| Malpighiales | Centroplacaceae | *Bhesa paniculata* | AY935722.1 | AY935825.1 | AY935897.1 | AY929340.1 | AY935793.1 |
| Malpighiales | Linaceae | *Reinwardtia indica* | L13188.2 | AJ235577.2 | AB048380.1 | AF207005.1 | AF479124.1 |
| Malpighiales | Linaceae | *Linum arboreum* | AY380351 | | | | |
| Malpighiales | Linaceae | *Linum perenne* | | AJ235521 | AB038182 | L24401 | |
| Malpighiales | Linaceae | *Linum usitatissimum* | | | | | EU307117.1 |
| Malpighiales | Linaceae | *Durandea pentagyna* | AY788173 | AY788218 | FJ670027.1 | AY788150 | |
| Malpighiales | Linaceae | *Hugonia platysepala* | Z75678 | | | 51320376 | |
| Malpighiales | Linaceae | *Hugonia busseana* | | | 219551909 | | |
| Malpighiales | Balanopaceae | *Balanops vieillardi* | AF206738 | AF209534 | EF135505 | AF206860 | |
| Malpighiales | Trigoniaceae | *Trigonia nivea* | AF206830 | | EF135598 | AF207047 | |
| Malpighiales | Trigoniaceae | *Trigonia boliviana* | | AB233640 | | | |
| Malpighiales | Dichapetalaceae | *Tapura fischeri* | GQ424471 | | | | |
| Malpighiales | Dichapetalaceae | *Tapura guianensis* | | 257852939 | FJ670009.1 | FJ669693.1 | |
| Malpighiales | Dichapetalaceae | *Dichapetalum rugosum* | GQ424469 | | | GQ424451 | |
| Malpighiales | Dichapetalaceae | *Dichapetalum brownii* | | AJ235455 | | | |
| Malpighiales | Dichapetalaceae | *Dichapetalum macrocarpum* | | | EF135527 | | |
| Malpighiales | Euphroniaceae | *Euphronia guianensis* | GQ424470 | AY788221 | EF135540 | AY674597 | |
| Malpighiales | Chrysobalanaceae | *Hirtella bicornis* | AF089756 | AY788225 | FJ670003.1 | AY674603 | |
| Malpighiales | Chrysobalanaceae | *Licania elaeosperma* | AB233846 | AB233638 | AB233742 | AB233534 | |
| Malpighiales | Chrysobalanaceae | *Licania heteromorpha* | | | | | AF222370 |
| Malpighiales | Chrysobalanaceae | *Atuna racemosa* | AF089758 | AY788203 | EF135503 | AY674575 | |
| Malpighiales | Chrysobalanaceae | *Chrysobalanus icaco* | GQ424476 | AF209562 | EF135519 | U42519 | AF479119 |
| Malpighiales | Caryocaraceae | *Caryocar glabrum* | Z75671 | AF209556 | EF135515 | AF206881 | |
| Malpighiales | Ochnaceae | *Medusagyne oppositifolia* | Z75670 | AJ235530 | FJ670030 | AF206959 | AF479120 |
| Malpighiales | Ochnaceae | *Touroulia guianensis* | Z75690 | 257852969 | FJ670037.1 | FJ669710.1 | |
| Malpighiales | Ochnaceae | *Quiina pteridophylla* | AF206815 | AF209664 | EF135589 | AF207003 | |
| Malpighiales | Ochnaceae | *Luxemburgia ciliosa* | Z75685 | | | | |
| Malpighiales | Ochnaceae | *Luxemburgia octandra* | | 257852961 | | FJ669705.1 | |
| Malpighiales | Ochnaceae | *Ochna sp.* | AY380354 | AY788240 | | AY674620 | |
| Malpighiales | Ochnaceae | *Ochna multiflora* | | | EF135572 | | |
| Malpighiales | Ochnaceae | *Cespedesia bonplandi* | AJ420168 | AY788208 | EF135518 | AY674583 | |
| Malpighiales | Ochnaceae | *Sauvagesia africana* | AB233909 | | | | |
| Malpighiales | Ochnaceae | *Sauvagesia calophyllum* | Z75686 | | | | |
| Malpighiales | Ochnaceae | *Sauvagesia erecta* | | FJ669984 | EF135593 | EF135604 | EF135618 |
| Malpighiales | Peraceae | *Pogonophora schomburgkiana* | AY788185 | AY788250 | EF135585 | AY788156 | EF135614 |
| Malpighiales | Peraceae | *Clutia pulchella* | AM234976 | | EF135521 | | 133900035 |
| Malpighiales | Peraceae | *Clutia tomentosa* | | AY788210 | | AY674585 | |
| Malpighiales | Peraceae | *Pera bicolor* | AY794968 | AY788244 | EF135578 | AY674624 | EF135613 |
| Malpighiales | Euphorbiaceae | *Neoscortechinia kingii* | AB267912 | AB267964 | EF135571 | AB268068 | EF135612 |
| Malpighiales | Euphorbiaceae | *Endospermum moluccanum* | AJ402950 | AY788220 | EF135533 | AY674595 | 133900038 |
| Malpighiales | Euphorbiaceae | *Tetrorchidium gabonense* | AY794872 | | | | |
| Malpighiales | Euphorbiaceae | *Tetrorchidium sp.* | | AY788257 | | AY788159 | |

| | | | | | | | |
|---|---|---|---|---|---|---|---|
| Malpighiales | Euphorbiaceae | *Tetrorchidium rubrivenium* | | | 155029460 | | |
| Malpighiales | Euphorbiaceae | *Omphalea diandra* | AY788183 | AY788241 | FJ670016.1 | AY674622 | |
| Malpighiales | Euphorbiaceae | *Suregada aequoreum* | AY794867 | | | | |
| Malpighiales | Euphorbiaceae | *Suregada boiviniana* | | AY788255 | | AY788157 | |
| Malpighiales | Euphorbiaceae | *Suregada glomerulata* | | | 155029458 | | |
| Malpighiales | Euphorbiaceae | *Moultonianthus leembruggianus* | AY794982 | 257852943 | FJ670015.1 | FJ669695.1 | |
| Malpighiales | Euphorbiaceae | *Hevea brasiliensis* | AB267943 | | AB268047 | | |
| Malpighiales | Euphorbiaceae | *Hevea sp.* | | AY788223 | | AY674601 | |
| Malpighiales | Euphorbiaceae | *Manihot esculenta* | AB233880 | AB233672 | AB233776 | AB233568 | EU002153 |
| Malpighiales | Euphorbiaceae | *Croton trigonocarpus* | EF405861 | | | | |
| Malpighiales | Euphorbiaceae | *Croton alabamensis* | | AY788214 | | AY674588 | |
| Malpighiales | Euphorbiaceae | *Croton kongensis* | | | GQ434077 | | |
| Malpighiales | Euphorbiaceae | *Codiaeum peltatum* | AB233876 | AB233668 | | | |
| Malpighiales | Euphorbiaceae | *Codiaeum variegatum* | | | EF135522 | AY674586 | EF135608.1 |
| Malpighiales | Euphorbiaceae | *Trigonostemon verrucosus* | AY788192 | AY788259 | FJ670020.1 | AY788160 | |
| Malpighiales | Euphorbiaceae | *Pimelodendron zoanthogyne* | AJ418812 | AY788247 | EF135582 | AY674628 | |
| Malpighiales | Euphorbiaceae | *Euphorbia polychroma* | AY794827 | AJ235472 | EF135539 | | AF479125 |
| Malpighiales | Euphorbiaceae | *Euphorbia pulcherrima* | | | | L37582 | |
| Malpighiales | Euphorbiaceae | *Omalanthus populneus* | AJ402978 | | | | |
| Malpighiales | Euphorbiaceae | *Homalanthus populneus* | | AY788226 | EF135548 | AY674604 | |
| Malpighiales | Euphorbiaceae | *Hura crepitans* | GQ981765 | AY788228 | 118917521 | AY674606 | |
| Malpighiales | Euphorbiaceae | *Conceveiba martiana* | AY788170 | AY788212 | FJ670011.1 | | |
| Malpighiales | Euphorbiaceae | *Lasiocroton bahamensis* | AY788181 | AY788233 | | AY788152 | |
| Malpighiales | Euphorbiaceae | *Lasiocroton macrophyllus* | | | 155029394 | | |
| Malpighiales | Euphorbiaceae | *Dalechampia spathulata* | AY788172 | AY788216 | EF135525 | AY788149 | 133900037 |
| Malpighiales | Euphorbiaceae | *Ricinus communis* | AY788188 | AY788253 | EF135590 | AY674633 | |
| Malpighiales | Euphorbiaceae | *Spathiostemon javensis* | AY788176 | AY788227 | FJ670017.1 | AY788151 | |
| Malpighiales | Euphorbiaceae | *Acalypha californica* | AY380341 | AY788199 | EF135499 | AY674572 | EF135605 |

**Supplementary Table 2.** Sampling fraction associated to each terminal. Sampling fraction associated to each species included as a terminal in diversification analyses. Each sampling fraction is derived from the total number of living species in the clade represented by each terminal, and the number of representatives for that clade. Clades correspond to a single, or two or more families.

| Order | Family | Clade | Terminal | Number of species in clade | Number of sampled species | Sampling fraction |
|---|---|---|---|---|---|---|
| Amborellales | Amborellaceae | Amborellaceae | *Amborella_trichopoda* | 1 | 1 | 1.000000 |
| Nymphaeales | Hydatellaceae | Hydatellaceae | *Trithuria_submersa* | 10 | 1 | 0.100000 |
| Nymphaeales | Nymphaeaceae | Nymphaeaceae | *Brasenia_schreberi* | 64 | 4 | 0.062500 |
| Nymphaeales | Nymphaeaceae | Nymphaeaceae | *Cabomba_caroliniana* | 64 | 4 | 0.062500 |
| Nymphaeales | Nymphaeaceae | Nymphaeaceae | *Nuphar_spp.* | 64 | 4 | 0.062500 |
| Nymphaeales | Nymphaeaceae | Nymphaeaceae | *Nymphaea_spp.* | 64 | 4 | 0.062500 |
| Austrobaileyales | Austrobaileyaceae | Austrobaileyaceae | *Austrobaileya_scandens* | 2 | 1 | 0.500000 |
| Austrobaileyales | Trimeniaceae | Trimeniaceae | *Trimenia_moorei* | 6 | 1 | 0.166667 |
| Austrobaileyales | Schisandraceae | Schisandraceae | *Illicium_spp.* | 92 | 3 | 0.032609 |
| Austrobaileyales | Schisandraceae | Schisandraceae | *Kadsura_japonica* | 92 | 3 | 0.032609 |
| Austrobaileyales | Schisandraceae | Schisandraceae | *Schisandra_spp.* | 92 | 3 | 0.032609 |
| Chloranthales | Chloranthaceae | Chloranthaceae | *Hedyosmum_arborescens* | 75 | 4 | 0.053333 |
| Chloranthales | Chloranthaceae | Chloranthaceae | *Ascarina* | 75 | 4 | 0.053333 |
| Chloranthales | Chloranthaceae | Chloranthaceae | *Chloranthus_spp.* | 75 | 4 | 0.053333 |
| Chloranthales | Chloranthaceae | Chloranthaceae | *Sarcandra_spp.* | 75 | 4 | 0.053333 |
| Canellales | Canellaceae | Canellaceae | *Canella_winterana* | 13 | 2 | 0.153846 |
| Canellales | Canellaceae | Canellaceae | *Cinnamodendron_spp.* | 13 | 2 | 0.153846 |
| Canellales | Winteraceae | Winteraceae | *Takhtajania_perrieri* | 90 | 3 | 0.033333 |
| Canellales | Winteraceae | Winteraceae | *Drimys_spp.* | 90 | 3 | 0.033333 |
| Canellales | Winteraceae | Winteraceae | *Tasmannia_spp.* | 90 | 3 | 0.033333 |
| Piperales | Piperaceae | Piperaceae | *Peperomia_spp.* | 3615 | 2 | 0.000553 |
| Piperales | Piperaceae | Piperaceae | *Piper_spp.* | 3615 | 2 | 0.000553 |
| Piperales | Saururaceae | Saururaceae | *Houttuynia_cordata* | 6 | 3 | 0.500000 |
| Piperales | Saururaceae | Saururaceae | *Anemopsis_californica* | 6 | 3 | 0.500000 |
| Piperales | Saururaceae | Saururaceae | *Saururus_spp.* | 6 | 3 | 0.500000 |
| Piperales | Aristolochiaceae | Aristolochiaceae plus Hydnoraceae | *Asarum_spp.* | 487 | 5 | 0.010267 |
| Piperales | Aristolochiaceae | Aristolochiaceae plus Hydnoraceae | *Saruma* | 487 | 5 | 0.010267 |
| Piperales | Aristolochiaceae | Aristolochiaceae plus Hydnoraceae | *Lactoris_fernandeziana* | 487 | 5 | 0.010267 |

| | | | | | | |
|---|---|---|---|---|---|---|
| Piperales | Aristolochiaceae | Aristolochiaceae plus Hydnoraceae | *Aristolochia_macrophylla* | 487 | 5 | 0.010267 |
| Piperales | Aristolochiaceae | Aristolochiaceae plus Hydnoraceae | *Thottea_tomentosa* | 487 | 5 | 0.010267 |
| Magnoliales | Himantandraceae | Himantandraceae | *Galbulimima_belgraveana* | 2 | 1 | 0.500000 |
| Magnoliales | Eupomatiaceae | Eupomatiaceae | *Eupomatia_bennettii* | 3 | 1 | 0.333333 |
| Magnoliales | Annonaceae | Annonaceae | *Cananga_odorata* | 2220 | 3 | 0.001351 |
| Magnoliales | Annonaceae | Annonaceae | *Annona_muricata* | 2220 | 3 | 0.001351 |
| Magnoliales | Annonaceae | Annonaceae | *Asimina_triloba* | 2220 | 3 | 0.001351 |
| Magnoliales | Magnoliaceae | Magnoliaceae | *Liriodendron_spp.* | 227 | 2 | 0.008811 |
| Magnoliales | Magnoliaceae | Magnoliaceae | *Magnolia_spp.* | 227 | 2 | 0.008811 |
| Magnoliales | Degeneriaceae | Degeneriaceae | *Degeneria_vitiensis* | 2 | 1 | 0.500000 |
| Magnoliales | Myristicaceae | Myristicaceae | *Mauloutchia_chapelieri* | 475 | 2 | 0.004211 |
| Magnoliales | Myristicaceae | Myristicaceae | *Myristica_fragrans* | 475 | 2 | 0.004211 |
| Laurales | Calycanthaceae | Calycanthaceae | *Idiospermum_australiense* | 11 | 3 | 0.272727 |
| Laurales | Calycanthaceae | Calycanthaceae | *Calycanthus_spp.* | 11 | 3 | 0.272727 |
| Laurales | Calycanthaceae | Calycanthaceae | *Chimonanthus_praecox* | 11 | 3 | 0.272727 |
| Laurales | Siparunaceae | Siparunaceae | *Siparuna_spp.* | 75 | 1 | 0.013333 |
| Laurales | Gomortegaceae | Gomortegaceae | *Gomortega_keule* | 1 | 1 | 1.000000 |
| Laurales | Atherospermataceae | Atherospermataceae | *Doryphora_spp.* | 16 | 3 | 0.187500 |
| Laurales | Atherospermataceae | Atherospermataceae | *Atherosperma* | 16 | 3 | 0.187500 |
| Laurales | Atherospermataceae | Atherospermataceae | *Daphnandra* | 16 | 3 | 0.187500 |
| Laurales | Lauraceae | Lauraceae | *Cryptocarya_spp.* | 2500 | 4 | 0.001600 |
| Laurales | Lauraceae | Lauraceae | *Cinnamomum_camphora* | 2500 | 4 | 0.001600 |
| Laurales | Lauraceae | Lauraceae | *Laurus_nobilis* | 2500 | 4 | 0.001600 |
| Laurales | Lauraceae | Lauraceae | *Sassafras_spp.* | 2500 | 4 | 0.001600 |
| Laurales | Hernandiaceae | Hernandiaceae | *Gyrocarpus_spp.* | 55 | 2 | 0.036364 |
| Laurales | Hernandiaceae | Hernandiaceae | *Hernandia_spp.* | 55 | 2 | 0.036364 |
| Laurales | Monimiaceae | Monimiaceae | *Peumus_boldus* | 200 | 3 | 0.015000 |
| Laurales | Monimiaceae | Monimiaceae | *Hedycarya_arborea* | 200 | 3 | 0.015000 |
| Laurales | Monimiaceae | Monimiaceae | *Hortonia_floribunda* | 200 | 3 | 0.015000 |
| Acorales | Acoraceae | Acoraceae | *Acorus_gramineus* | 4 | 1 | 0.250000 |
| Alismatales | Araceae | Araceae | *Orontium_aquaticum* | 4095 | 4 | 0.000977 |
| Alismatales | Araceae | Araceae | *Lemna* | 4095 | 4 | 0.000977 |
| Alismatales | Araceae | Araceae | *Spathiphyllum_spp.* | 4095 | 4 | 0.000977 |
| Alismatales | Araceae | Araceae | *Xanthosoma* | 4095 | 4 | 0.000977 |
| Alismatales | Tofieldiaceae | Tofieldiaceae | *Pleea_tenuifolia* | 31 | 2 | 0.064516 |
| Alismatales | Tofieldiaceae | Tofieldiaceae | *Tofieldia_calyculata* | 31 | 2 | 0.064516 |
| Alismatales | Potamogetonaceae | Potamogetonaceae plus Zosteraceae plus Cymodoceaceae plus Ruppiaceae | *Potamogeton_spp.* | 152 | 1 | 0.006579 |

| | | | | | | |
|---|---|---|---|---|---|---|
| | | plus Posidoniaceae plus Maundiaceae | | | | |
| Alismatales | Juncaginaceae | Juncaginaceae plus Scheuchzeriaceae plus Aponogetonaceae | *Triglochin_maritima* | 59 | 1 | 0.016949 |
| Alismatales | Alismataceae | Alismataceae | *Alisma* | 88 | 1 | 0.011364 |
| Alismatales | Hydrocharitaceae | Hydrocharitaceae plus Butomaceae | *Hydrocharis_spp.* | 117 | 2 | 0.017094 |
| Alismatales | Hydrocharitaceae | Hydrocharitaceae plus Butomaceae | *Najas_spp.* | 117 | 2 | 0.017094 |
| Petrosaviales | Petrosaviaceae | Petrosaviaceae | *Japonolirion_osense* | 4 | 2 | 0.500000 |
| Petrosaviales | Petrosaviaceae | Petrosaviaceae | *Petrosavia_spp.* | 4 | 2 | 0.500000 |
| Dioscoreales | Nartheciaceae | Nartheciaceae | *Metanarthecium_luteoviride* | 41 | 1 | 0.024390 |
| Dioscoreales | Taccaceae | Taccaceae plus Thismiaceae | *Tacca_spp.* | 67 | 1 | 0.014925 |
| Dioscoreales | Burmanniaceae | Burmanniaceae | *Burmannia_spp.* | 95 | 1 | 0.010526 |
| Dioscoreales | Dioscoreaceae | Dioscoreaceae | *Dioscorea_spp.* | 870 | 1 | 0.001149 |
| Pandanales | Cyclanthaceae | Cyclanthaceae plus Pandanaceae | *Carludovica_palmata* | 1110 | 1 | 0.000901 |
| Pandanales | Stemonaceae | Stemonaceae | *Croomia_pauciflora* | 27 | 1 | 0.037037 |
| Pandanales | Velloziaceae | Velloziaceae plus Triuridaceae | *Acanthochlamys_bracteata* | 290 | 3 | 0.010345 |
| Pandanales | Velloziaceae | Velloziaceae plus Triuridaceae | *Barbacenia_elegans* | 290 | 3 | 0.010345 |
| Pandanales | Velloziaceae | Velloziaceae plus Triuridaceae | *Xerophyta_spp.* | 290 | 3 | 0.010345 |
| Liliales | Campynemataceae | Campynemataceae plus Corsiaceae | *Campynema_linearis* | 34 | 1 | 0.029412 |
| Liliales | Melanthiaceae | Melanthiaceae | *Trillium_spp.* | 170 | 1 | 0.005882 |
| Liliales | Colchicaceae | Colchicaceae plus Petermaniaceae | *Colchicum_spp.* | 246 | 1 | 0.004065 |
| Liliales | Alstroemeriaceae | Alstroemeriaceae plus Luzuriagaceae | *Alstroemeria_spp.* | 170 | 2 | 0.011765 |
| Liliales | Alstroemeriaceae | Alstroemeriaceae plus Luzuriagaceae | *Bomarea_spp.* | 170 | 2 | 0.011765 |

| | | | | | | |
|---|---|---|---|---|---|---|
| Liliales | Philesiaceae | Philesiaceae plus Rhipogonaceae | *Lapageria_rosea* | 8 | 2 | 0.250000 |
| Liliales | Philesiaceae | Philesiaceae plus Rhipogonaceae | *Philesia_buxifolia* | 8 | 2 | 0.250000 |
| Liliales | Liliaceae | Liliaceae | *Lilium_spp.* | 610 | 1 | 0.001639 |
| Liliales | Smilacaceae | Smilacaceae | *Smilax_glauca* | 315 | 1 | 0.003175 |
| Asparagales | Orchidaceae | Orchidaceae | *Cypripedium_spp.* | 22075 | 3 | 0.000136 |
| Asparagales | Orchidaceae | Orchidaceae | *Oncidium* | 22075 | 3 | 0.000136 |
| Asparagales | Orchidaceae | Orchidaceae | *Phalaenopsis* | 22075 | 3 | 0.000136 |
| Asparagales | Boryaceae | Boryaceae | *Borya_septentrionalis* | 12 | 1 | 0.083333 |
| Asparagales | Blandfordiaceae | Blandfordiaceae | *Blandfordia* | 4 | 1 | 0.250000 |
| Asparagales | Hypoxidaceae | Hypoxidaceae plus Asteliaceae plus Lanariaceae | *Molineria_capitulata* | 257 | 3 | 0.011673 |
| Asparagales | Hypoxidaceae | Hypoxidaceae plus Asteliaceae plus Lanariaceae | *Hypoxis_spp.* | 257 | 3 | 0.011673 |
| Asparagales | Hypoxidaceae | Hypoxidaceae plus Asteliaceae plus Lanariaceae | *Rhodohypoxis_millioides* | 257 | 3 | 0.011673 |
| Asparagales | Ixioliriaceae | Ixolirionaceae | *Ixiolirion_tataricum* | 3 | 1 | 0.333333 |
| Asparagales | Tecophilaeaceae | Tecophilaceae | *Tecophilaea_cyanocrocus* | 25 | 1 | 0.040000 |
| Asparagales | Iridaceae | Iridaceae plus Dorianthaceae | *Alophia_drummondii* | 2037 | 4 | 0.001964 |
| Asparagales | Iridaceae | Iridaceae plus Dorianthaceae | *Iris* | 2037 | 4 | 0.001964 |
| Asparagales | Iridaceae | Iridaceae plus Dorianthaceae | *Aristea_glauca* | 2037 | 4 | 0.001964 |
| Asparagales | Iridaceae | Iridaceae plus Dorianthaceae | *Gladiolus_buckerveldii* | 2037 | 4 | 0.001964 |
| Asparagales | Asphodelaceae | Asphodelaceae | *Bulbine_spp.* | 785 | 1 | 0.001274 |
| Asparagales | Xanthorrhoeaceae | Xanthorrhoeaceae plus Xeronemataceae | *Xanthorrhoea* | 117 | 1 | 0.008547 |
| Asparagales | Amaryllidaceae | Amaryllidaceae | *Crinum_spp.* | 1605 | 3 | 0.001869 |
| Asparagales | Amaryllidaceae | Amaryllidaceae | *Allium* | 1605 | 3 | 0.001869 |
| Asparagales | Amaryllidaceae | Amaryllidaceae | *Nothoscordum_spp.* | 1605 | 3 | 0.001869 |
| Asparagales | Asparagaceae | Asparagaceae | *Asparagus_spp.* | 2480 | 5 | 0.002016 |
| Asparagales | Asparagaceae | Asparagaceae | *Agave* | 2480 | 5 | 0.002016 |
| Asparagales | Asparagaceae | Asparagaceae | *Yucca* | 2480 | 5 | 0.002016 |
| Asparagales | Asparagaceae | Asparagaceae | *Beaucarnea* | 2480 | 5 | 0.002016 |
| Asparagales | Asparagaceae | Asparagaceae | *Lomandra* | 2480 | 5 | 0.002016 |
| unplaced Dasypogonaceae | Dasypogonaceae | Dasypogonaceae | *Calectasia_spp.* | 16 | 2 | 0.125000 |
| unplaced Dasypogonaceae | Dasypogonaceae | Dasypogonaceae | *Dasypogon_bromeliifolius* | 16 | 2 | 0.125000 |
| Arecales | Arecaceae | Arecaceae | *Chamaedorea* | 2361 | 5 | 0.002118 |

| | | | | | | |
|---|---|---|---|---|---|---|
| Arecales | Arecaceae | Arecaceae | *Elaeis* | 2361 | 5 | 0.002118 |
| Arecales | Arecaceae | Arecaceae | *Caryota_mitis* | 2361 | 5 | 0.002118 |
| Arecales | Arecaceae | Arecaceae | *Phoenix_dactylifera* | 2361 | 5 | 0.002118 |
| Arecales | Arecaceae | Arecaceae | *Trachycarpus_fortunei* | 2361 | 5 | 0.002118 |
| Commelinales | Commelinaceae | Commelinaceae plus Hanguanaceae | *Tradescantia_spp.* | 662 | 1 | 0.001511 |
| Commelinales | Philydraceae | Philydraceae | *Philydrum_lanuginosum* | 5 | 1 | 0.200000 |
| Commelinales | Haemodoraceae | Haemodoraceae | *Anigozanthos_flavidus* | 116 | 1 | 0.008621 |
| Commelinales | Pontederiaceae | Pontederiaceae | *Pontederia_cordata* | 33 | 1 | 0.030303 |
| Zingiberales | Strelitziaceae | Strelitiziaceae plus Lowiaceae | *Strelitzia* | 22 | 3 | 0.136364 |
| Zingiberales | Strelitziaceae | Strelitiziaceae plus Lowiaceae | *Phenakospermum_guyannense* | 22 | 3 | 0.136364 |
| Zingiberales | Strelitziaceae | Strelitiziaceae plus Lowiaceae | *Ravenala_madagascariensis* | 22 | 3 | 0.136364 |
| Zingiberales | Musaceae | Musaceae plus Heliconiaceae | *Musa_spp.* | 241 | 1 | 0.004149 |
| Zingiberales | Zingiberaceae | Zingiberaceae plus Costaceae | *Zingiber_spp.* | 1450 | 1 | 0.000690 |
| Zingiberales | Cannaceae | Cannaceae | *Canna_spp.* | 10 | 1 | 0.100000 |
| Zingiberales | Marantaceae | Marantaceae | *Maranta_bicolor* | 550 | 1 | 0.001818 |
| Poales | Bromeliaceae | Bromeliaceae | *Puya_raimondii* | 1770 | 2 | 0.001130 |
| Poales | Bromeliaceae | Bromeliaceae | *Vriesea* | 1770 | 2 | 0.001130 |
| Poales | Typhaceae | Typhaceae | *Sparganium* | 25 | 2 | 0.080000 |
| Poales | Typhaceae | Typhaceae | *Typha_spp.* | 25 | 2 | 0.080000 |
| Poales | Rapateaceae | Rapateaceae | *Stegolepis* | 94 | 1 | 0.010638 |
| Poales | Eriocaulaceae | Eriocaulaceae | *Eriocaulon_spp.* | 1160 | 1 | 0.000862 |
| Poales | Xyridaceae | Xyridaceae | *Xyris_spp.* | 260 | 1 | 0.003846 |
| Poales | Mayacaceae | Mayacaceae | *Mayaca_spp.* | 10 | 1 | 0.100000 |
| Poales | Juncaceae | Juncaceae plus Thurniaceae | *Juncus_effusus* | 434 | 1 | 0.002304 |
| Poales | Cyperaceae | Cyperaceae | *Cyperus_spp.* | 5430 | 2 | 0.000368 |
| Poales | Cyperaceae | Cyperaceae | *Rhynchospora_spp.* | 5430 | 2 | 0.000368 |
| Poales | Anarthriaceae | Anarthriaceae | *Anarthria_spp.* | 11 | 1 | 0.090909 |
| Poales | Centrolepidaceae | Centrolepidaceae | *Centrolepis_strigosa* | 35 | 1 | 0.028571 |
| Poales | Restionaceae | Restionaceae | *Restio_spp.* | 500 | 1 | 0.002000 |
| Poales | Flagellariaceae | Flagellariaceae | *Flagellaria_indica* | 4 | 1 | 0.250000 |
| Poales | Ecdeiocoleaceae | Ecdeiocoleaceae | *Ecdeiocolea_monostachya* | 3 | 1 | 0.333333 |
| Poales | Joinvilleaceae | Joinvilleaceae | *Joinvillea_spp.* | 2 | 1 | 0.500000 |
| Poales | Poaceae | Poaceae | *Oryza_sativa* | 11337 | 5 | 0.000441 |
| Poales | Poaceae | Poaceae | *Hordeum* | 11337 | 5 | 0.000441 |
| Poales | Poaceae | Poaceae | *Danthonia_spp.* | 11337 | 5 | 0.000441 |
| Poales | Poaceae | Poaceae | *Saccharum* | 11337 | 5 | 0.000441 |
| Poales | Poaceae | Poaceae | *Zea* | 11337 | 5 | 0.000441 |
| Ceratophyllales | Ceratophyllaceae | Ceratophyllaceae | *Ceratophyllum_demersum* | 6 | 1 | 0.166667 |
| Ranunculales | Eupteleaceae | Eupteleaceae | *Euptelea_polyandra* | 2 | 1 | 0.500000 |
| Ranunculales | Papaveraceae | Papaveraceae | *Eschscholzia* | 760 | 3 | 0.003947 |
| Ranunculales | Papaveraceae | Papaveraceae | *Dicentra_spp.* | 760 | 3 | 0.003947 |

| | | | | | | |
|---|---|---|---|---|---|---|
| Ranunculales | Papaveraceae | Papaveraceae | *Hypecoum* | 760 | 3 | 0.003947 |
| Ranunculales | Circaeasteraceae | Circaeasteraceae | *Circaeaster_agrestis* | 2 | 2 | 1.000000 |
| Ranunculales | Circaeasteraceae | Circaeasteraceae | *Kingdonia_uniflora* | 2 | 2 | 1.000000 |
| Ranunculales | Lardizabalaceae | Lardizabalaceae | *Akebia_quinata* | 40 | 4 | 0.100000 |
| Ranunculales | Lardizabalaceae | Lardizabalaceae | *Lardizabala_biternata* | 40 | 4 | 0.100000 |
| Ranunculales | Lardizabalaceae | Lardizabalaceae | *Decaisnea* | 40 | 4 | 0.100000 |
| Ranunculales | Lardizabalaceae | Lardizabalaceae | *Sargentodoxa* | 40 | 4 | 0.100000 |
| Ranunculales | Menispermaceae | Menispermaceae | *Cissampelos_pareira* | 442 | 4 | 0.009050 |
| Ranunculales | Menispermaceae | Menispermaceae | *Cocculus_spp.* | 442 | 4 | 0.009050 |
| Ranunculales | Menispermaceae | Menispermaceae | *Menispermum* | 442 | 4 | 0.009050 |
| Ranunculales | Menispermaceae | Menispermaceae | *Tinospora* | 442 | 4 | 0.009050 |
| Ranunculales | Berberidaceae | Berberidaceae | *Mahonia_spp.* | 701 | 4 | 0.005706 |
| Ranunculales | Berberidaceae | Berberidaceae | *Podophyllum_peltatum* | 701 | 4 | 0.005706 |
| Ranunculales | Berberidaceae | Berberidaceae | *Caulophyllum_thalictroides* | 701 | 4 | 0.005706 |
| Ranunculales | Berberidaceae | Berberidaceae | *Nandina_domestica* | 701 | 4 | 0.005706 |
| Ranunculales | Ranunculaceae | Ranunculaceae | *Glaucidium_palmatum* | 2525 | 4 | 0.001584 |
| Ranunculales | Ranunculaceae | Ranunculaceae | *Hydrastis_canadensis* | 2525 | 4 | 0.001584 |
| Ranunculales | Ranunculaceae | Ranunculaceae | *Ranunculus_spp.* | 2525 | 4 | 0.001584 |
| Ranunculales | Ranunculaceae | Ranunculaceae | *Xanthorhiza_simplicissima* | 2525 | 4 | 0.001584 |
| Sabiales | Sabiaceae | Sabiaceae | *Meliosma_veitchiorum* | 100 | 2 | 0.020000 |
| Sabiales | Sabiaceae | Sabiaceae | *Sabia* | 100 | 2 | 0.020000 |
| Proteales | Nelumbonaceae | Nelumbonaceae | *Nelumbo_lutea* | 2 | 1 | 0.500000 |
| Proteales | Platanaceae | Platanaceae | *Platanus_occidentalis* | 10 | 1 | 0.100000 |
| Proteales | Proteaceae | Proteaceae | *Petrophile* | 1600 | 3 | 0.001875 |
| Proteales | Proteaceae | Proteaceae | *Grevillea_spp.* | 1600 | 3 | 0.001875 |
| Proteales | Proteaceae | Proteaceae | *Roupala* | 1600 | 3 | 0.001875 |
| Trochodendrales | Trochodendraceae | Trochodendraceae | *Tetracentron_sinense* | 2 | 2 | 1.000000 |
| Trochodendrales | Trochodendraceae | Trochodendraceae | *Trochodendron_aralioides* | 2 | 2 | 1.000000 |
| Buxales | Didymelaceae | Didymelaceae | *Didymeles_perrieri* | 2 | 1 | 0.500000 |
| Buxales | Buxaceae | Buxaceae | *Buxus_sempervirens* | 70 | 2 | 0.028571 |
| Buxales | Buxaceae | Buxaceae | *Pachysandra_spp.* | 70 | 2 | 0.028571 |
| Gunnerales | Gunneraceae | Gunneraceae | *Gunnera_spp.* | 50 | 1 | 0.020000 |
| Gunnerales | Myrothamnaceae | Myrothamnaceae | *Myrothamnus_spp.* | 2 | 1 | 0.500000 |
| Dilleniales | Dilleniaceae | Dilleniaceae | *Dillenia_spp.* | 410 | 3 | 0.007317 |
| Dilleniales | Dilleniaceae | Dilleniaceae | *Hibbertia* | 410 | 3 | 0.007317 |
| Dilleniales | Dilleniaceae | Dilleniaceae | *Tetracera_asiatica* | 410 | 3 | 0.007317 |
| Santalales | Coulaceae | Coulaceae plus Erythropalaceae plus Strombosiaceae | *Ochanostachys_amentacea* | 61 | 3 | 0.049180 |

| | | | | | | |
|---|---|---|---|---|---|---|
| Santalales | Coulaceae | Coulaceae plus Erythropalaceae plus Strombosiaceae | *Coula_edulis* | 61 | 3 | 0.049180 |
| Santalales | Coulaceae | Coulaceae plus Erythropalaceae plus Strombosiaceae | *Minquartia_guianensis* | 61 | 3 | 0.049180 |
| Santalales | Olacaceae | Olacaceae plus Ximeniaceae plus Aptandraceae | *Heisteria_spp.* | 104 | 2 | 0.019231 |
| Santalales | Olacaceae | Olacaceae plus Ximeniaceae plus Aptandraceae | *Ximenia_americana* | 104 | 2 | 0.019231 |
| Santalales | Santalaceae | Santalaceae | *Osyris_spp.* | 990 | 2 | 0.002020 |
| Santalales | Santalaceae | Santalaceae | *Santalum_album* | 990 | 2 | 0.002020 |
| Santalales | Opiliaceae | Opiliaceae plus Octoknemaceae | *Opilia_spp.* | 50 | 1 | 0.020000 |
| Santalales | Loranthaceae | Loranthaceae | *Gaiadendron* | 950 | 1 | 0.001053 |
| Santalales | Misodendraceae | Misodendraceae | *Misodendrum_spp.* | 8 | 1 | 0.125000 |
| Santalales | Schoepfiaceae | Schoepfiaceae | *Schoepfia_schreberi* | 55 | 1 | 0.018182 |
| Berberidopsidales | Aextoxicaceae | Aextoxicaceae | *Aextoxicon_punctatum* | 1 | 1 | 1.000000 |
| Berberidopsidales | Berberidopsidaceae | Berberidopsidaceae | *Berberidopsis_corallina* | 3 | 1 | 0.333333 |
| Caryophyllales | Droseraceae | Droseraceae | *Drosera_spp.* | 115 | 1 | 0.008696 |
| Caryophyllales | Nepenthaceae | Nepenthaceae | *Nepenthes_spp.* | 90 | 1 | 0.011111 |
| Caryophyllales | Drosophyllaceae | Drosophyllaceae | *Drosophyllum* | 1 | 1 | 1.000000 |
| Caryophyllales | Ancistrocladaceae | Ancistrocladaceae | *Ancistrocladus* | 12 | 1 | 0.083333 |
| Caryophyllales | Dioncophyllaceae | Dioncophyllaceae | *Triphyophyllum_peltatum* | 3 | 1 | 0.333333 |
| Caryophyllales | Frankeniaceae | Frankeniaceae | *Frankenia* | 90 | 1 | 0.011111 |
| Caryophyllales | Tamaricaceae | Tamaricaceae | *Tamarix_spp.* | 90 | 1 | 0.011111 |
| Caryophyllales | Polygonaceae | Polygonaceae | *Fagopyrum* | 1110 | 2 | 0.001802 |
| Caryophyllales | Polygonaceae | Polygonaceae | *Polygonum_spp.* | 1110 | 2 | 0.001802 |
| Caryophyllales | Plumbaginaceae | Plumbaginaceae | *Limonium* | 836 | 2 | 0.002392 |
| Caryophyllales | Plumbaginaceae | Plumbaginaceae | *Plumbago_spp.* | 836 | 2 | 0.002392 |
| Caryophyllales | Rhabdodendraceae | Rhabdodendraceae | *Rhabdodendron_spp.* | 3 | 1 | 0.333333 |
| Caryophyllales | Simmondsiaceae | Simmondsiaceae | *Simmondsia_chinensis* | 1 | 1 | 1.000000 |

| | | | | | | |
|---|---|---|---|---|---|---|
| Caryophyllales | Asteropeiaceae | Asteropeiaceae | *Asteropeia_micraster* | 8 | 1 | 0.125000 |
| Caryophyllales | Physenaceae | Physenaceae | *Physena* | 2 | 1 | 0.500000 |
| Caryophyllales | Caryophyllaceae | Caryophyllaceae | *Stellaria_spp.* | 2200 | 1 | 0.000455 |
| Caryophyllales | Achatocarpaceae | Achatocarpaceae | *Phaulothamnus* | 7 | 1 | 0.142857 |
| Caryophyllales | Amaranthaceae | Amaranthaceae | *Spinacia* | 2500 | 3 | 0.001200 |
| Caryophyllales | Amaranthaceae | Amaranthaceae | *Beta* | 2500 | 3 | 0.001200 |
| Caryophyllales | Amaranthaceae | Amaranthaceae | *Celosia_spp.* | 2500 | 3 | 0.001200 |
| Caryophyllales | Stegospermataceae | Stegnospermataceae | *Stegnosperma* | 3 | 1 | 0.333333 |
| Caryophyllales | Limeaceae | Limeaceae | *Limeum* | 21 | 1 | 0.047619 |
| Caryophyllales | Lophiocarpaceae | Lophiocarpaceae | *Corbichonia_decumbens* | 6 | 1 | 0.166667 |
| Caryophyllales | Barbeuiaceae | Barbeuiaceae | *Barbeuia_spp.* | 1 | 1 | 1.000000 |
| Caryophyllales | Aizoaceae | Aizoaceae | *Delosperma* | 2035 | 2 | 0.000983 |
| Caryophyllales | Aizoaceae | Aizoaceae | *Lampranthus* | 2035 | 2 | 0.000983 |
| Caryophyllales | Gisekiaceae | Gisekiaceae | *Gisekia* | 5 | 1 | 0.200000 |
| Caryophyllales | Phytolaccaceae | Phytolaccaceae | *Phytolacca_americana* | 65 | 2 | 0.030769 |
| Caryophyllales | Sarcobataceae | Sarcobataceae | *Sarcobatus* | 2 | 1 | 0.500000 |
| Caryophyllales | Phytolaccaceae | Phytolaccaceae | *Rivina* | 65 | 2 | 0.030769 |
| Caryophyllales | Nyctaginaceae | Nyctaginaceae | *Bougainvillea_spp.* | 395 | 2 | 0.005063 |
| Caryophyllales | Nyctaginaceae | Nyctaginaceae | *Mirabilis_jalapa* | 395 | 2 | 0.005063 |
| Caryophyllales | unassigned Hypertellis | Hypertellis | *Hypertelis_spp.* | 8 | 1 | 0.125000 |
| Caryophyllales | Molluginaceae | Molluginaceae | *Mollugo_verticillata* | 87 | 1 | 0.011494 |
| Caryophyllales | Montiaceae | Montiaceae | *Claytonia* | 77 | 1 | 0.012987 |
| Caryophyllales | Talinaceae | Talinaceae | *Talinum* | 27 | 1 | 0.037037 |
| Caryophyllales | Portulacaceae | Portulacaceae plus Anacampserotaceae | *Portulaca_spp.* | 132 | 1 | 0.007576 |
| Caryophyllales | Cactaceae | Cactaceae | *Opuntia* | 1866 | 2 | 0.001072 |

| | | | | | | |
|---|---|---|---|---|---|---|
| Caryophyllales | Cactaceae | Cactaceae | *Pereskia_aculeata* | 1866 | 2 | 0.001072 |
| Caryophyllales | Halophytaceae | Halophytaceae | *Halophytum* | 1 | 1 | 1.000000 |
| Caryophyllales | Didieraceae | Didieraceae | *Alluaudia* | 16 | 1 | 0.062500 |
| Caryophyllales | Basellaceae | Basellaceae | *Anredera_cordifolia* | 19 | 2 | 0.105263 |
| Caryophyllales | Basellaceae | Basellaceae | *Basella* | 19 | 2 | 0.105263 |
| Cornales | Hydrostachyaceae | Hydrostachyaceae | *Hydrostachys_spp.* | 20 | 1 | 0.050000 |
| Cornales | Cornaceae | Cornaceae | *Alangium_spp.* | 85 | 2 | 0.023529 |
| Cornales | Cornaceae | Cornaceae | *Cornus_mas* | 85 | 2 | 0.023529 |
| Cornales | Nyssaceae | Nyssaceae | *Camptotheca_acuminata* | 22 | 2 | 0.090909 |
| Cornales | Nyssaceae | Nyssaceae | *Nyssa_spp.* | 22 | 2 | 0.090909 |
| Cornales | Curtisiaceae | Curtisiaceae | *Curtisia_dentata* | 1 | 1 | 1.000000 |
| Cornales | Grubbiaceae | Grubbiaceae | *Grubbia_spp.* | 3 | 1 | 0.333333 |
| Cornales | Loasaceae | Loasaceae | *Eucnide_spp.* | 265 | 3 | 0.011321 |
| Cornales | Loasaceae | Loasaceae | *Mentzelia* | 265 | 3 | 0.011321 |
| Cornales | Loasaceae | Loasaceae | *Petalonyx_spp.* | 265 | 3 | 0.011321 |
| Cornales | Hydrangeaceae | Hydrangeaceae | *Fendlera_rupicola* | 190 | 3 | 0.015789 |
| Cornales | Hydrangeaceae | Hydrangeaceae | *Hydrangea_spp.* | 190 | 3 | 0.015789 |
| Cornales | Hydrangeaceae | Hydrangeaceae | *Philadelphus_spp.* | 190 | 3 | 0.015789 |
| Ericales | Marcgraviaceae | Marcgraviaceae | *Marcgravia_spp.* | 130 | 1 | 0.007692 |
| Ericales | Balsaminaceae | Balsaminaceae | *Impatiens_spp.* | 1001 | 1 | 0.000999 |
| Ericales | Tetrameristaceae | Tetrameristaceae | *Tetramerista_spp.* | 5 | 1 | 0.200000 |
| Ericales | Lecythidaceae | Lecythidaceae | *Barringtonia_spp.* | 340 | 2 | 0.005882 |
| Ericales | Lecythidaceae | Lecythidaceae | *Couroupita* | 340 | 2 | 0.005882 |
| Ericales | Fouquieriaceae | Fouquieriaceae | *Fouquieria_spp.* | 11 | 1 | 0.090909 |
| Ericales | Polemoniaceae | Polemoniaceae | *Cobaea_scandens* | 385 | 4 | 0.010390 |
| Ericales | Polemoniaceae | Polemoniaceae | *Gilia_spp.* | 385 | 4 | 0.010390 |
| Ericales | Polemoniaceae | Polemoniaceae | *Phlox_spp.* | 385 | 4 | 0.010390 |
| Ericales | Polemoniaceae | Polemoniaceae | *Polemonium* | 385 | 4 | 0.010390 |
| Ericales | Sapotaceae | Sapotaceae | *Manilkara_zapota* | 1100 | 1 | 0.000909 |
| Ericales | Ebenaceae | Ebenaceae | *Lissocarpa_benthamii* | 548 | 3 | 0.005474 |
| Ericales | Ebenaceae | Ebenaceae | *Diospyros_spp.* | 548 | 3 | 0.005474 |
| Ericales | Ebenaceae | Ebenaceae | *Euclea_crispa* | 548 | 3 | 0.005474 |
| Ericales | Primulaceae (Myrsinaceae) | Primulaceae | *Anagallis_spp.* | 2590 | 5 | 0.001931 |
| Ericales | Primulaceae | Primulaceae | *Maesa* | 2590 | 5 | 0.001931 |
| Ericales | Primulaceae | Primulaceae | *Clavija* | 2590 | 5 | 0.001931 |
| Ericales | Primulaceae | Primulaceae | *Androsace_spp.* | 2590 | 5 | 0.001931 |
| Ericales | Primulaceae | Primulaceae | *Primula* | 2590 | 5 | 0.001931 |
| Ericales | Pentaphylacaceae | Pentaphylacaceae plus Sladeniaceae | *Eurya_spp.* | 340 | 2 | 0.005882 |
| Ericales | Pentaphylacaceae | Pentaphylacaceae plus Sladeniaceae | *Ternstroemia_spp.* | 340 | 2 | 0.005882 |

| | | | | | | |
|---|---|---|---|---|---|---|
| Ericales | Theaceae | Theaceae plus Mitrastemonaceae | *Camellia_sinensis* | 462 | 1 | 0.002165 |
| Ericales | Symplocaceae | Symplocaceae | *Symplocos_spp.* | 320 | 1 | 0.003125 |
| Ericales | Diapensiaceae | Diapensiaceae | *Galax_spp.* | 18 | 1 | 0.055556 |
| Ericales | Styracaceae | Styracaceae | *Halesia_spp.* | 160 | 2 | 0.012500 |
| Ericales | Styracaceae | Styracaceae | *Styrax_spp.* | 160 | 2 | 0.012500 |
| Ericales | Sarraceniaceae | Sarraceniaceae | *Sarracenia_spp.* | 32 | 1 | 0.031250 |
| Ericales | Actinidiaceae | Actinidiaceae | *Actinidia_spp.* | 355 | 1 | 0.002817 |
| Ericales | Roridulaceae | Roridulaceae | *Roridula_gorgonias* | 2 | 1 | 0.500000 |
| Ericales | Clethraceae | Clethraceae | *Clethra_spp.* | 75 | 1 | 0.013333 |
| Ericales | Cyrillaceae | Cyrillaceae | *Cyrilla_racemiflora* | 2 | 1 | 0.500000 |
| Ericales | Ericaceae | Ericaceae | *Enkianthus_campanulatus* | 3995 | 4 | 0.001001 |
| Ericales | Ericaceae | Ericaceae | *Arbutus* | 3995 | 4 | 0.001001 |
| Ericales | Ericaceae | Ericaceae | *Rhododendron_spp.* | 3995 | 4 | 0.001001 |
| Ericales | Ericaceae | Ericaceae | *Vaccinium_spp.* | 3995 | 4 | 0.001001 |
| unassigned Icacinaceae | Icacinaceae | Icacinaceae | *Icacina* | 150 | 1 | 0.006667 |
| Garryales | Eucommiaceae | Eucommiaceae | *Eucommia* | 1 | 1 | 1.000000 |
| Garryales | Garryaceae (Aucubaceae) | Garryaceae | *Aucuba_japonica* | 17 | 2 | 0.117647 |
| Garryales | Garryaceae | Garryaceae | *Garrya_elliptica* | 17 | 2 | 0.117647 |
| unassigned Oncothecaceae | Oncothecaceae | Oncothecaceae | *Oncotheca* | 2 | 1 | 0.500000 |
| unassigned Vahliaceae | Vahliaceae | Vahliaceae | *Vahlia_capensis* | 8 | 1 | 0.125000 |
| Solanales | Montiniaceae | Montiniaceae | *Montinia_caryophyllacea* | 5 | 1 | 0.200000 |
| Solanales | Hydroleaceae | Hydroleaceae | *Hydrolea_ovata* | 12 | 1 | 0.083333 |
| Solanales | Sphenocleaceae | Sphenocleaceae | *Sphenoclea* | 2 | 1 | 0.500000 |
| Solanales | Convolvulaceae | Convolvulaceae | *Cuscuta* | 1625 | 2 | 0.001231 |
| Solanales | Convolvulaceae | Convolvulaceae | *Ipomoea_spp.* | 1625 | 2 | 0.001231 |
| Solanales | Solanaceae | Solanaceae | *Petunia_spp.* | 2460 | 5 | 0.002033 |
| Solanales | Solanaceae | Solanaceae | *Atropa* | 2460 | 5 | 0.002033 |
| Solanales | Solanaceae | Solanaceae | *Solanum* | 2460 | 5 | 0.002033 |
| Solanales | Solanaceae | Solanaceae | *Nicotiana_spp.* | 2460 | 5 | 0.002033 |
| Solanales | Solanaceae | Solanaceae | *Nolana_spp.* | 2460 | 5 | 0.002033 |
| Gentianales | Rubiaceae | Rubiaceae | *Luculia* | 13150 | 4 | 0.000304 |
| Gentianales | Rubiaceae | Rubiaceae | *Coffea_arabica* | 13150 | 4 | 0.000304 |
| Gentianales | Rubiaceae | Rubiaceae | *Galium* | 13150 | 4 | 0.000304 |
| Gentianales | Rubiaceae | Rubiaceae | *Mitchella_repens* | 13150 | 4 | 0.000304 |
| Gentianales | Loganiaceae | Loganiaceae | *Spigelia_spp.* | 420 | 2 | 0.004762 |
| Gentianales | Loganiaceae | Loganiaceae | *Strychnos* | 420 | 2 | 0.004762 |
| Gentianales | Gelsemiaceae | Gelsemiaceae | *Gelsemium* | 11 | 1 | 0.090909 |
| Gentianales | Apocynaceae | Apocynaceae | *Nerium_oleander* | 4555 | 1 | 0.000220 |
| Gentianales | Gentianaceae | Gentianaceae | *Exacum_affine* | 1675 | 2 | 0.001194 |
| Gentianales | Gentianaceae | Gentianaceae | *Gentiana_spp.* | 1675 | 2 | 0.001194 |
| Boraginales | Boraginaceae | Boraginaceae | *Borago_officinalis* | 2755 | 3 | 0.001089 |
| Boraginales | Boraginaceae | Boraginaceae | *Ehretia* | 2755 | 3 | 0.001089 |
| Boraginales | Boraginaceae | Boraginaceae | *Hydrophyllum* | 2755 | 3 | 0.001089 |

| | | | | | | |
|---|---|---|---|---|---|---|
| Lamiales | Plocospermataceae | Plocospermtaceae | *Plocosperma_buxifolium* | 1 | 1 | 1.000000 |
| Lamiales | Oleaceae | Oleaceae plus Carlemanniaceae | *Olea_europaea* | 620 | 3 | 0.004839 |
| Lamiales | Oleaceae | Oleaceae plus Carlemanniaceae | *Jasminum* | 620 | 3 | 0.004839 |
| Lamiales | Oleaceae | Oleaceae plus Carlemanniaceae | *Syringa* | 620 | 3 | 0.004839 |
| Lamiales | Tetrachondraceae | Tetrachondraceae | *Polypremum_procumbens* | 3 | 1 | 0.333333 |
| Lamiales | unplaced Peltanthera | Peltanthera | *Peltanthera* | 1 | 1 | 1.000000 |
| Lamiales | Calceolariaceae | Calceolariaceae | *Calceolaria* | 260 | 1 | 0.003846 |
| Lamiales | Gesneriaceae | Gesneriaceae | *Rhynchoglossum_spp.* | 3312 | 2 | 0.000604 |
| Lamiales | Gesneriaceae | Gesneriaceae | *Titanotrichum_oldhamii* | 3312 | 2 | 0.000604 |
| Lamiales | Stilbaceae | Stilbaceae | *Halleria* | 39 | 1 | 0.025641 |
| Lamiales | Plantaginaceae | Plantaginaceae | *Antirrhinum* | 1900 | 3 | 0.001579 |
| Lamiales | Plantaginaceae | Plantaginaceae | *Plantago_spp.* | 1900 | 3 | 0.001579 |
| Lamiales | Plantaginaceae | Plantaginaceae | *Veronica_spp.* | 1900 | 3 | 0.001579 |
| Lamiales | Scrophulariaceae | Scrophulariaceae | *Myoporum_mauritianum* | 1800 | 3 | 0.001667 |
| Lamiales | Scrophulariaceae | Scrophulariaceae | *Scrophularia_spp.* | 1800 | 3 | 0.001667 |
| Lamiales | Scrophulariaceae | Scrophulariaceae | *Verbascum_thapsus* | 1800 | 3 | 0.001667 |
| Lamiales | Bignoniaceae | Bignoniaceae | *Campsis_radicans* | 800 | 2 | 0.002500 |
| Lamiales | Bignoniaceae | Bignoniaceae | *Catalpa* | 800 | 2 | 0.002500 |
| Lamiales | Martyniaceae | Martyniaceae plus Schlegeliaceae | *Martynia* | 44 | 1 | 0.022727 |
| Lamiales | Pedaliaceae | Pedaliaceae | *Sesamum* | 70 | 1 | 0.014286 |
| Lamiales | Byblidaceae | Byblidaceae plus Linderniaceae | *Byblis_spp.* | 201 | 1 | 0.004975 |
| Lamiales | Lentibulariaceae | Lentibulariaceae | *Pinguicula* | 330 | 2 | 0.006061 |
| Lamiales | Lentibulariaceae | Lentibulariaceae | *Utricularia_spp.* | 330 | 2 | 0.006061 |
| Lamiales | Acanthaceae | Acanthaceae | *Thunbergia_spp.* | 4000 | 4 | 0.001000 |
| Lamiales | Acanthaceae | Acanthaceae | *Acanthus* | 4000 | 4 | 0.001000 |
| Lamiales | Acanthaceae | Acanthaceae | *Barleria_prionitis* | 4000 | 4 | 0.001000 |
| Lamiales | Acanthaceae | Acanthaceae | *Justicia_americana* | 4000 | 4 | 0.001000 |
| Lamiales | Thomandersiaceae | Thomandersiaceae | *Thomandersia* | 6 | 1 | 0.166667 |
| Lamiales | Verbenaceae | Verbenaceae | *Verbena_spp.* | 918 | 1 | 0.001089 |
| Lamiales | Lamiaceae | Lamiaceae | *Callicarpa_spp.* | 7173 | 2 | 0.000279 |
| Lamiales | Lamiaceae | Lamiaceae | *Lamium_spp.* | 7173 | 2 | 0.000279 |
| Lamiales | Paulowniaceae | Paulowniaceae | *Paulownia_tomentosa* | 8 | 1 | 0.125000 |
| Lamiales | Orobanchaceae | Orobanchaceae | *Pedicularis_spp.* | 2060 | 1 | 0.000485 |
| Lamiales | Mazaceae | Mazaceae | *Mazus_spp.* | 33 | 1 | 0.030303 |
| Lamiales | Phrymaceae | Phrymaceae | *Phryma* | 188 | 1 | 0.005319 |
| Aquifoliales | Aquifoliaceae | Aquifoliaceae | *Ilex_spp.* | 405 | 1 | 0.002469 |

| | | | | | | |
|---|---|---|---|---|---|---|
| Aquifoliales | Helwingiaceae | Helwingiaceae | *Helwingia_spp.* | 3 | 1 | 0.333333 |
| Aquifoliales | Phyllonomaceae | Phyllonomaceae | *Phyllonoma_spp.* | 4 | 1 | 0.250000 |
| Aquifoliales | Stemonuraceae | Stemonuraceae | *Gomphandra* | 95 | 2 | 0.021053 |
| Aquifoliales | Stemonuraceae | Stemonuraceae | *Irvingbaileya_spp.* | 95 | 2 | 0.021053 |
| Aquifoliales | Cardiopteridaceae | Cardiopteridaceae | *Citronella* | 43 | 3 | 0.069767 |
| Aquifoliales | Cardiopteridaceae | Cardiopteridaceae | *Cardiopteris_spp.* | 43 | 3 | 0.069767 |
| Aquifoliales | Cardiopteridaceae | Cardiopteridaceae | *Gonocaryum_litorale* | 43 | 3 | 0.069767 |
| Escalloniales | Escalloniaceae | Escalloniaceae | *Polyosma_spp.* | 130 | 6 | 0.046154 |
| Escalloniales | Escalloniaceae | Escalloniaceae | *Anopterus* | 130 | 6 | 0.046154 |
| Escalloniales | Escalloniaceae | Escalloniaceae | *Eremosyne_pectinata* | 130 | 6 | 0.046154 |
| Escalloniales | Escalloniaceae | Escalloniaceae | *Escallonia_spp.* | 130 | 6 | 0.046154 |
| Escalloniales | Escalloniaceae | Escalloniaceae | *Forgesia* | 130 | 6 | 0.046154 |
| Escalloniales | Escalloniaceae | Escalloniaceae | *Valdivia* | 130 | 6 | 0.046154 |
| Asterales | Pentaphragmataceae | Pentaphragmataceae | *Pentaphragma* | 30 | 1 | 0.033333 |
| Asterales | Rousseaceae | Rousseaceae | *Roussea_simplex* | 13 | 4 | 0.307692 |
| Asterales | Rousseaceae | Rousseaceae | *Carpodetus* | 13 | 4 | 0.307692 |
| Asterales | Rousseaceae | Rousseaceae | *Abrophyllum* | 13 | 4 | 0.307692 |
| Asterales | Rousseaceae | Rousseaceae | *Cuttsia* | 13 | 4 | 0.307692 |
| Asterales | Campanulaceae | Campanulaceae | *Cyphia* | 2380 | 6 | 0.002521 |
| Asterales | Campanulaceae | Campanulaceae | *Campanula_spp.* | 2380 | 6 | 0.002521 |
| Asterales | Campanulaceae | Campanulaceae | *Trachelium* | 2380 | 6 | 0.002521 |
| Asterales | Campanulaceae | Campanulaceae | *Pseudonemacladus* | 2380 | 6 | 0.002521 |
| Asterales | Campanulaceae | Campanulaceae | *Dialypetalum* | 2380 | 6 | 0.002521 |
| Asterales | Campanulaceae | Campanulaceae | *Lobelia_spp.* | 2380 | 6 | 0.002521 |
| Asterales | Phellinaceae | Phellinaceae | *Phelline_spp.* | 12 | 1 | 0.083333 |
| Asterales | Argophyllaceae | Argophyllaceae | *Argophyllum* | 21 | 2 | 0.095238 |
| Asterales | Argophyllaceae | Argophyllaceae | *Corokia_cotoneaster* | 21 | 2 | 0.095238 |
| Asterales | Alseuosmiaceae | Alseusomiaceae | *Platyspermation* | 10 | 4 | 0.400000 |
| Asterales | Alseuosmiaceae | Alseusomiaceae | *Crispiloba* | 10 | 4 | 0.400000 |
| Asterales | Alseuosmiaceae | Alseusomiaceae | *Alseuosmia* | 10 | 4 | 0.400000 |
| Asterales | Alseuosmiaceae | Alseusomiaceae | *Wittsteinia* | 10 | 4 | 0.400000 |
| Asterales | Stylidaceae | Stylidaceae | *Donatia_spp.* | 245 | 3 | 0.012245 |
| Asterales | Stylidaceae | Stylidaceae | *Forstera* | 245 | 3 | 0.012245 |
| Asterales | Stylidaceae | Stylidaceae | *Stylidium* | 245 | 3 | 0.012245 |
| Asterales | Menyanthaceae | Menyanthaceae | *Fauria* | 58 | 4 | 0.068966 |
| Asterales | Menyanthaceae | Menyanthaceae | *Menyanthes_trifoliata* | 58 | 4 | 0.068966 |
| Asterales | Menyanthaceae | Menyanthaceae | *Nymphoides* | 58 | 4 | 0.068966 |
| Asterales | Menyanthaceae | Menyanthaceae | *Villarsia* | 58 | 4 | 0.068966 |
| Asterales | Goodeniaceae | Goodeniaceae | *Dampiera* | 430 | 3 | 0.006977 |
| Asterales | Goodeniaceae | Goodeniaceae | *Goodenia* | 430 | 3 | 0.006977 |
| Asterales | Goodeniaceae | Goodeniaceae | *Scaevola_spp.* | 430 | 3 | 0.006977 |
| Asterales | Calyceraceae | Calyceraceae | *Boopis_spp.* | 60 | 3 | 0.050000 |
| Asterales | Calyceraceae | Calyceraceae | *Acicarpha* | 60 | 3 | 0.050000 |
| Asterales | Calyceraceae | Calyceraceae | *Moschopsis* | 60 | 3 | 0.050000 |
| Asterales | Asteraceae | Asteraceae | *Barnadesia* | 23600 | 9 | 0.000381 |
| Asterales | Asteraceae | Asteraceae | *Gerbera* | 23600 | 9 | 0.000381 |
| Asterales | Asteraceae | Asteraceae | *Echinops* | 23600 | 9 | 0.000381 |
| Asterales | Asteraceae | Asteraceae | *Tragopogon_dubius* | 23600 | 9 | 0.000381 |
| Asterales | Asteraceae | Asteraceae | *Cichorium* | 23600 | 9 | 0.000381 |

| | | | | | | |
|---|---|---|---|---|---|---|
| Asterales | Asteraceae | Asteraceae | *Lactuca* | 23600 | 9 | 0.000381 |
| Asterales | Asteraceae | Asteraceae | *Tagetes* | 23600 | 9 | 0.000381 |
| Asterales | Asteraceae | Asteraceae | *Guizotia* | 23600 | 9 | 0.000381 |
| Asterales | Asteraceae | Asteraceae | *Helianthus_annuus* | 23600 | 9 | 0.000381 |
| Bruniales | Bruniaceae | Bruniaceae | *Berzelia_lanuginosa* | 81 | 2 | 0.024691 |
| Bruniales | Bruniaceae | Bruniaceae | *Brunia* | 81 | 2 | 0.024691 |
| Bruniales | Columelliaceae | Columelliaceae | *Columellia* | 5 | 2 | 0.400000 |
| Bruniales | Columelliaceae | Columelliaceae | *Desfontainia_spinosa* | 5 | 2 | 0.400000 |
| Paracryphiales | Paracryphiaceae | Paracryphiaceae | *Quintinia* | 36 | 3 | 0.083333 |
| Paracryphiales | Paracryphiaceae | Paracryphiaceae | *Paracryphia_alticola* | 36 | 3 | 0.083333 |
| Paracryphiales | Paracryphiaceae | Paracryphiaceae | *Sphenostemon* | 36 | 3 | 0.083333 |
| Dipsacales | Adoxaceae | Adoxaceae | *Viburnum_spp.* | 200 | 5 | 0.025000 |
| Dipsacales | Adoxaceae | Adoxaceae | *Sambucus* | 200 | 5 | 0.025000 |
| Dipsacales | Adoxaceae | Adoxaceae | *Sinadoxa* | 200 | 5 | 0.025000 |
| Dipsacales | Adoxaceae | Adoxaceae | *Adoxa_moschatellina* | 200 | 5 | 0.025000 |
| Dipsacales | Adoxaceae | Adoxaceae | *Tetradoxa* | 200 | 5 | 0.025000 |
| Dipsacales | Caprifoliaceae (Diervillaceae) | Caprifoliaceae | *Diervilla* | 890 | 22 | 0.024719 |
| Dipsacales | Caprifoliaceae (Diervillaceae) | Caprifoliaceae | *Weigela* | 890 | 22 | 0.024719 |
| Dipsacales | Caprifoliaceae | Caprifoliaceae | *Heptacodium* | 890 | 22 | 0.024719 |
| Dipsacales | Caprifoliaceae | Caprifoliaceae | *Lonicera* | 890 | 22 | 0.024719 |
| Dipsacales | Caprifoliaceae | Caprifoliaceae | *Symphoricarpos* | 890 | 22 | 0.024719 |
| Dipsacales | Caprifoliaceae | Caprifoliaceae | *Leycesteria* | 890 | 22 | 0.024719 |
| Dipsacales | Caprifoliaceae | Caprifoliaceae | *Triosteum* | 890 | 22 | 0.024719 |
| Dipsacales | Caprifoliaceae (Linnaeaceae) | Caprifoliaceae | *Linnaea* | 890 | 22 | 0.024719 |
| Dipsacales | Caprifoliaceae (Linnaeaceae) | Caprifoliaceae | *Abelia_spp.* | 890 | 22 | 0.024719 |
| Dipsacales | Caprifoliaceae (Linnaeaceae) | Caprifoliaceae | *Dipelta* | 890 | 22 | 0.024719 |
| Dipsacales | Caprifoliaceae (Linnaeaceae) | Caprifoliaceae | *Kolkwitzia* | 890 | 22 | 0.024719 |
| Dipsacales | Caprifoliaceae (Morinaceae) | Caprifoliaceae | *Morina* | 890 | 22 | 0.024719 |
| Dipsacales | Caprifoliaceae (Morinaceae) | Caprifoliaceae | *Zabelia* | 890 | 22 | 0.024719 |
| Dipsacales | Caprifoliaceae (Dipsacaceae) | Caprifoliaceae | *Triplostegia* | 890 | 22 | 0.024719 |
| Dipsacales | Caprifoliaceae (Dipsacaceae) | Caprifoliaceae | *Scabiosa_spp.* | 890 | 22 | 0.024719 |
| Dipsacales | Caprifoliaceae (Dipsacaceae) | Caprifoliaceae | *Dipsacus_spp.* | 890 | 22 | 0.024719 |
| Dipsacales | Caprifoliaceae (Dipsacaceae) | Caprifoliaceae | *Pterocephalodes* | 890 | 22 | 0.024719 |
| Dipsacales | Caprifoliaceae (Valerianaceae) | Caprifoliaceae | *Patrinia* | 890 | 22 | 0.024719 |
| Dipsacales | Caprifoliaceae (Valerianaceae) | Caprifoliaceae | *Nardostachys* | 890 | 22 | 0.024719 |

| | | | | | | |
|---|---|---|---|---|---|---|
| Dipsacales | Caprifoliaceae (Valerianaceae) | Caprifoliaceae | *Valerianella* | 890 | 22 | 0.024719 |
| Dipsacales | Caprifoliaceae (Valerianaceae) | Caprifoliaceae | *Centranthus* | 890 | 22 | 0.024719 |
| Dipsacales | Caprifoliaceae (Valerianaceae) | Caprifoliaceae | *Valeriana_spp.* | 890 | 22 | 0.024719 |
| Apiales | Pennantiaceae | Pennantiaceae | *Pennantia_corymbosa* | 4 | 1 | 0.250000 |
| Apiales | Torricelliaceae | Torricelliaceae | *Aralidium* | 10 | 3 | 0.300000 |
| Apiales | Torricelliaceae | Torricelliaceae | *Melanophylla_spp.* | 10 | 3 | 0.300000 |
| Apiales | Torricelliaceae | Torricelliaceae | *Torricellia* | 10 | 3 | 0.300000 |
| Apiales | Griseliniaceae | Griseliniaceae | *Griselinia_spp.* | 6 | 1 | 0.166667 |
| Apiales | Pittosporaceae | Pittosporaceae | *Pittosporum* | 200 | 2 | 0.010000 |
| Apiales | Pittosporaceae | Pittosporaceae | *Sollya_heterophylla* | 200 | 2 | 0.010000 |
| Apiales | Araliaceae | Arailaceae | *Hydrocotyle* | 1450 | 10 | 0.006897 |
| Apiales | Araliaceae | Arailaceae | *Aralia* | 1450 | 10 | 0.006897 |
| Apiales | Araliaceae | Arailaceae | *Panax_spp.* | 1450 | 10 | 0.006897 |
| Apiales | Araliaceae | Arailaceae | *Cussonia* | 1450 | 10 | 0.006897 |
| Apiales | Araliaceae | Arailaceae | *Pseudopanax* | 1450 | 10 | 0.006897 |
| Apiales | Araliaceae | Arailaceae | *Tetraplasandra* | 1450 | 10 | 0.006897 |
| Apiales | Araliaceae | Arailaceae | *Polyscias* | 1450 | 10 | 0.006897 |
| Apiales | Araliaceae | Arailaceae | *Hedera_helix* | 1450 | 10 | 0.006897 |
| Apiales | Araliaceae | Arailaceae | *Schefflera* | 1450 | 10 | 0.006897 |
| Apiales | Araliaceae | Arailaceae | *Tetrapanax* | 1450 | 10 | 0.006897 |
| Apiales | Myodocarpaceae | Myodocarpaceae | *Delarbrea_michieana* | 19 | 2 | 0.105263 |
| Apiales | Myodocarpaceae | Myodocarpaceae | *Myodocarpus* | 19 | 2 | 0.105263 |
| Apiales | Apiaceae | Apiaceae | *Mackinlaya* | 3780 | 11 | 0.002910 |
| Apiales | Apiaceae | Apiaceae | *Platysace* | 3780 | 11 | 0.002910 |
| Apiales | Apiaceae | Apiaceae | *Azorella* | 3780 | 11 | 0.002910 |
| Apiales | Apiaceae | Apiaceae | *Arctopus* | 3780 | 11 | 0.002910 |
| Apiales | Apiaceae | Apiaceae | *Sanicula* | 3780 | 11 | 0.002910 |
| Apiales | Apiaceae | Apiaceae | *Heteromorpha* | 3780 | 11 | 0.002910 |
| Apiales | Apiaceae | Apiaceae | *Daucus* | 3780 | 11 | 0.002910 |
| Apiales | Apiaceae | Apiaceae | *Anethum* | 3780 | 11 | 0.002910 |
| Apiales | Apiaceae | Apiaceae | *Apium_graveolens* | 3780 | 11 | 0.002910 |
| Apiales | Apiaceae | Apiaceae | *Angelica* | 3780 | 11 | 0.002910 |
| Apiales | Apiaceae | Apiaceae | *Coriandrum* | 3780 | 11 | 0.002910 |
| Saxifragales | Peridiscaceae | Peridiscaceae | *Peridiscus_lucidus* | 11 | 2 | 0.181818 |
| Saxifragales | Peridiscaceae | Peridiscaceae | *Soyauxia_spp.* | 11 | 2 | 0.181818 |
| Saxifragales | Paeoniaceae | Paeoniaceae | *Paeonia_californica* | 33 | 1 | 0.030303 |
| Saxifragales | Altingiaceae | Altingiaceae | *Altingia_spp.* | 13 | 2 | 0.153846 |
| Saxifragales | Altingiaceae | Altingiaceae | *Liquidambar_styraciflua* | 13 | 2 | 0.153846 |
| Saxifragales | Cercidiphyllaceae | Cercidiphyllaceae | *Cercidiphyllum_japonicum* | 2 | 1 | 0.500000 |
| Saxifragales | Daphniphyllaceae | Daphiphyllaceae | *Daphniphyllum_spp.* | 10 | 1 | 0.100000 |
| Saxifragales | Hamamelidaceae | Hamamelidaceae | *Exbucklandia_populnea* | 82 | 5 | 0.060976 |
| Saxifragales | Hamamelidaceae | Hamamelidaceae | *Rhodoleia_spp.* | 82 | 5 | 0.060976 |

| | | | | | | |
|---|---|---|---|---|---|---|
| Saxifragales | Hamamelidaceae | Hamamelidaceae | *Disanthus_cercidifolius* | 82 | 5 | 0.060976 |
| Saxifragales | Hamamelidaceae | Hamamelidaceae | *Corylopsis_spp.* | 82 | 5 | 0.060976 |
| Saxifragales | Hamamelidaceae | Hamamelidaceae | *Hamamelis_spp.* | 82 | 5 | 0.060976 |
| Saxifragales | Pterostemonaceae | Pterostemonaceae | *Pterostemon_rotundifolius* | 3 | 1 | 0.333333 |
| Saxifragales | Iteaceae | Iteaceae | *Choristylis* | 18 | 2 | 0.111111 |
| Saxifragales | Iteaceae | Iteaceae | *Itea* | 18 | 2 | 0.111111 |
| Saxifragales | Grossulariaceae | Grossulariaceae | *Ribes_aureum* | 150 | 1 | 0.006667 |
| Saxifragales | Saxifragaceae | Saxifragaceae | *Saxifraga* | 540 | 3 | 0.005556 |
| Saxifragales | Saxifragaceae | Saxifragaceae | *Heuchera_spp.* | 540 | 3 | 0.005556 |
| Saxifragales | Saxifragaceae | Saxifragaceae | *Sullivantia_oregana* | 540 | 3 | 0.005556 |
| Saxifragales | Crassulaceae | Crassulaceae | *Crassula_marnierana* | 1400 | 4 | 0.002857 |
| Saxifragales | Crassulaceae | Crassulaceae | *Kalanchoe* | 1400 | 4 | 0.002857 |
| Saxifragales | Crassulaceae | Crassulaceae | *Dudleya_viscida* | 1400 | 4 | 0.002857 |
| Saxifragales | Crassulaceae | Crassulaceae | *Sedum_spp.* | 1400 | 4 | 0.002857 |
| Saxifragales | Aphanopetalaceae | Aphanopetalaceae | *Aphanopetalum_resinosum* | 2 | 1 | 0.500000 |
| Saxifragales | Tetracarpaeaceae | Tetracarpaea | *Tetracarpaea_tasmanica* | 1 | 1 | 1.000000 |
| Saxifragales | Penthoraceae | Penthoraceae | *Penthorum_sedoides* | 2 | 1 | 0.500000 |
| Saxifragales | Haloragaceae | Haloragaceae | *Haloragis_spp.* | 145 | 2 | 0.013793 |
| Saxifragales | Haloragaceae | Haloragaceae | *Myriophyllum* | 145 | 2 | 0.013793 |
| Vitales | Vitaceae | Vitaceae | *Leea_guineensis* | 850 | 2 | 0.002353 |
| Vitales | Vitaceae | Vitaceae | *Vitis_spp.* | 850 | 2 | 0.002353 |
| Geraniales | Geraniaceae | Geraniaceae | *Geranium_spp.* | 805 | 2 | 0.002484 |
| Geraniales | Geraniaceae | Geraniaceae | *Pelargonium_cotyledonis* | 805 | 2 | 0.002484 |
| Geraniales | Vivianiaceae | Vivianiaceae | *Viviania_marifolia* | 18 | 1 | 0.055556 |
| Geraniales | Greyiaceae | Greyiaceae plus Francoaceae | *Greyia_radlkoferi* | 5 | 1 | 0.200000 |
| Geraniales | Melianthaceae | Melianthaceae | *Melianthus* | 8 | 1 | 0.125000 |
| Myrtales | Combretaceae | Combretaceae | *Terminalia_spp.* | 500 | 1 | 0.002000 |
| Myrtales | Lythraceae | Lythraceae | *Lythrum_spp.* | 620 | 1 | 0.001613 |
| Myrtales | Onagraceae | Onagraceae | *Fuchsia_procumbens* | 656 | 3 | 0.004573 |
| Myrtales | Onagraceae | Onagraceae | *Clarkia_xantiana* | 656 | 3 | 0.004573 |
| Myrtales | Onagraceae | Onagraceae | *Oenothera* | 656 | 3 | 0.004573 |
| Myrtales | Melastomataceae | Melastomataceae | *Clidemia_spp.* | 5005 | 2 | 0.000400 |
| Myrtales | Melastomataceae | Melastomataceae | *Mouriri_cyphocarpa* | 5005 | 2 | 0.000400 |
| Myrtales | Crypteroniaceae | Crypteroniaceae | *Crypteronia_paniculata* | 10 | 1 | 0.100000 |
| Myrtales | Penaeaceae | Penaeaceae plus Alzateaceae | *Olinia* | 31 | 1 | 0.032258 |
| Myrtales | Vochysiaceae | Vochysiaceae | *Qualea* | 190 | 2 | 0.010526 |
| Myrtales | Vochysiaceae | Vochysiaceae | *Vochysia_spp.* | 190 | 2 | 0.010526 |
| Myrtales | Myrtaceae | Myrtaceae | *Heteropyxis_natalensis* | 4620 | 4 | 0.000866 |
| Myrtales | Myrtaceae | Myrtaceae | *Metrosideros_spp.* | 4620 | 4 | 0.000866 |
| Myrtales | Myrtaceae | Myrtaceae | *Eucalyptus* | 4620 | 4 | 0.000866 |

| | | | | | | |
|---|---|---|---|---|---|---|
| Myrtales | Myrtaceae | Myrtaceae | *Myrtus* | 4620 | 4 | 0.000866 |
| Crossosomatales | Aphloiaceae | Aphloiaceae | *Aphloia_theiformis* | 1 | 1 | 1.000000 |
| Crossosomatales | Geissolomataceae | Geissolomataceae | *Geissoloma_marginatum* | 1 | 1 | 1.000000 |
| Crossosomatales | Ixerbaceae | Ixerbaceae | *Ixerba* | 1 | 1 | 1.000000 |
| Crossosomatales | Strasburgeriaceae | Strasburgeriaceae | *Strasburgeria_spp.* | 2 | 1 | 0.500000 |
| Crossosomatales | Staphyleaceae | Staphyleaceae | *Staphylea_spp.* | 45 | 1 | 0.022222 |
| Crossosomatales | Guamatelaceae | Guamatelaceae | *Guamatela_tuerckheimii* | 1 | 1 | 1.000000 |
| Crossosomatales | Crossosomataceae | Crossosomataceae | *Crossosoma_spp.* | 12 | 1 | 0.083333 |
| Crossosomatales | Stachyuraceae | Stachyuraceae | *Stachyurus* | 5 | 1 | 0.200000 |
| Picramniales | Picramniaceae | Picramniaceae | *Picramnia_spp.* | 49 | 1 | 0.020408 |
| Sapindales | Biebersteiniaceae | Biebersteiniaceae | *Biebersteinia_orphanidis* | 5 | 1 | 0.200000 |
| Sapindales | Nitrariaceae | Nitrariaceae | *Nitraria* | 16 | 1 | 0.062500 |
| Sapindales | Burseraceae | Burseraceae plus Kirkiaceae | *Bursera_inaguensis* | 763 | 1 | 0.001311 |
| Sapindales | Anacardiaceae | Anacardiaceae | *Rhus_spp.* | 873 | 2 | 0.002291 |
| Sapindales | Anacardiaceae | Anacardiaceae | *Schinus* | 873 | 2 | 0.002291 |
| Sapindales | Sapindaceae | Sapindaceae | *Aesculus_spp.* | 1630 | 2 | 0.001227 |
| Sapindales | Sapindaceae | Sapindaceae | *Cupaniopsis_anacardioides* | 1630 | 2 | 0.001227 |
| Sapindales | Simaroubaceae | Simaroubaceae | *Ailanthus_altissima* | 110 | 1 | 0.009091 |
| Sapindales | Rutaceae | Rutaceae | *Citrus_spp.* | 2070 | 1 | 0.000483 |
| Sapindales | Meliaceae | Meliaceae | *Swietenia_macrophylla* | 615 | 2 | 0.003252 |
| Sapindales | Meliaceae | Meliaceae | *Trichilia_emetica* | 615 | 2 | 0.003252 |
| Huerteales | Gerrardiniaceae | Gerrardiniaceae plus Petenaceae | *Gerrardina_foliosa* | 3 | 1 | 0.333333 |
| Huerteales | Tapisciaceae | Tapisiaceae | *Tapiscia_sinensis* | 5 | 1 | 0.200000 |
| Huerteales | Dipentodontaceae | Dipentodontaceae | *Dipentodon* | 16 | 2 | 0.125000 |
| Huerteales | Dipentodontaceae | Dipentodontaceae | *Perrottetia_ovata* | 16 | 2 | 0.125000 |
| Malvales | Neuradaceae | Neuradaceae | *Neurada_procumbens* | 10 | 1 | 0.100000 |
| Malvales | Bixaceae | Bixaceae plus Sphaerosepalaceae | *Bixa* | 39 | 1 | 0.025641 |
| Malvales | Dipterocarpaceae | Dipterocarpaceae plus Sarcolaenaceae | *Anisoptera_marginata* | 740 | 1 | 0.001351 |
| Malvales | Cistaceae | Cistaceae plus Cytinaceae plus Muntingiaceae | *Helianthemum* | 188 | 1 | 0.005319 |
| Malvales | Thymelaeaceae | Thymelaeaceae | *Thymelaea_hirsuta* | 891 | 1 | 0.001122 |
| Malvales | Malvaceae | Malvaceae | *Bombax_spp.* | 4225 | 5 | 0.001183 |
| Malvales | Malvaceae | Malvaceae | *Ochroma_pyramidale* | 4225 | 5 | 0.001183 |

| | | | | | | |
|---|---|---|---|---|---|---|
| Malvales | Malvaceae | Malvaceae | *Durio_zibethinus* | 4225 | 5 | 0.001183 |
| Malvales | Malvaceae | Malvaceae | *Gossypium* | 4225 | 5 | 0.001183 |
| Malvales | Malvaceae | Malvaceae | *Sterculia_spp.* | 4225 | 5 | 0.001183 |
| Brassicales | Caricaceae | Caricaceae plus Moringaceae | *Carica_papaya* | 46 | 1 | 0.021739 |
| Brassicales | Tropaeolaceae | Tropaeolaceae plus Akaniaceae | *Tropaeolum_spp.* | 107 | 1 | 0.009346 |
| Brassicales | Limnanthaceae | Limnanthaceae plus Setchellanthaceae | *Floerkea_proserpinacoides* | 9 | 1 | 0.111111 |
| Brassicales | Bataceae | Bataceae plus Salvadoraceae | *Batis_maritima* | 13 | 1 | 0.076923 |
| Brassicales | Koeberliniaceae | Koeberliniaceae | *Koeberlinia_spinosa* | 2 | 1 | 0.500000 |
| Brassicales | Gyrostemonaceae | Gyrostemonaceae | *Gyrostemon_spp.* | 18 | 1 | 0.055556 |
| Brassicales | Resedaceae | Resedaceae | *Reseda* | 75 | 1 | 0.013333 |
| Brassicales | Tovariaceae | Tovariaceae | *Tovaria_pendula* | 2 | 1 | 0.500000 |
| Brassicales | Pentadiplandraceae | Pentadiplandraceae plus Emblingiaceae | *Pentadiplandra_brazzeana* | 2 | 1 | 0.500000 |
| Brassicales | Capparaceae | Capparaceae | *Capparis_spp.* | 480 | 1 | 0.002083 |
| Brassicales | Brassicaceae | Brassicaceae plus Cleomaceae | *Arabidopsis_thaliana* | 4010 | 3 | 0.000748 |
| Brassicales | Brassicaceae | Brassicaceae plus Cleomaceae | *Brassica_spp.* | 4010 | 3 | 0.000748 |
| Brassicales | Brassicaceae | Brassicaceae plus Cleomaceae | *Raphanus_sativus* | 4010 | 3 | 0.000748 |
| Zygophyllales | Krameriaceae | Krameriaceae | *Krameria_ixine* | 18 | 1 | 0.055556 |
| Zygophyllales | Zygophyllaceae | Zygophyllaceae | *Guaiacum* | 285 | 2 | 0.007018 |
| Zygophyllales | Zygophyllaceae | Zygophyllaceae | *Larrea_tridentata* | 285 | 2 | 0.007018 |
| Fabales | Quillajaceae | Quillajaceae | *Quillaja_saponaria* | 3 | 1 | 0.333333 |
| Fabales | Polygalaceae | Polygalaceae | *Polygala_spp.* | 965 | 1 | 0.001036 |
| Fabales | Surianaceae | Surianaceae | *Stylobasium_spathulatum* | 8 | 1 | 0.125000 |
| Fabales | Fabaceae | Fabaceae | *Schotia_spp.* | 19500 | 15 | 0.000769 |
| Fabales | Fabaceae | Fabaceae | *Bauhinia_spp.* | 19500 | 15 | 0.000769 |
| Fabales | Fabaceae | Fabaceae | *Cercis_canadensis* | 19500 | 15 | 0.000769 |
| Fabales | Fabaceae | Fabaceae | *Ceratonia_siliqua* | 19500 | 15 | 0.000769 |
| Fabales | Fabaceae | Fabaceae | *Acacia_spp.* | 19500 | 15 | 0.000769 |
| Fabales | Fabaceae | Fabaceae | *Albizia_julibrissin* | 19500 | 15 | 0.000769 |
| Fabales | Fabaceae | Fabaceae | *Mimosa_spp.* | 19500 | 15 | 0.000769 |
| Fabales | Fabaceae | Fabaceae | *Indigofera_spp.* | 19500 | 15 | 0.000769 |
| Fabales | Fabaceae | Fabaceae | *Erythrina_spp.* | 19500 | 15 | 0.000769 |
| Fabales | Fabaceae | Fabaceae | *Glycine_max* | 19500 | 15 | 0.000769 |
| Fabales | Fabaceae | Fabaceae | *Lotus* | 19500 | 15 | 0.000769 |
| Fabales | Fabaceae | Fabaceae | *Astragalus_membranaceus* | 19500 | 15 | 0.000769 |
| Fabales | Fabaceae | Fabaceae | *Cicer* | 19500 | 15 | 0.000769 |
| Fabales | Fabaceae | Fabaceae | *Medicago_spp.* | 19500 | 15 | 0.000769 |
| Fabales | Fabaceae | Fabaceae | *Pisum* | 19500 | 15 | 0.000769 |
| Rosales | Rosaceae | Rosaceae | *Spiraea_spp.* | 2520 | 3 | 0.001190 |
| Rosales | Rosaceae | Rosaceae | *Photinia_spp.* | 2520 | 3 | 0.001190 |

| | | | | | | |
|---|---|---|---|---|---|---|
| Rosales | Rosaceae | Rosaceae | *Prunus_persica* | 2520 | 3 | 0.001190 |
| Rosales | Dirachmaceae | Dirachmaceae | *Dirachma_socotrana* | 2 | 1 | 0.500000 |
| Rosales | Elaeagnaceae | Elaeagnaceae | *Elaeagnus_spp.* | 45 | 1 | 0.022222 |
| Rosales | Barbeyaceae | Barbeyaceae | *Barbeya_oleoides* | 1 | 1 | 1.000000 |
| Rosales | Rhamnaceae | Rhamnaceae | *Ceanothus_spp.* | 925 | 2 | 0.002162 |
| Rosales | Rhamnaceae | Rhamnaceae | *Rhamnus* | 925 | 2 | 0.002162 |
| Rosales | Ulmaceae | Ulmaceae | *Zelkova_spp.* | 35 | 1 | 0.028571 |
| Rosales | Cannabaceae | Cannabaceae | *Celtis_spp.* | 170 | 3 | 0.017647 |
| Rosales | Cannabaceae | Cannabaceae | *Cannabis* | 170 | 3 | 0.017647 |
| Rosales | Cannabaceae | Cannabaceae | *Humulus_lupulus* | 170 | 3 | 0.017647 |
| Rosales | Moraceae | Moraceae | *Ficus_spp.* | 1125 | 2 | 0.001778 |
| Rosales | Moraceae | Moraceae | *Morus_spp.* | 1125 | 2 | 0.001778 |
| Rosales | Urticaceae | Urticaceae | *Pilea_spp.* | 2625 | 3 | 0.001143 |
| Rosales | Urticaceae | Urticaceae | *Boehmeria_spp.* | 2625 | 3 | 0.001143 |
| Rosales | Urticaceae | Urticaceae | *Urtica* | 2625 | 3 | 0.001143 |
| Fagales | Nothofagaceae | Nothofagaceae | *Nothofagus_antarctica* | 35 | 1 | 0.028571 |
| Fagales | Fagaceae | Fagaceae | *Fagus_spp.* | 670 | 3 | 0.004478 |
| Fagales | Fagaceae | Fagaceae | *Chrysolepis_spp.* | 670 | 3 | 0.004478 |
| Fagales | Fagaceae | Fagaceae | *Quercus_spp.* | 670 | 3 | 0.004478 |
| Fagales | Betulaceae | Betulaceae plus Ticodendraceae | *Alnus* | 146 | 1 | 0.006849 |
| Fagales | Casuarinaceae | Casuarinaceae | *Casuarina* | 95 | 1 | 0.010526 |
| Fagales | Juglandaceae | Juglandaceae | *Juglans_spp.* | 50 | 1 | 0.020000 |
| Fagales | Myricaceae | Myricaceae plus Rhoipteleaceae | *Morella_cerifera* | 58 | 2 | 0.034483 |
| Fagales | Myricaceae | Myricaceae plus Rhoipteleaceae | *Myrica* | 58 | 2 | 0.034483 |
| Cucurbitales | Anisophylleaceae | Anisophylleaceae | *Anisophyllea_fallax* | 34 | 1 | 0.029412 |
| Cucurbitales | Coriariaceae | Coriariaceae | *Coriaria_ruscifolia* | 5 | 1 | 0.200000 |
| Cucurbitales | Corynocarpaceae | Corynocarpaceae | *Corynocarpus_laevigatus* | 6 | 1 | 0.166667 |
| Cucurbitales | Tetramelaceae | Tetramelaceae | *Tetrameles_nudiflora* | 2 | 1 | 0.500000 |
| Cucurbitales | Begoniaceae | Begoniaceae | *Begonia_spp.* | 1501 | 1 | 0.000666 |
| Cucurbitales | Datiscaceae | Datiscaceae | *Datisca_spp.* | 2 | 1 | 0.500000 |
| Cucurbitales | Cucurbitaceae | Cucurbitaceae | *Xerosicyos_danguyi* | 960 | 5 | 0.005208 |
| Cucurbitales | Cucurbitaceae | Cucurbitaceae | *Dendrosicyos_socotranus* | 960 | 5 | 0.005208 |
| Cucurbitales | Cucurbitaceae | Cucurbitaceae | *Cucurbita* | 960 | 5 | 0.005208 |
| Cucurbitales | Cucurbitaceae | Cucurbitaceae | *Coccinia_sessilifolia* | 960 | 5 | 0.005208 |
| Cucurbitales | Cucurbitaceae | Cucurbitaceae | *Cucumis* | 960 | 5 | 0.005208 |
| Celastrales | Lepidobotryaceae | Lepidobotryaceae | *Lepidobotrys* | 3 | 2 | 0.666667 |
| Celastrales | Lepidobotryaceae | Lepidobotryaceae | *Ruptiliocarpon* | 3 | 2 | 0.666667 |
| Celastrales | Celastraceae (Parnassiaceae) | Celastraceae (incl. Parnassiaceae) | *Parnassia_spp.* | 1400 | 12 | 0.008571 |
| Celastrales | Celastraceae | Celastraceae (incl. Parnassiaceae) | *Siphonodon_spp.* | 1400 | 12 | 0.008571 |

| | | | | | | |
|---|---|---|---|---|---|---|
| Celastrales | Celastraceae | Celastraceae (incl. Parnassiaceae) | *Denhamia* | 1400 | 12 | 0.008571 |
| Celastrales | Celastraceae | Celastraceae (incl. Parnassiaceae) | *Stackhousia_spp.* | 1400 | 12 | 0.008571 |
| Celastrales | Celastraceae | Celastraceae (incl. Parnassiaceae) | *Maytenus* | 1400 | 12 | 0.008571 |
| Celastrales | Celastraceae | Celastraceae (incl. Parnassiaceae) | *Plagiopteron* | 1400 | 12 | 0.008571 |
| Celastrales | Celastraceae | Celastraceae (incl. Parnassiaceae) | *Brexia_madagascariensis* | 1400 | 12 | 0.008571 |
| Celastrales | Celastraceae | Celastraceae (incl. Parnassiaceae) | *Elaeodendron* | 1400 | 12 | 0.008571 |
| Celastrales | Celastraceae | Celastraceae (incl. Parnassiaceae) | *Celastrus* | 1400 | 12 | 0.008571 |
| Celastrales | Celastraceae | Celastraceae (incl. Parnassiaceae) | *Tripterygium* | 1400 | 12 | 0.008571 |
| Celastrales | Celastraceae | Celastraceae (incl. Parnassiaceae) | *Euonymus* | 1400 | 12 | 0.008571 |
| Celastrales | Celastraceae | Celastraceae (incl. Parnassiaceae) | *Paxistima* | 1400 | 12 | 0.008571 |
| Oxalidales | Huaceae | Huaceae | *Afrostyrax_lepidophyllus* | 3 | 2 | 0.666667 |
| Oxalidales | Huaceae | Huaceae | *Hua_gabonii* | 3 | 2 | 0.666667 |
| Oxalidales | Connaraceae | Connaraceae | *Connarus_spp.* | 180 | 2 | 0.011111 |
| Oxalidales | Connaraceae | Connaraceae | *Rourea_minor* | 180 | 2 | 0.011111 |
| Oxalidales | Oxalidaceae | Oxalidaceae | *Oxalis_spp.* | 770 | 3 | 0.003896 |
| Oxalidales | Oxalidaceae | Oxalidaceae | *Averrhoa_carambola* | 770 | 3 | 0.003896 |
| Oxalidales | Oxalidaceae | Oxalidaceae | *Dapania* | 770 | 3 | 0.003896 |
| Oxalidales | Brunelliaceae | Brunelliaceae | *Brunellia_acutangula* | 55 | 1 | 0.018182 |
| Oxalidales | Cephalotaceae | Cephalotaceae | *Cephalotus* | 1 | 1 | 1.000000 |
| Oxalidales | Cunoniaceae | Cunoniaceae | *Davidsonia* | 280 | 2 | 0.007143 |
| Oxalidales | Cunoniaceae | Cunoniaceae | *Eucryphia* | 280 | 2 | 0.007143 |
| Oxalidales | Elaeocarpaceae | Elaeocarpaceae | *Sloanea_spp.* | 605 | 3 | 0.004959 |
| Oxalidales | Elaeocarpaceae | Elaeocarpaceae | *Crinodendron_spp.* | 605 | 3 | 0.004959 |
| Oxalidales | Elaeocarpaceae | Elaeocarpaceae | *Elaeocarpus* | 605 | 3 | 0.004959 |
| Malpighiales | Humiriaceae | Humiriaceae | *Sacoglottis* | 50 | 3 | 0.060000 |
| Malpighiales | Humiriaceae | Humiriaceae | *Humiria_balsaminifera* | 50 | 3 | 0.060000 |
| Malpighiales | Humiriaceae | Humiriaceae | *Vantanea* | 50 | 3 | 0.060000 |
| Malpighiales | Ixonanthaceae | Ixonanthaceae | *Ochthocosmus* | 21 | 1 | 0.047619 |
| Malpighiales | Irvingiaceae | Irvingiaceae | *Irvingia* | 10 | 2 | 0.200000 |
| Malpighiales | Irvingiaceae | Irvingiaceae | *Klainedoxa* | 10 | 2 | 0.200000 |
| Malpighiales | Clusiaceae | Clusiaceae | *Symphonia_tanalensis* | 595 | 3 | 0.005042 |

| | | | | | | |
|---|---|---|---|---|---|---|
| Malpighiales | Clusiaceae | Clusiaceae | *Clusia_gundlachii* | 595 | 3 | 0.005042 |
| Malpighiales | Clusiaceae | Clusiaceae | *Garcinia* | 595 | 3 | 0.005042 |
| Malpighiales | Bonnetiaceae | Bonnetiaceae | *Archytaea* | 35 | 2 | 0.057143 |
| Malpighiales | Bonnetiaceae | Bonnetiaceae | *Bonnetia* | 35 | 2 | 0.057143 |
| Malpighiales | Calophyllaceae | Calophyllaceae | *Mesua* | 460 | 3 | 0.006522 |
| Malpighiales | Calophyllaceae | Calophyllaceae | *Calophyllum* | 460 | 3 | 0.006522 |
| Malpighiales | Calophyllaceae | Calophyllaceae | *Mammea* | 460 | 3 | 0.006522 |
| Malpighiales | Podostemaceae | Podostemaceae | *Marathrum_spp.* | 270 | 2 | 0.007407 |
| Malpighiales | Podostemaceae | Podostemaceae | *Podostemum_ceratophyllum* | 270 | 2 | 0.007407 |
| Malpighiales | Hypericaceae | Hypericaceae | *Cratoxylum_cochinchinense* | 560 | 4 | 0.007143 |
| Malpighiales | Hypericaceae | Hypericaceae | *Eliea_articulata* | 560 | 4 | 0.007143 |
| Malpighiales | Hypericaceae | Hypericaceae | *Hypericum_perforatum* | 560 | 4 | 0.007143 |
| Malpighiales | Hypericaceae | Hypericaceae | *Vismia_spp.* | 560 | 4 | 0.007143 |
| Malpighiales | Achariaceae | Achariaceae | *Erythrospermum* | 145 | 5 | 0.034483 |
| Malpighiales | Achariaceae | Achariaceae | *Hydnocarpus_spp.* | 145 | 5 | 0.034483 |
| Malpighiales | Achariaceae | Achariaceae | *Pangium* | 145 | 5 | 0.034483 |
| Malpighiales | Achariaceae | Achariaceae | *Acharia_tragodes* | 145 | 5 | 0.034483 |
| Malpighiales | Achariaceae | Achariaceae | *Kiggelaria* | 145 | 5 | 0.034483 |
| Malpighiales | Passifloraceae | Passifloraceae | *Malesherbia_linearifolia* | 935 | 4 | 0.004278 |
| Malpighiales | Passifloraceae | Passifloraceae | *Turnera_spp.* | 935 | 4 | 0.004278 |
| Malpighiales | Passifloraceae | Passifloraceae | *Paropsia* | 935 | 4 | 0.004278 |
| Malpighiales | Passifloraceae | Passifloraceae | *Passiflora_spp.* | 935 | 4 | 0.004278 |
| Malpighiales | Goupiaceae | Goupiaceae | *Goupia* | 2 | 1 | 0.500000 |
| Malpighiales | Violaceae | Violaceae | *Rinorea_pubiflora* | 800 | 5 | 0.006250 |
| Malpighiales | Violaceae | Violaceae | *Viola_spp.* | 800 | 5 | 0.006250 |
| Malpighiales | Violaceae | Violaceae | *Hymenanthera* | 800 | 5 | 0.006250 |
| Malpighiales | Violaceae | Violaceae | *Hybanthus* | 800 | 5 | 0.006250 |
| Malpighiales | Violaceae | Violaceae | *Leonia* | 800 | 5 | 0.006250 |
| Malpighiales | Lacistemataceae | Lacistemataceae | *Lacistema_aggregatum* | 14 | 2 | 0.142857 |
| Malpighiales | Lacistemataceae | Lacistemataceae | *Lozania* | 14 | 2 | 0.142857 |
| Malpighiales | Salicaceae | Salicaceae | *Casearia_spp.* | 1010 | 11 | 0.010891 |
| Malpighiales | Salicaceae | Salicaceae | *Lunania* | 1010 | 11 | 0.010891 |
| Malpighiales | Salicaceae | Salicaceae | *Scyphostegia* | 1010 | 11 | 0.010891 |
| Malpighiales | Salicaceae | Salicaceae | *Abatia* | 1010 | 11 | 0.010891 |
| Malpighiales | Salicaceae | Salicaceae | *Prockia* | 1010 | 11 | 0.010891 |
| Malpighiales | Salicaceae | Salicaceae | *Dovyalis* | 1010 | 11 | 0.010891 |
| Malpighiales | Salicaceae | Salicaceae | *Flacourtia* | 1010 | 11 | 0.010891 |
| Malpighiales | Salicaceae | Salicaceae | *Poliothyrsis* | 1010 | 11 | 0.010891 |
| Malpighiales | Salicaceae | Salicaceae | *Idesia* | 1010 | 11 | 0.010891 |
| Malpighiales | Salicaceae | Salicaceae | *Populus_spp.* | 1010 | 11 | 0.010891 |
| Malpighiales | Salicaceae | Salicaceae | *Salix* | 1010 | 11 | 0.010891 |
| Malpighiales | Elatinaceae | Elatinaceae | *Bergia* | 35 | 2 | 0.057143 |
| Malpighiales | Elatinaceae | Elatinaceae | *Elatine* | 35 | 2 | 0.057143 |
| Malpighiales | Malpighiaceae | Malpighiaceae | *Byrsonima_crassifolia* | 1250 | 6 | 0.004800 |
| Malpighiales | Malpighiaceae | Malpighiaceae | *Acridocarpus_natalitius* | 1250 | 6 | 0.004800 |
| Malpighiales | Malpighiaceae | Malpighiaceae | *Tetrapterys_glabrifolia* | 1250 | 6 | 0.004800 |
| Malpighiales | Malpighiaceae | Malpighiaceae | *Thryallis* | 1250 | 6 | 0.004800 |

| | | | | | | |
|---|---|---|---|---|---|---|
| Malpighiales | Malpighiaceae | Malpighiaceae | *Dicella* | 1250 | 6 | 0.004800 |
| Malpighiales | Malpighiaceae | Malpighiaceae | *Malpighia* | 1250 | 6 | 0.004800 |
| Malpighiales | Phyllanthaceae | Phyllanthaceae | *Aporusa* | 1745 | 6 | 0.003438 |
| Malpighiales | Phyllanthaceae | Phyllanthaceae | *Bischofia_javanica* | 1745 | 6 | 0.003438 |
| Malpighiales | Phyllanthaceae | Phyllanthaceae | *Phyllanthus_spp.* | 1745 | 6 | 0.003438 |
| Malpighiales | Phyllanthaceae | Phyllanthaceae | *Heywoodia* | 1745 | 6 | 0.003438 |
| Malpighiales | Phyllanthaceae | Phyllanthaceae | *Croizatia* | 1745 | 6 | 0.003438 |
| Malpighiales | Phyllanthaceae | Phyllanthaceae | *Lachnostylis* | 1745 | 6 | 0.003438 |
| Malpighiales | Picrodendraceae | Picrodendraceae | *Podocalyx* | 80 | 7 | 0.087500 |
| Malpighiales | Picrodendraceae | Picrodendraceae | *Androstachys_johnsonii* | 80 | 7 | 0.087500 |
| Malpighiales | Picrodendraceae | Picrodendraceae | *Tetracoccus* | 80 | 7 | 0.087500 |
| Malpighiales | Picrodendraceae | Picrodendraceae | *Petalostigma* | 80 | 7 | 0.087500 |
| Malpighiales | Picrodendraceae | Picrodendraceae | *Dissiliaria* | 80 | 7 | 0.087500 |
| Malpighiales | Picrodendraceae | Picrodendraceae | *Austrobuxus* | 80 | 7 | 0.087500 |
| Malpighiales | Picrodendraceae | Picrodendraceae | *Micrantheum* | 80 | 7 | 0.087500 |
| Malpighiales | Putranjivaceae | Putranjivaceae | *Drypetes* | 210 | 1 | 0.004762 |
| Malpighiales | Lophopyxidaceae | Lophopyxidaceae | *Lophopyxis* | 1 | 1 | 1.000000 |
| Malpighiales | Balanopaceae | Balanopaceae | *Balanops_vieillardi* | 9 | 1 | 0.111111 |
| Malpighiales | Trigoniaceae | Trigoniaceae | *Trigonia* | 28 | 1 | 0.035714 |
| Malpighiales | Dichapetalaceae | Dichapetalaceae | *Dichapetalum* | 165 | 2 | 0.012121 |
| Malpighiales | Dichapetalaceae | Dichapetalaceae | *Tapura* | 165 | 2 | 0.012121 |
| Malpighiales | Euphroniaceae | Euphroniaceae | *Euphronia* | 3 | 1 | 0.333333 |
| Malpighiales | Chrysobalanaceae | Chrysobalanaceae | *Atuna* | 460 | 4 | 0.008696 |
| Malpighiales | Chrysobalanaceae | Chrysobalanaceae | *Chrysobalanus_icaco* | 460 | 4 | 0.008696 |
| Malpighiales | Chrysobalanaceae | Chrysobalanaceae | *Hirtella* | 460 | 4 | 0.008696 |
| Malpighiales | Chrysobalanaceae | Chrysobalanaceae | *Licania_spp.* | 460 | 4 | 0.008696 |
| Malpighiales | Pandaceae | Pandaceae | *Microdesmis* | 15 | 3 | 0.200000 |
| Malpighiales | Pandaceae | Pandaceae | *Galearia* | 15 | 3 | 0.200000 |
| Malpighiales | Pandaceae | Pandaceae | *Panda* | 15 | 3 | 0.200000 |
| Malpighiales | Centroplacaceae | Centroplacaceae | *Bhesa_paniculata* | 6 | 2 | 0.333333 |
| Malpighiales | Centroplacaceae | Centroplacaceae | *Centroplacus* | 6 | 2 | 0.333333 |
| Malpighiales | Linaceae | Linaceae | *Reinwardtia_indica* | 300 | 4 | 0.013333 |
| Malpighiales | Linaceae | Linaceae | *Linum* | 300 | 4 | 0.013333 |
| Malpighiales | Linaceae | Linaceae | *Durandea* | 300 | 4 | 0.013333 |
| Malpighiales | Linaceae | Linaceae | *Hugonia* | 300 | 4 | 0.013333 |
| Malpighiales | Ctenolophonaceae | Ctenolophonaceae | *Ctenolophon* | 3 | 1 | 0.333333 |
| Malpighiales | Erythroxylaceae | Erythroxylaceae | *Aneulophus_africanus* | 240 | 2 | 0.008333 |
| Malpighiales | Erythroxylaceae | Erythroxylaceae | *Erythroxylum_spp.* | 240 | 2 | 0.008333 |
| Malpighiales | Rhizophoraceae | Rhizophoraceae | *Cassipourea_lanceolata* | 149 | 6 | 0.040268 |
| Malpighiales | Rhizophoraceae | Rhizophoraceae | *Paradrypetes* | 149 | 6 | 0.040268 |
| Malpighiales | Rhizophoraceae | Rhizophoraceae | *Crossostylis_grandiflora* | 149 | 6 | 0.040268 |
| Malpighiales | Rhizophoraceae | Rhizophoraceae | *Rhizophora_spp.* | 149 | 6 | 0.040268 |
| Malpighiales | Rhizophoraceae | Rhizophoraceae | *Bruguiera* | 149 | 6 | 0.040268 |
| Malpighiales | Rhizophoraceae | Rhizophoraceae | *Carallia* | 149 | 6 | 0.040268 |

| | | | | | | |
|---|---|---|---|---|---|---|
| Malpighiales | Caryocaraceae | Caryocaraceae | *Caryocar* | 21 | 1 | 0.047619 |
| Malpighiales | Ochnaceae | Ochnaceae | *Medusagyne_oppositifolia* | 495 | 7 | 0.014141 |
| Malpighiales | Ochnaceae | Ochnaceae | *Quiina* | 495 | 7 | 0.014141 |
| Malpighiales | Ochnaceae | Ochnaceae | *Touroulia* | 495 | 7 | 0.014141 |
| Malpighiales | Ochnaceae | Ochnaceae | *Luxemburgia* | 495 | 7 | 0.014141 |
| Malpighiales | Ochnaceae | Ochnaceae | *Ochna* | 495 | 7 | 0.014141 |
| Malpighiales | Ochnaceae | Ochnaceae | *Cespedesia* | 495 | 7 | 0.014141 |
| Malpighiales | Ochnaceae | Ochnaceae | *Sauvagesia_spp.* | 495 | 7 | 0.014141 |
| Malpighiales | Peraceae | Peraceae | *Pogonophora_schomburgkiana* | 135 | 3 | 0.022222 |
| Malpighiales | Peraceae | Peraceae | *Clutia* | 135 | 3 | 0.022222 |
| Malpighiales | Peraceae | Peraceae | *Pera_bicolor* | 135 | 3 | 0.022222 |
| Malpighiales | Euphorbiaceae | Euphorbiaceae plus Rafflesiaceae | *Neoscortechinia_kingii* | 5755 | 21 | 0.003649 |
| Malpighiales | Euphorbiaceae | Euphorbiaceae plus Rafflesiaceae | *Endospermum* | 5755 | 21 | 0.003649 |
| Malpighiales | Euphorbiaceae | Euphorbiaceae plus Rafflesiaceae | *Omphalea* | 5755 | 21 | 0.003649 |
| Malpighiales | Euphorbiaceae | Euphorbiaceae plus Rafflesiaceae | *Tetrorchidium* | 5755 | 21 | 0.003649 |
| Malpighiales | Euphorbiaceae | Euphorbiaceae plus Rafflesiaceae | *Suregada* | 5755 | 21 | 0.003649 |
| Malpighiales | Euphorbiaceae | Euphorbiaceae plus Rafflesiaceae | *Moultonianthus* | 5755 | 21 | 0.003649 |
| Malpighiales | Euphorbiaceae | Euphorbiaceae plus Rafflesiaceae | *Hevea* | 5755 | 21 | 0.003649 |
| Malpighiales | Euphorbiaceae | Euphorbiaceae plus Rafflesiaceae | *Manihot_esculenta* | 5755 | 21 | 0.003649 |
| Malpighiales | Euphorbiaceae | Euphorbiaceae plus Rafflesiaceae | *Croton* | 5755 | 21 | 0.003649 |
| Malpighiales | Euphorbiaceae | Euphorbiaceae plus Rafflesiaceae | *Codiaeum* | 5755 | 21 | 0.003649 |
| Malpighiales | Euphorbiaceae | Euphorbiaceae plus Rafflesiaceae | *Trigonostemon* | 5755 | 21 | 0.003649 |
| Malpighiales | Euphorbiaceae | Euphorbiaceae plus Rafflesiaceae | *Pimelodendron* | 5755 | 21 | 0.003649 |
| Malpighiales | Euphorbiaceae | Euphorbiaceae plus Rafflesiaceae | *Euphorbia_spp.* | 5755 | 21 | 0.003649 |

| | | | | | | |
|---|---|---|---|---|---|---|
| Malpighiales | Euphorbiaceae | Euphorbiaceae plus Rafflesiaceae | *Homalanthus* | 5755 | 21 | 0.003649 |
| Malpighiales | Euphorbiaceae | Euphorbiaceae plus Rafflesiaceae | *Hura* | 5755 | 21 | 0.003649 |
| Malpighiales | Euphorbiaceae | Euphorbiaceae plus Rafflesiaceae | *Conceveiba* | 5755 | 21 | 0.003649 |
| Malpighiales | Euphorbiaceae | Euphorbiaceae plus Rafflesiaceae | *Dalechampia* | 5755 | 21 | 0.003649 |
| Malpighiales | Euphorbiaceae | Euphorbiaceae plus Rafflesiaceae | *Lasiocroton* | 5755 | 21 | 0.003649 |
| Malpighiales | Euphorbiaceae | Euphorbiaceae plus Rafflesiaceae | *Ricinus* | 5755 | 21 | 0.003649 |
| Malpighiales | Euphorbiaceae | Euphorbiaceae plus Rafflesiaceae | *Acalypha_californica* | 5755 | 21 | 0.003649 |
| Malpighiales | Euphorbiaceae | Euphorbiaceae plus Rafflesiaceae | *Spathiostemon* | 5755 | 21 | 0.003649 |

**Supplementary Table 4.** Thirty core angiosperm rate shifts, including shifts in adjacent branches in a distinct phylogenetic region. Core shifts correspond to those within 95% of the magnitude of the highest Marginal Odds Ratio (MOR) in each analysis with different prior for the expected number of shifts (i.e., 0.1, 1, 5, 10, 50, 100); and that are found in at least four of the six analyses. Core shifts detected with high probability on a single phylogenetic branch, and with moderate probability on adjacent branches on a distict phylogenetic region, are indicated. The latter are indicated by the number of the shift, followed by a letter that indicates one of the adjacent branches. The Marginal Odds Ratio (MOR), time, speciation and extinction rates of each shift, estimated with prior for the expected number of shifts = 100, are indicated.

| Core Shift Number | Core Shift Name | Branches (MOR) | Clade Content | Number of Species | Mean Time of shift (min-max) Ma | Mean Speciation Rate (min-max) | Mean Extinction Rate (min-max) |
|---|---|---|---|---|---|---|---|
| 1* | ca. Mesangiospermae | 1a (88.76) | Nymphaeales, Austrobaileyales, Mesangiospermae | 276765 | 139.17 (138.97-139.40) | 0.0989 (0.0910-0.1086) | 0.0275 (0.0189-0.0382) |
| | | 1b (38.68) | Mesangiospermae | 276601 | 136.88 (135.92-137.68) | 0.0998 (0.0918-0.1098) | 0.0278 (0.0188-0.0386) |
| 2 | Vitales+Rosids | 2 (11.94) | Vitales, Rosidae | 81498 | 121.98 (121.32-122.40) | 0.1087 (0.0942-0.1279) | 0.0305 (0.0152-0.0516) |
| 3 | ca. Fabidae | 3a (132.07) | Fabidae | 51100 | 117.68 (116.81-118.58) | 0.1123 (0.0957-0.1351) | 0.0312 (0.0135-0.0551) |
| | | 3b (47.93) | Fabidae minus Zygophyllales | 50797 | 116.28 (115.79-116.81) | 0.1131 (0.0964-0.1361) | 0.0316 (0.0135-0.0560) |
| | | 3c (23.54) | Celastrales, Oxalidales, Malpighiales | 19309 | 113.80 (112.04-115.78) | 0.1041 (0.0894-0.1254) | 0.0221 (0.0074-0.0469) |
| | | 3d (121.01) | Oxalidales, Malpighiales | 17906 | 111.70 (111.43-112.02) | 0.0985 (0.0848-0.1214) | 0.0181 (0.0040-0.0429) |
| 4 | Ericales | 4 (47.84) | Ericales | 11871 | 107.76 (103.59-112.34) | 0.0976 (0.0753-0.1457) | 0.0244 (0.0026-0.0807) |
| 5 | Myrtales | 5 (23.42) | Myrtales | 11632 | 105.45 (96.64-116.37) | 0.1435 (0.0818-0.2837) | 0.0586 (0.0011-0.2147) |
| 6* | Arecales+Commelinales+Zingiberales | 6 (17.19) | Arecales, Commelinales, Zingiberales | 5450 | 102.14 (98.21-106.73) | 0.1174 (0.0702-0.1939) | 0.0385 (0.0018-0.1204) |
| 7 | ca. Ranunculaceae | 7a (47.86) | Menispermaceae, Berberidaceae, Ranunculaceae | 3668 | 93.78 (89.93-98.17) | 0.1328 (0.0837-0.2644) | 0.0484 (0.0013-0.1948) |
| | | 7b (8.60) | Berberidaceae, Ranunculaceae | 3226 | 84.70 (80.29-89.90) | 0.1411 (0.0869-0.2847) | 0.0528 (0.0012-0.2135) |

| | | | | | | | |
|---|---|---|---|---|---|---|---|
| 8* | Lamiidae | 9 (133.2) | Gentianales, Solanales, Boraginales, Lamiales | 50437 | 91.14 (89.75-92.67) | 0.1427 (0.1269-0.1663) | 0.0201 (0.0069-0.0477) |
| 9* | ca. Fabaceae | 9a (12.94) | Surianaceae, Fabaceae | 19508 | 93.83 (92.16-96.03) | 0.2565 (0.1247-0.4681) | 0.1533 (0.0111-0.3789) |
| | | 9b (47.43) | Fabaceae | 19500 | 88.05 (84.76-92.13) | 0.2780 (0.1316-0.5094) | 0.1675 (0.0104-0.4132) |
| 10 | Asparagaceae+ | 11 (57.19) | Tecophilaceae, Iridaceae, Asphodelaceae, Xanthorrhoeaceae, Amarillidaceae, Asparagaceae | 4227 | 84.25 (80.66-88.94) | 0.1638 (0.1083-0.3145) | 0.0540 (0.0012-0.2181) |
| 11 | ca. Sapindales | 11a (12.14) | Sapindales minus Biebersteiniaceae | 6077 | 85.35 (83.42-87.50) | 0.1251 (0.0755-0.2373) | 0.0391 (0.0009-0.1620) |
| | | 11b (110.93) | Sapindales minus Biebersteiniaceae and Nitrariaceae | 6061 | 81.50 (79.92-83.35) | 0.1355 (0.0760-0.2622) | 0.0434 (0.0006-0.1821) |
| 12 | Polygonaceae+ Polygonaceae | 12 (14.40) | Polygonaceae, Plumbaginaceae | 1946 | 76.87 (67.91-93.26) | 0.1566 (0.0412-0.3736) | 0.0735 (0.0001-0.2850) |
| 13* | ca. Asteraceae | 13a (63.71) | Stylidaceae, Menyanthaceae, Goodeniaceae, Calyceraceae, Asteraceae | 24393 | 76.79 (76.48-77.10) | 0.2258 (0.1329-0.4081) | 0.1027 (0.0059-0.2936) |
| | | 13b (30.30) | Goodeniaceae, Calyceraceae, Asteraceae | 24090 | 61.14 (57.05-68.26) | 0.3394 (0.1914-0.6527) | 0.1643 (0.0086-0.4950) |
| | | 13c (21.05) | Asteraceae | 23600 | 47.07 (45.27-49.27) | 0.3770 (0.2075-0.7203) | 0.1788 (0.0079-0.5359) |
| 14 | Dipsacales | 14 (32.36) | Dipsacales | 1090 | 75.76 (70.94-81.84) | 0.1690 (0.1078-0.2938) | 0.0685 (0.0039-0.2108) |
| 15* | Montiniaceae+ Hydroleaceae+ Sphenocleaceae | 15 (61.74) | Montiniaceae, Hydroleaceae, Sphenocleaceae | 19 | 75.76 (72.04-79.24) | 0.0705 (0.0190-0.2240) | 0.0461 (0.0010-0.2185) |
| 16 | Orchidaceae | 16 (9.96) | Orchidaceae | 22075 | 73.07 (59.75-108.78) | 0.2349 (0.1059-0.5024) | 0.1225 (0.0042-0.3954) |
| 17 | Crassulaceae | 17 (11.74) | Crassulaceae | 1400 | 72.91 (60.69-95.33) | 0.1713 (0.0415-0.3842) | 0.0861 (0.0002-0.2998) |
| 18 | Moraceae+Urticaceae | 18 (12.64) | Moraceae, Urticaceae | 1125 | 70.93 (68.52-73.43) | 0.1199 (0.0728-0.2930) | 0.0333 (0.0001-0.1998) |

| | | | | | | | |
|---|---|---|---|---|---|---|---|
| 19* | ca. Apiales | 19a (61.66) | Pittosporaceae. Araliaceae, Myodocarpaceae, Apiaceae | 5449 | 66.28 (63.37-70.53) | 0.2895 (0.1584-0.5645) | 0.1584 (0.0151-0.4485) |
| | | 19b (7.54) | Araliaceae, Myodocarpaceae, Apiaceae | 5249 | 61.49 (60.20-63.37) | 0.2949 (0.1626-0.5716) | 0.1609 (0.0153-0.4540) |
| 20 | ca. Cyperaceae | 20a (10.38) | Juncaceae, Cyperaceae | 5864 | 62.74 (55.19-77.05) | 0.1760 (0.0691-0.3829) | 0.0671 (0.0009-0.2713) |
| | | 20b (9.82) | Cyperaceae | 5430 | 37.29 (29.93-55.14) | 0.2310 (0.0633-0.4817) | 0.0907 (0.0009-0.3505) |
| 21 | ca. Euphorbiaceae | 21a (21.68) | Euphorbiaceae | 5755 | 67.13 (61.91-74.23) | 0.2125 (0.1399-0.3738) | 0.0819 (0.0034-0.2613) |
| | | 21b (43.88) | Euphorbiaceae minus Neoscortechinia | 5750 | 59.34 (57.09-61.88) | 0.2190 (0.1423-0.3907) | 0.0854 (0.0030-0.2720) |
| 22 | Campanulaceae | 22 (16.54) | Campanulaceae | 2380 | 54.34 (45.59-75.98) | 0.2348 (0.1284-0.4606) | 0.1059 (0.0036-0.3586) |
| 23 | Celastraceae | 23 (16.95) | Celastraceae excluding Parnassia | 1335 | 51.15 (42.83-68.40) | 0.2474 (0.1365-0.4677) | 0.1155 (0.0036-0.3642) |
| 24 | Amaranthaceae | 24 (16.53) | Amaranthaceae | 1379 | 51.01 (43.67-64.07) | 0.2065 (0.1077-0.4156) | 0.0796 (0.0020-0.3031) |
| 25* | Piperaceae | 25 (17.50) | Piperaceae | 3615 | 48.79 (39.37-65.47) | 0.1988 (0.0403-0.4062) | 0.0709 (0.0003-0.2782) |
| 26* | ca. Brassicaceae+Cleomaceae+Capparaceae | 26a (31.40) | Capparaceae, Brassicaceae, Cleomaceae | 4490 | 47.96 (44.22-54.18) | 0.2469 (0.1425-0.4518) | 0.0819 (0.0023-0.2986) |
| | | 26b (10.40) | Brassicaceae, Cleomaceae | 4010 | 32.79 (27.45-44.01) | 0.2888 (0.1596-0.5419) | 0.0963 (0.0022-0.3489) |
| 27 | Poaceae | 27 (26.03) | Poaceae | 11337 | 45.27 (39.75-58.45) | 0.3029 (0.1900-0.5542) | 0.1058 (0.0031-0.3676) |
| 28 | Cucurbitaceae | 28 (15.19) | Cucurbitaceae | 960 | 42.61 (35.54-57.08) | 0.2418 (0.0800-0.4747) | 0.0925 (0.0007-0.3326) |
| 29 | ca. Malvaceae | 29a (11.25) | Malvales excluding Neuradaceae | 6083 | 72.70 (70.14-76.17) | 0.1763 (0.1042-0.3372) | 0.0646 (0.0026-0.2361) |
| | | 29b (14.75) | Malvaceae | 4225 | 39.53 (33.31-59.69) | 0.2944 (0.1611-0.5645) | 0.1149 (0.0033-0.4014) |
| 30* | ca. Cactaceae | 30a (17.92) | Talinaceae, Portulacaceae, Cactaceae | 2025 | 35.90 (33.33-39.44) | 0.1904 (0.0976-0.3605) | 0.0570 (0.0013-0.2281) |

| | | | | | |
|---|---|---|---|---|---|
| 30b (21.47) | Portulacaceae, Cactaceae | 276765 | 30.80 (28.82-33.29) | 0.2108 (0.0974-0.4308) | 0.0629 (0.0013-0.2426) |

**Supplementary Table 5.** Values of Marginal Odds Ratio (MOR) and associated parameters. Absolute magnitude of the maximal Marginal Odds Ratio (MOR) in in six analyses conducted under different magnitudes for the prior on the expected number of shifts (0.1, 1, 5, 10, 50, 100), and associated parameters.

| | 0.1 | 1 | 5 | 10 | 50 | 100 |
|---|---|---|---|---|---|---|
| Magnitude of highest MOR | 254,012.78 | 16,281.31 | 3288.44 | 2049.28 | 292.89 | 133.28 |
| Magnitude of 95% cutoff | 12,700.64 | 814.07 | 164.42 | 102.46 | 14.64 | 6.66 |
| Number of shifts within the cutoff | 22 | 44 | 47 | 41 | 59 | 69 |
| Number of shifts within the cutoff only present in this analysis | 2 | 2 | 3 | 1 | 17 | 24 |
| Number of shifts with MOR >0 | 611 | 1191 | 1344 | 1386 | 1400 | 1386 |
